# First Broad-spectrum Sámi and Norse diets in northern Norway over the last 4000 years revealed by sedaDNA

**DOI:** 10.64898/2026.11.20.765463

**Authors:** Aya Komatsu, Stephen Wickler, Inger G. Alsos, Lucas Elliott, Helge Høeg, Torbjørn Alm, Antony Brown

**Affiliations:** The Arctic University Museum of Norway, UiT – The Arctic University of Norway, Tromsø, Norway; Natural History Museum, University of Oslo

## Abstract

The comprehensive specification of palaeo-diets has depended upon the preservation of plants and bones in archaeological deposits, which is limited by preservation conditions. Although middens at settlement sites in northern Norway can have comprehensive faunal archives they still may not reflect the full diet. Here, we apply sedaDNA metabarcoding and shotgun sequencing to two midden complexes in Finnmark, Norway. At both sites, this doubled the species identified. At the Baŋkgohppi “Gressbakken” house site (c.4.5-3.8 BP), items were added to all food categories (Figure below) including mammals such as reindeer, beaver, and bowhead whale, suggesting deep-sea fishing practice. At the Finnes coastal Sámi Early Metal Age site (c.07-0.8 BP) and later Norse Medieval re-occupation, sedaDNA added terrestrial mammal (reindeer) and domesticated stock (goat, cattle) to the diet. The combination of traditional data with sedaDNA data provides us for the first time with a broad spectrum, suggesting seasonally balanced diets and fishing activity in northern Norway over the last 4000 years.

**Significance:** This paper is the first to utilize sedaDNA matebarcoding and shotgun sequencing for the estimation of past human diet and nutrition from archaeological sites resulting in a doubling of dietary breadth and balanced nutrition. It is also the first analysis of sedaDNA for archaeological sites in subarctic Europe. Despite the sites investigated having comprehensive assemblages of mammal and fish bones, plant remains, and microfossils (pollen), the sedaDNA doubles the number of food groups exploited in both later prehistory and the Medieval Period Finnmark. The data also reveals deep-sea fishing c. 2456-1896 BCE due to the presence of bowhead whale DNA at Baŋkgohppi. This study represents a step-change in progress towards comprehensive dietary reconstruction for past communities including those under environmental stress.

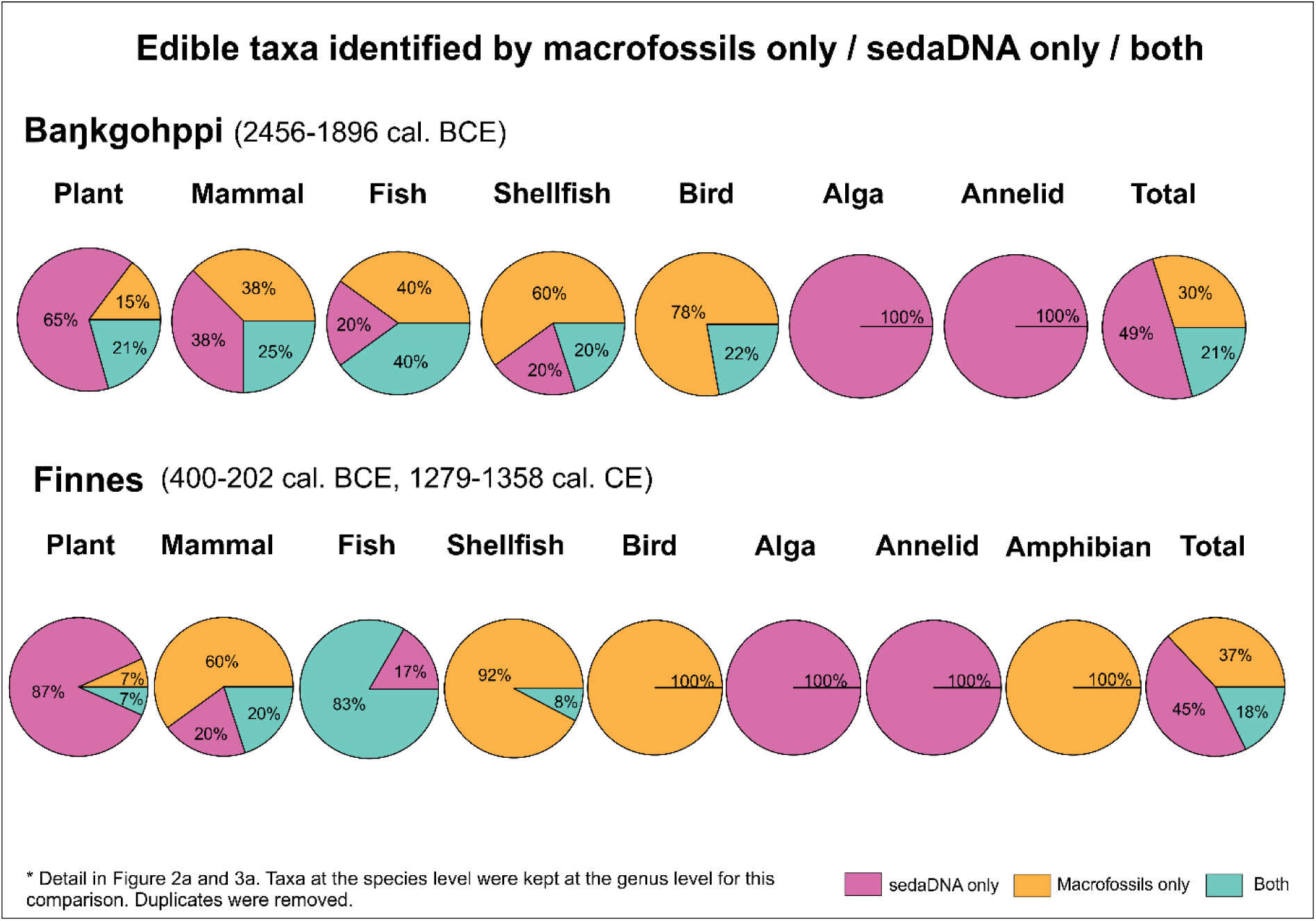

## Introduction

The dietary history of Northern Peoples reveals their sensitivity and resilience to changing environmental conditions – which is highly relevant in the face of the rapidly changing Arctic climate we are experiencing today (Arctic Council 2021; Bogdanova et al. 2021; Pearce et al. 2010). Of particular value is the determination of dietary breadth and the dietary space for adaptation to changing resource distributions (Brown et al. 2022a). The reconstruction of the past diet of humans has been undertaken for decades primarily using bone assemblages and plant macrofossils (particularly charred seeds). More recently ancient DNA (aDNA) analyses have been applied to paleofeces (Maixner et al. 2021), dental calculus (Sawafuji et al. 2020), and herbivore middens (Murray et al. 2012). The analysis of sedimentary ancient DNA (sedaDNA) using shotgun sequencing and metabarcoding has been undertaken at both Inuit and Viking archaeological sites in Greenland (Seersholm et al. 2016, 2022) but not in northern Eurasia. As DNA tends to be well-preserved in colder climates (Sirois and Buckley 2019), it should be suitable for the estimation of past biodiversity and past human diet in northern Norway using both metabarcoding and shotgun sequencing as illustrated from lake sediments (Brown et al. 2022b; Rijal et al. 2021). In this paper, sediment samples from two well-stratified archaeological sites in northern Norway are analyzed using sedaDNA metabarcoding and shotgun sequencing (Figure 1). The results of the analyses are compared with dietary data from macro and micro remains from the excavations, with the aim of providing a more comprehensive diet analysis than previously possible and new evidence of food procurement activities.

**Figure 1.**
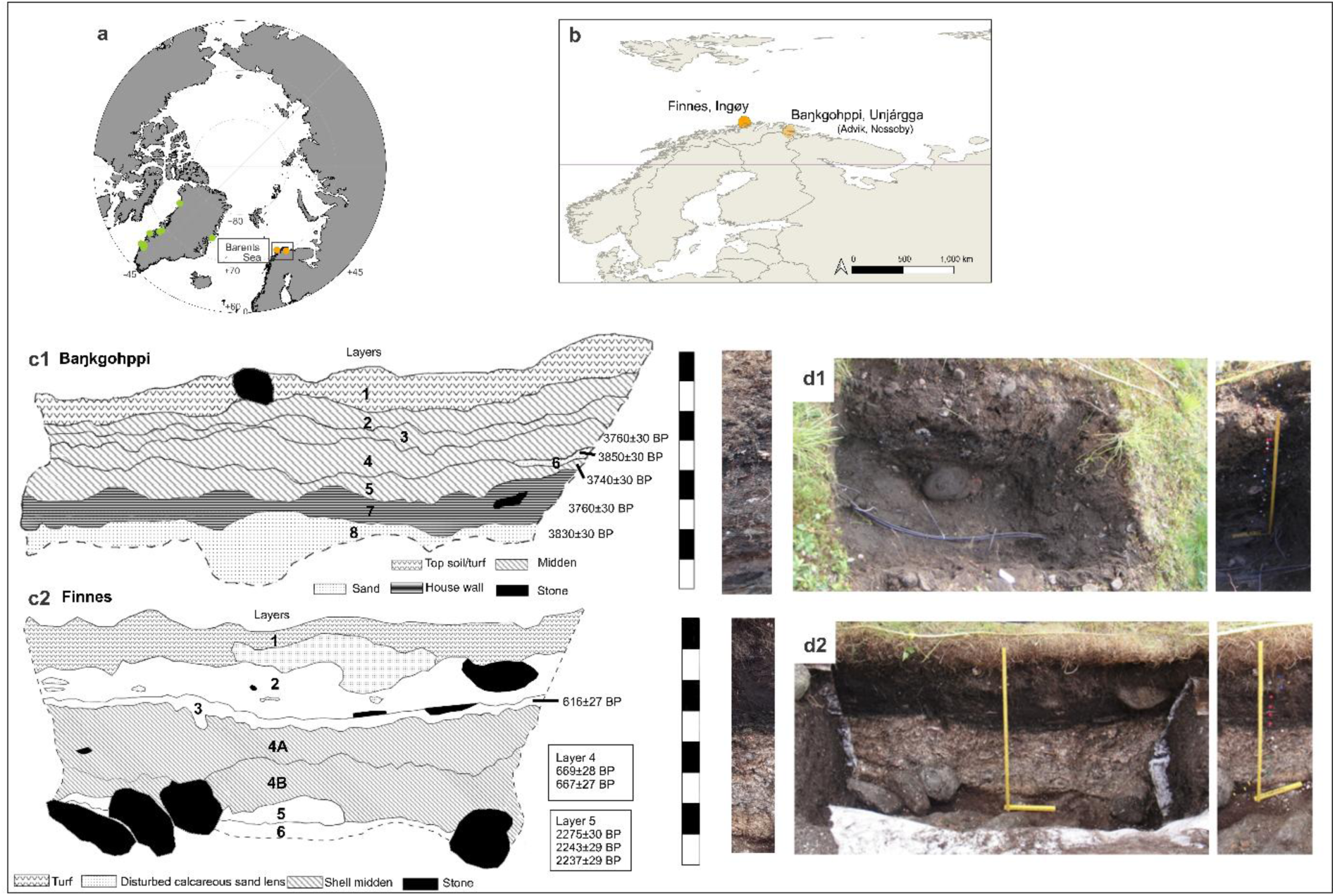
Arctic map, sites, and stratigraphy. a. Arctic map with archaeological middens and sites where sedaDNA analyses have been performed (green) and sites included in this study (orange). 1b. Location of each site included in this study. 1c and 1d. Stratigraphy and photographs of the excavated sites.

For the reconstruction of past diets, middens have been targeted as they are food-waste mounds, and the preservation of both animal/fish bones and plant remains can be expected. However, not all plant and animal remains are preserved, biassing assemblages to the most robust mammal bones and seeds, whereas sedaDNA can be preserved if it is bound to clay and sediment particles during the process of decomposition (Capo et al. 2021; Giguet-Covex et al. 2014; Pedersen et al. 2015; Taberlet et al. 2018). With the advancement of next-generation sequencing and improvements in sedaDNA extraction techniques, sedaDNA studies have allowed high-throughput sequencing at lower costs permitting application to an increasingly wide range of environments and sample types (Alsos et al. 2024; Brown et al. 2022a; Capo et al. 2021).

Poor preservation of sedaDNA remains a problem as temperature, humidity, salinity, and acidity of the environment all influence preservation (Dabney et al. 2013). The correlation between temperature and DNA degradation (Sirois and Buckley 2019) and low rates of organic decomposition make northern Norway (0.5°C mat) well-suited for sedaDNA. Other potential limitations are contamination and the differential abundance of some species. Contamination of modern DNA during sampling and laboratory work as well as mixing from upper stratigraphic layers can potentially affect the sedaDNA profile; hence, precautions are taken (see methods) and interpretation requires caution (Haile et al. 2007). The abundance or high biomass of a few species (e.g. whales) could hide the species with a low abundance of DNA (Pedersen et al. 2013), so, the combination of sedaDNA with other proxies such as faunal and floral remains is valuable, allowing cross-validation but also revealing a higher diversity of species than the individual methods alone (Parducci et al. 2019; Pedersen et al. 2015). Therefore, combining these methods should facilitate a more comprehensive understanding of past diets in northern Norway.

### Archaeological sites

Two sites in Finnmark, northern Norway were included in this study (Figure 1). The sites are Baŋkgohppi, near Nesseby excavated originally in 2013 CE (Martens et al. 2017) and Finnes, Ingøy originally excavated in 2011 CE (Wickler 2016, 2026). The samples for sedaDNA analyses were collected from the sections re-excavated for this project. These sections were exposed, checked against the original stratigraphy, and then excavated, taken back to provide fresh material for sampling. Baŋkgohppi is located in Nesseby, Finnmark (Lat. 70.076129, Long. 28.787790) on a beach terrace on Varanger Fjord at an altitude of 12-14m asl and ca. 250m from the present coastline. The site consists of multiple gressbakken-type houses, mounds, and middens, dating back to 2456-1896 cal. BCE (Martens et al. 2017; Calibration: Bronk Ramsey 2009, 2021; Reimer et al. 2020; a plan in S1). Gressbakken-type houses are named after the common house type found at the Gressbakken site, characterized by rectangular symmetrical houses with an entrance in the middle of the front long wall as well as on the back side of the wall and a fireplace (Schanche 1989). One house was partially excavated by the *InSituFarms* project, and a 0.7×3 m trench was opened, drawn, sampled, and had instruments installed in order to monitor environmental conditions (Martens et al. 2017). Due to high alkalinity of the underlying calcareous sand and low conductivity, excellent preservation conditions were observed, and a wide range of faunal and floral remains reported (S2, Martens et al. 2017). The excavation area included midden and the wall of the house, and the sediments consisted high botanical and zoological contents due to the high lime content of the midden (see the stratigraphy in Figure 1 and the description and chronology in S3; Martens et al. 2017). The vegetation of the site today is open birch (*Betula*) woodland with some pine (*Pinus*) inland. An impressive set of hunting pit traps is also located on the highest beachline above the site, suggesting that the location had been used to drive wild reindeer towards the coast.

The Finnes site (Lat. 71.07306, Long. 23.99038) is a multiphase marine exploitation and settlement site located along the northwestern shoreline of Ingøy Island on the outermost coastline of western Finnmark. The island is 18.6km^2^ in area and has beaches on the coast and inland marshes with the highest point being only 150m asl (Andreassen 2003). Surrounded by three ocean current systems; the North Atlantic Current, the Arctic Current, and the Norwegian Coastal Current (Markonic 2017), and having many islets around the island to provide shelter from the open ocean, the site lies in close proximity to rich fishing grounds (Andreassen 2003). The Finnes site is located at an altitude of 5m asl and ca. 24m from the present coast. The vegetation of the site today is almost treeless grassland with herbs and grasses typical of rough grazed pasture.

Rescue excavations at Finnes were undertaken by Tromsø University Museum in 2011 following extensive site disturbance due to unapproved “excavation” by a local historian. The excavation in 2011 CE included 0.5m^2^ test pits and a 3.6×1.3m trench in the floor area of a turf structure from the final phase of occupation, a nineteenth-century fishing settlement (see the plan in S4, stratigraphy in Figure 1, and the description and chronology in S5). The site was found to be occupied over two intervals: the early metal period (ca. 400-202 cal. BCE) and medieval period (1279-1358 cal. CE) with an unoccupied period in between (Wickler 2016, 2026; Calibration: Bronk Ramsey 2009, 2021; Reimer et al. 2020). A rich faunal assemblage (S6) resulted from the initial use of the site in the early metal age (Layer 7 from 2011 CE excavation, and the site is interpreted as a temporary fishing and marine hunting site most likely used by the coastal Sámi. The importance of marine exploitation resumed in the initial medieval period (Layer 6 in 2011 CE excavation) when the site was re-established after a lengthy hiatus (Wickler 2016, 2026). The most intensive occupation was observed in the second phase of medieval occupation (1299-1400 cal. CE, Layer 3 in 2011 CE excavation) with artifacts such as double-sided antler combs and bakestones. The cultural deposit from this phase has been severely truncated by disturbance from construction of turf structures in the 19th century fishing village. No animal bone was preserved in this layer due to an abundance of acidic turf. The evidence from layer 3 (2011 CE excavation) suggests that this fishing settlement expanded in the final phase of the medieval occupation (Wickler 2016, 2026). Permanent Norse settlement and mound formation can be directly linked to commercialized cod fishing and the stockfish trade as demonstrated by the presence of imported objects. Finnes exemplifies the rapid expansion of Norse fishing settlement into coastal Sámi areas along the outer coast of Finnmark that commenced around the middle of the 1200s and its impact on interethnic maritime socioeconomic relations. Critically for this study the stratigraphies of both sites revealed no intrusive material or mixing and no earthworms were encountered at either site.

## Results

The occupation of Baŋkgohppi is estimated to be 2448 to 1826 cal. BCE (Martens et al. 2017). Figure 2 shows sedaDNA and macrofossils of edible plants, mammals, fish, shellfish, echinoderm, birds, algae, and annelids. Of 13 samples taken in the field, 11 of them had sedaDNA. Of the 79-plant identified using the P6-Loop (S7), 31 are regarded as edible taxa (S8, frequent taxa in Figure 2). The 12S primer produced 159 taxa including algae, bacteria, birds, chromista, eukaryotes, fish, mammals, plants, and protozoa of which 24 edible taxa were identified, and 21 included (seven plants, four algae, six mammals, two birds, and two fish) after removing contaminants and aggregating higher taxonomic levels to the levels where nutritional values are available (S9, S8, selected taxa in Figure 2). From the 16S primer, 17 taxa were identified including mammals, fish, shellfish, Echinodermata, birds, and annelids (S10) which included 12 edible taxa (four mammals, three fish, two shellfish, one bird, and two annelids) (S8, selected taxa in Figure 2). Crowberry (*Empetrum nigrum*), Atlantic cod (*Gadus morhua*), common ling (*Molva molva*), and common guillemot (*Uria aalge*) were identified by 2 primers (P6-Loop and 12S or 12S and 16S) and macro/microfossils whilst taxa that were identified by 2 primers or a primer and microfossil included blueberry (*Vaccinium myrtillus*), fireweed (*Chamaenerion angustifolium*), pine (*Pinus sylvestris*) (P6-Loop and 12S), bowhead whale (*Balaena mysticetus*), beaver (*Castor fiber*), reindeer (*Ragnifer tarandus*) (12S and 16S), willow (*Salix*) (P6-Loop and microfossils), Greenland seal (*Phoca groenlandica*) (12S and macrofossils), saithe (*Pollachius virens*), and periwinkle (*Littorina*) (16S and macrofossils).

**Figure 2.**
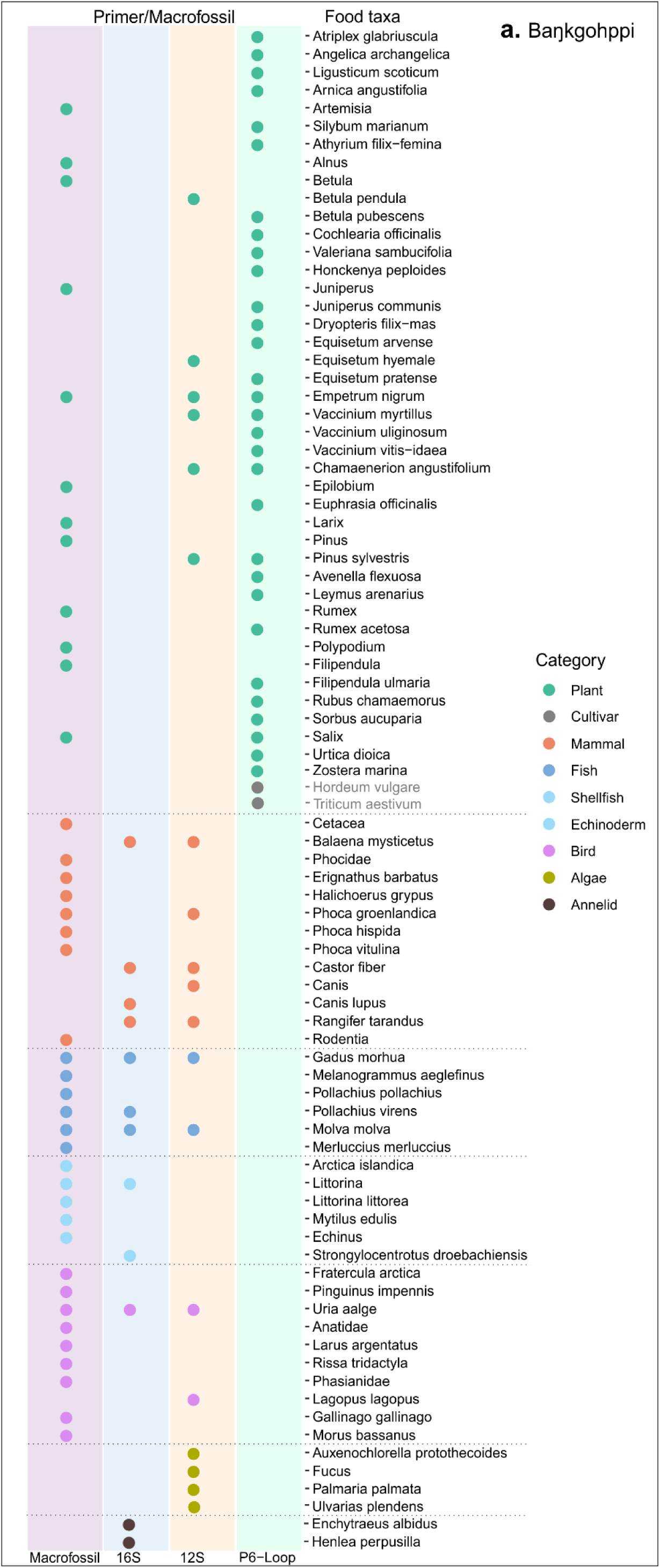
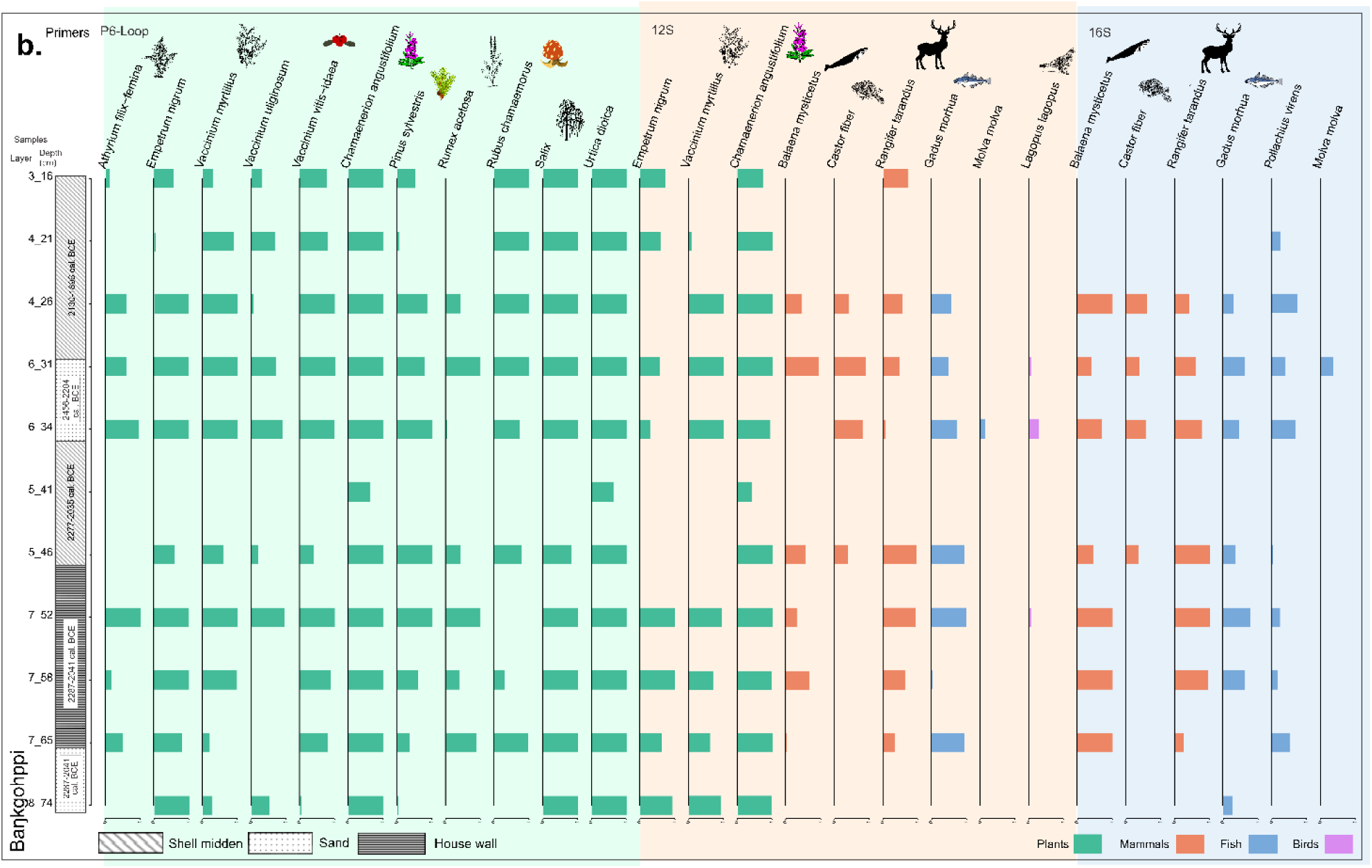
Identified edible taxa at Ba.kgohppi. Edible taxa identified at Baŋkgohppi including sedaDNA and macrofossils (2a), sedaDNA only using three primers P6-Loop, 12S, and16S (selected taxa 2b, complete figure in S8).

Using all the primers, 30 out of 52 edible taxa excluding annelids and duplicates were present in more than five samples including three species of cowberry/blueberry (*Vaccinium myrtillus*, *Vaccinium uliginosum*, *Vaccinium vitis-idaea*), barley (*Hordeum vulgare*), wheat (*Triticum aestivum*), meadowsweet (*Filipendula ulmaria*), stinging nettles (*Urtica dioica*) and silver birch (*Betula pendula*) (S8). The mammals present were the bowhead whale, reindeer, and dog/wolf, and the fish were cod (Gadidae) and saithe as well as the green sea urchin (*Strongylocentrotus droebachiensis*) (S8), the spines of which were also found. Sea urchin was a major food item in the upper layers while it was sea urchin and cod for the lower layer. Layers in between (layers 4-7), the major food items were whale and reindeer. Beaver was also found in 4 samples at Baŋkgohppi (S8, Figure 2). S11 shows the top ten plant families and their relative abundance based on total reads at Baŋkgohppi. Betulaceae was the major family throughout the sequence suggesting an environment not dissimilar to that on the site today. A significant taxon in the plant data was marine eel grass (*Zostera marina*) which was only present in the lowest two samples (S7) and it might be eaten in Norway as one of the common names in Norwegian was matløk (food onion) in 18 CE (Alm 2003).

Stratigraphy at Finnes was revised based on the contexts revealed (Figure 1, S5), and both macrofossils and sedaDNA identified were from the medieval period (1279-1400 cal. CE). The 11 levels from the site that had sedaDNA showed less complementarity between the sedaDNA and the macrofossils than at Baŋkgohppi (Figure 3, S12-15) and edible taxa were found in all food groups; plants, mammals, fish, shellfish, birds, amphibians, and the edible algae dulse (*Palmaria palmata*).

**Figure 3.**
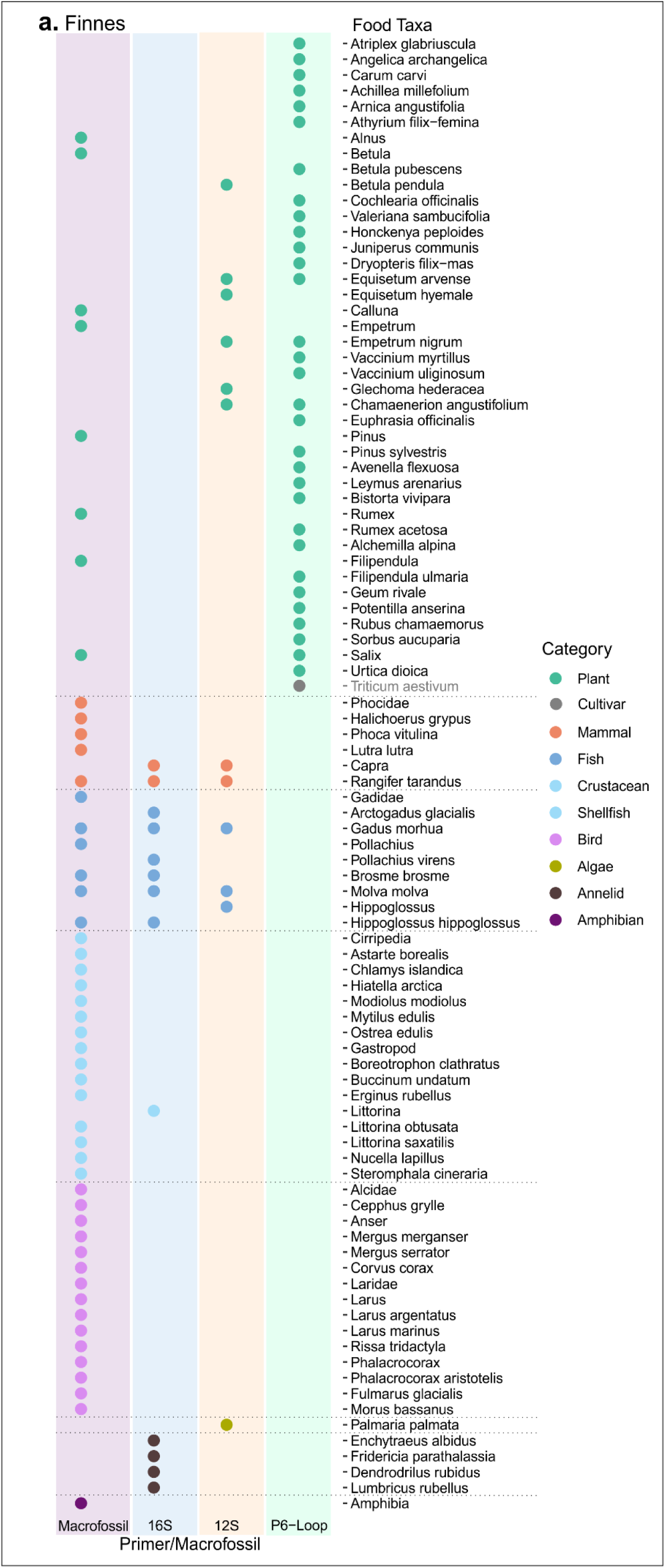
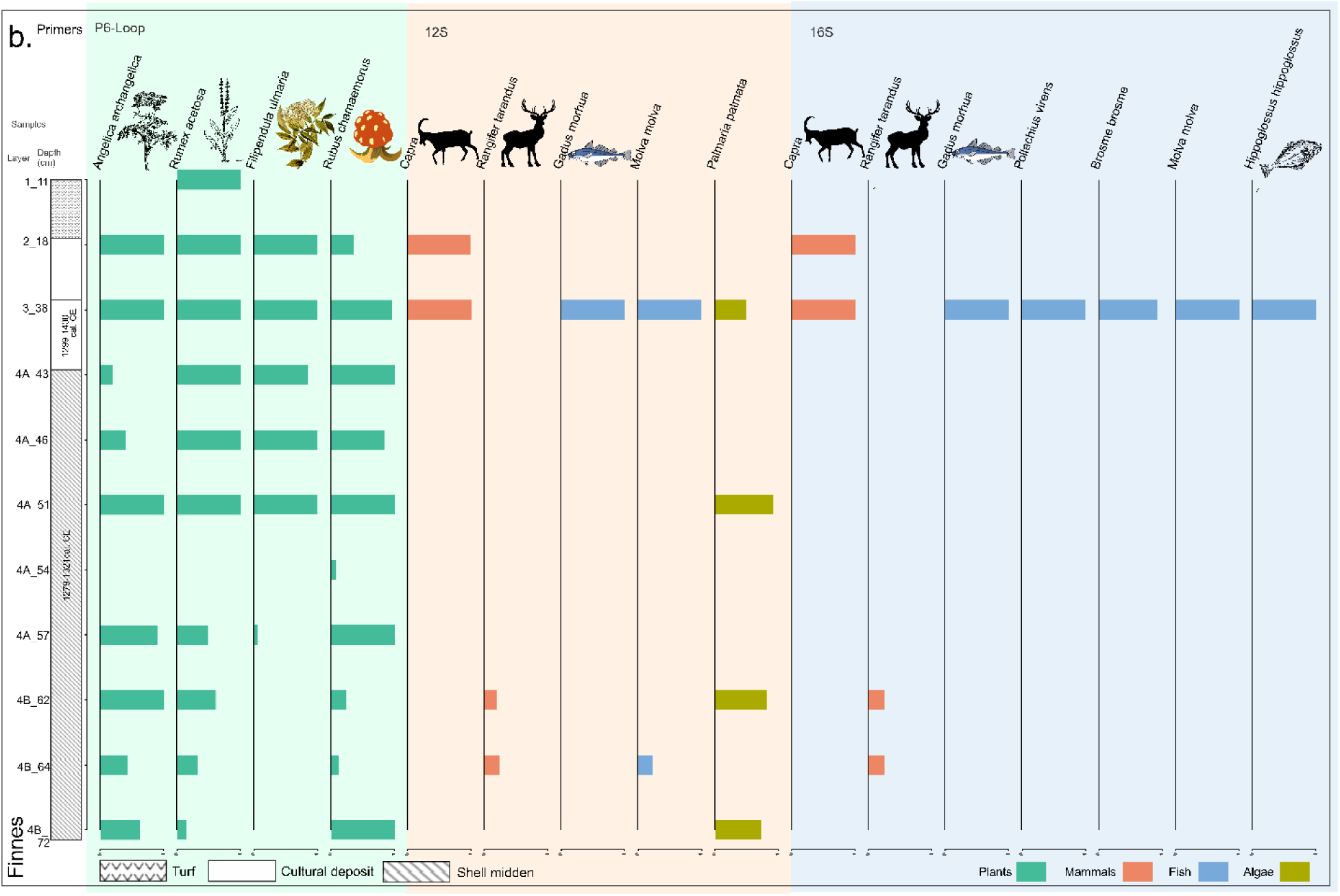
Identified edible taxa at Finnes. Edible taxa identified at Finnes including sedaDNA and macrofossils (3a), sedaDNA only, using three primers P6-Loop, 12S, and16S (selected taxa 3b, complete figure S15).

The number of plant taxa identified using the P6-Loop primer dataset was 68 (S12). A total of 32 edible taxa remained after removing contaminants and duplicates (S15). The 12S primer produced 157 taxa which included algae, bacteria, chromista, eukaryotes, fish, mammals, plants, and protozoa. This included 13 edible taxa (six plants, three mammals, three fish, and one alga, S13) and 12 edible taxa at species level except goats and halibuts (*Hippoglossus*, two mammals, three fish, six plants, and one alga, S15). The 16S primer revealed 19 edible taxa (S14) and 13 after removing contaminants and taxa at higher levels than genus (two mammals, six fish, and one shellfish, four annelids, S15). Reindeer, Atlantic cods, common lings, and halibuts (at genus level) were identified by 12S, 16S and macrofossil whilst goat (*Capra*) was in 12S and 16S, pollack (*Pollachius*) and cusk (*Brosme brosme*) were in 16S and the faunal assemblage (Figure 3). The 16S data reveals four samples dominated by only one food type although one has seven (goats, Arctic cods, Atlantic cods, saithe, cusks, common lings, and Atlantic halibuts) excluding annelids (S15), and this may represent swamping by the most common DNA template sequences. The top 10 plant families and their relative abundance based on the total reads at Finnes are shown in S16.

The shotgun sequencing of three samples (GB_6_31, GB_5_46, IG_4B_72) and negative library control resulted in 98.9 million reads. Across all three samples, 29 plant taxa and 5 animal taxa were identified with damage patterns supporting their ancient origin (Figure 4, S17-23). The only taxa identified in the negative library control were *Homo* and *Sus* which were not present in our dataset. All plant taxa identified through shotgun sequencing were also present in the metabarcoding dataset while only the two rodent taxa identified through shotgun (*Microtus* and Cricetidae) were absent from the 16S and 12S datasets. These identifications could represent true positives of a Microtus species or misassigned *Castor fiber* sequences since this taxon was absent from our reference database, but present in the metabarcoding data (S24).

**Figure 4.**
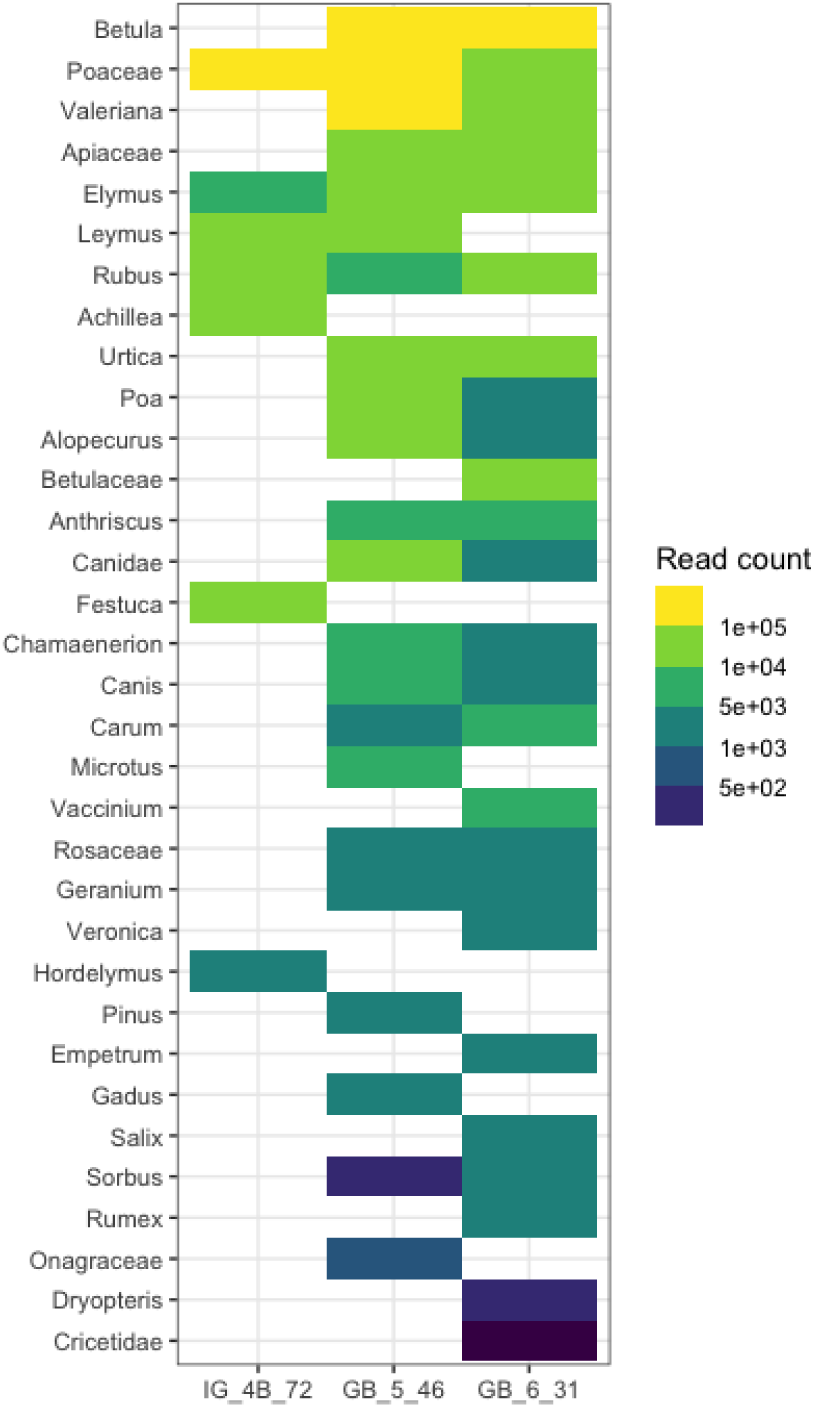
All taxa identified by shotgun sequencing with a significant damage pattern at Baŋkgohppi (GB_5_46 and GB_6_31) and Finnes (IG_4B_72).

## Discussion and dietary implications

This first application of sedaDNA analysis to archaeological sites in northern Norway has implications at two levels, firstly in the occupation and activities undertaken at each site, and secondly specification of the inhabitants’ diet. The sedaDNA from the hunter-gatherer-fisher site of Baŋkgohppi adds significantly to the plants and the mammals. Of particular significance is the bowhead whale, as it is the only baleen whale species that resides year-round in the Arctic and subarctic regions and lives in close association with sea ice. So, if not from scavenged carcasses, this indicates deep-sea fishing with harpoons as is known to have been used by palaeo-inuit peoples at about this date (Seersholm et al. 2016). Given their geographical distribution and the currents in the Barents Sea, they are the least likely of the whales to be stranded on the Finnmark coast although common in Svalbard (Lønne and Fuglei 1997). The site also confirms increasing evidence that beaver was a common component of the Finnmark environment in the mid-Holocene (Brown et al. 2022b) as well as willow ptarmigan (*Lagopus lagopus*). The strong occurrence of shrubs with berries which ripen as late as September and birds which breed in the early summer suggests that the site must have been occupied from at least May to October but most likely year-round. The site can be compared with a Gressbakken phase house (R3) at the site of Mortensnes on the northern shore of Varanger Fjord. Bones recovered from this house were whale, reindeer, Greenland seal, common seal, harbour porpoise, walrus, dog, shag, cormorant, raven, Atlantic cod, and pollock (Brown et al. 2022b; Schanche 1989). Two species recovered at Mortensnes but not at Baŋkgohppi, hare and otter, may reflect the more open environment and a nearby stream and lake.

At Finnes, the occurrence of domesticates (cattle and goats) in the medieval period (1190-1320 CE) is important as it suggests that although Finnes was a fishing community, it was permanently settled at this time. The occurrence of reindeer could also be due to hunting and/or herding. There is a vigorous debate concerning the date of the shift from hunting to herding in northern Fennoscandia, but some authorities suggest that this could be as early as the late Medieval period (15^th^ century CE); although it is argued that there may have been small-scale herding from the late Iron Age onwards (Salmi 2022). The most comparable site at Mortensnes for the Medieval period (R17, grave) yielded sheep/goat, cattle, seal, reindeer, pig, dog, a wide range of marine fish, and nine bird species dominated by black-legged kittiwake and rock ptarmigan. This probably represents the larger bird-cliff near Mortensnes as well as the open environment (Brown et al. 2022b) – a situation very comparable to the Finnes site discussed here.

At both sites, sedaDNA doubled the food species revealed and added species to all food groups (algae). Terrestrial mammals such as reindeer as well as wheat were also added as foods species to both sites by sedaDNA. It also added beaver and bowhead whales to the prehistorical site of Gressbakken. These food taxa were also found at another co-eval site across the Varanger Fjord at Mortensnes (Brown et al. 2022b). Goats were added to the medieval phase of the Finnes site by the sedaDNA analysis. Although cultivars such as Triticum and Hordeum were present in several samples at both sites (S7, 8, 12, 15, 25, 26) and not in negative controls, they were excluded from the nutritional analysis following the generally accepted geographical range of cereals even though cereal pollen has been recovered from the region around c.800-450 BCE (Jensen and Elverland 2009) and c. 0 BCE/CE (Brown et al. 2022b) as well as the results of the shotgun analysis. Regardless of the pipeline, those in samples GB_6_31, GB_5_46, and IG_4B_72 had such low read occurrences that there was a high probability that they were due to random matches (read counts for Hordeum vulgare 208, 90, 118 and for Triticum aestivum 654, 185, and 264 in the samples GB_6_31, GB_5_46, and IG_4B_72, whose read count cut-off were 1607, 1341, 1700, respectively, S17-19). Pine was in a negative control for Baŋkgohppi; however, it was kept in the analysis since they were in 10 of the 11 samples that had sedaDNA using P6-Loop (layers 3-8, S27) as well as the known presence in the region at this time (Clarke et al. 2019).

Food group reconstruction with and without sedaDNA shows macrofossils and sedaDNA complement each other (Figure 5a). The top three food categories changes from bird, plant, mammal and shellfish for macro/microfossils to plant, algae, mammal for sedaDNA, but combining both macro/microfossils and sedaDNA reveals all the food items, showing plant, bird, and mammal as the top three food categories in Baŋkgohppi. At Finnes, the top three food categories from macro/microfossils were shellfish, bird, and mammal whereas those of sedaDNA were plant, mammal, fish, and algae excluding annelid. This is due to primers not designed specifically for birds and shellfish, so they maybe underrepresented in sedaDNA. The differences between macro/microfossils and sedaDNA are partially due to the identification of species at different taxonomic levels. Since edible plant rating was used to decide which taxa to keep for the analysis of plants, taxa above the species levels for plants were generally removed from the analysis. Interestingly when food items from both macro/microfossils and sedaDNA were combined, plant, fish, and mammal had similar results at both sites. Although major food categories were plant, bird, and mammal for Baŋkgohppi and plant, shellfish, and bird for Finnes, food items for both sites satisfied all the requirements of macronutrients, minerals, and vitamins except for biotin (Figure 5c).

**Figure 5.**
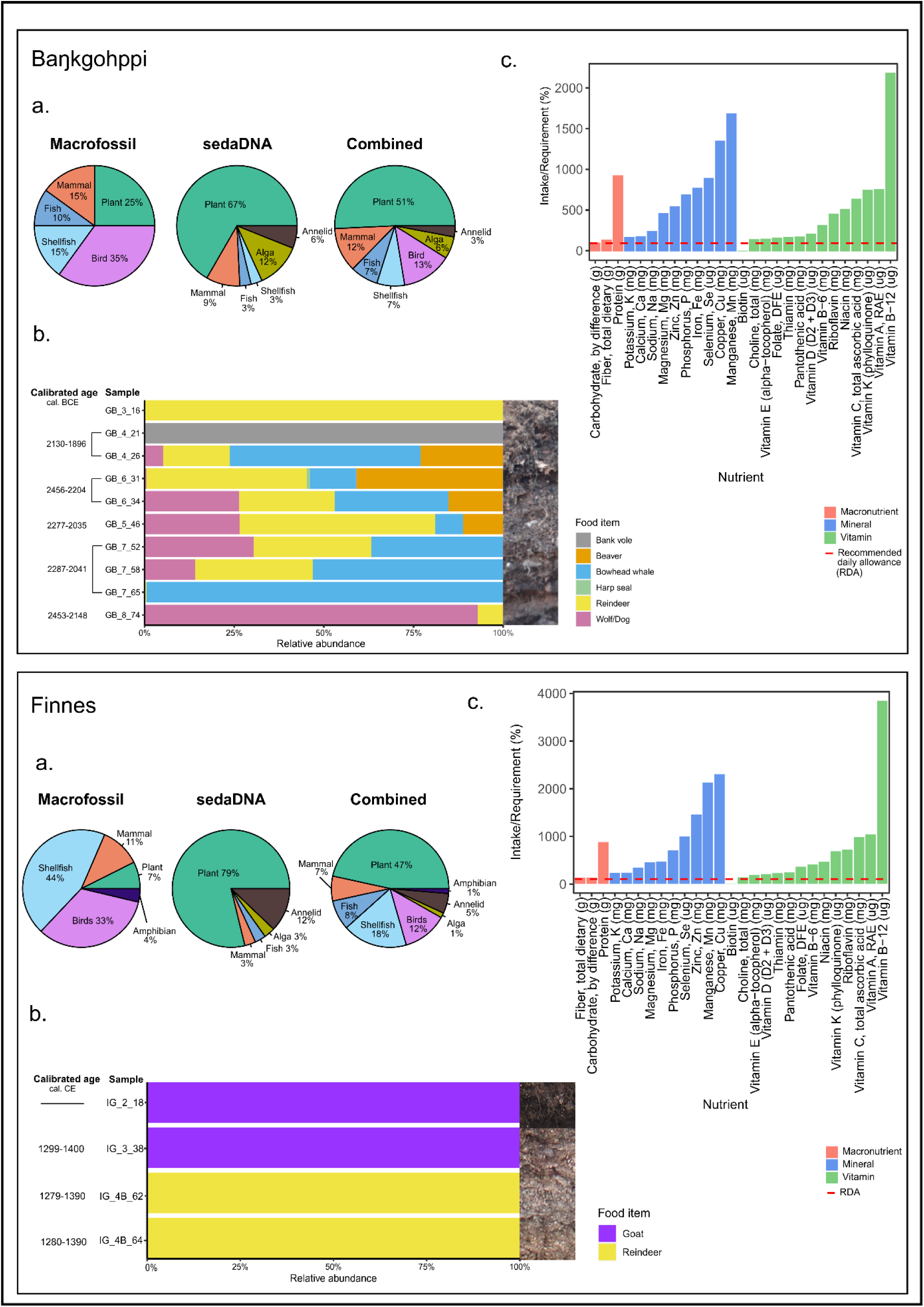
Food group reconstruction with and without sedaDNA and a nutritional analysis example at Baŋkgohppi and Finnes. 5a. Proportions of food groups identified by macrofossils, sedaDNA, and combined after removing duplicates 5b. Relative abundance of mammals. 5c. Nutritional analysis (intake of nutrients relative to RDA) using the food items found at each site when cooked.

For nutritional analyses, we highlight nutrients that were lower than RDA and higher than tolerable upper intake level (TUIL). The full list of food included in the analysis, percentages of RDA for each nutrient, and what food items contributed to each nutrient intake are shown in S28-35. Nutritional calculations were similar between raw and cooked food materials except estimations with raw materials having lower calorie intake (Table 1, S30, 31). Although multiple micronutrients were above TUIL, this may be over-estimated due to the assumption of almost all available internal organs were consumed. In general, the combined (sedaDNA and macrofossils) nutrition calculation was high in protein but deficient in biotin for both sites although Finnes had lower overall calorific value than Baŋkgohppi. Assuming the residents at both sites ate internal organs and different parts of the bodies of mammals, these parts were included in the nutrition calculation, resulting in higher calorie intake.

**Table 1.**
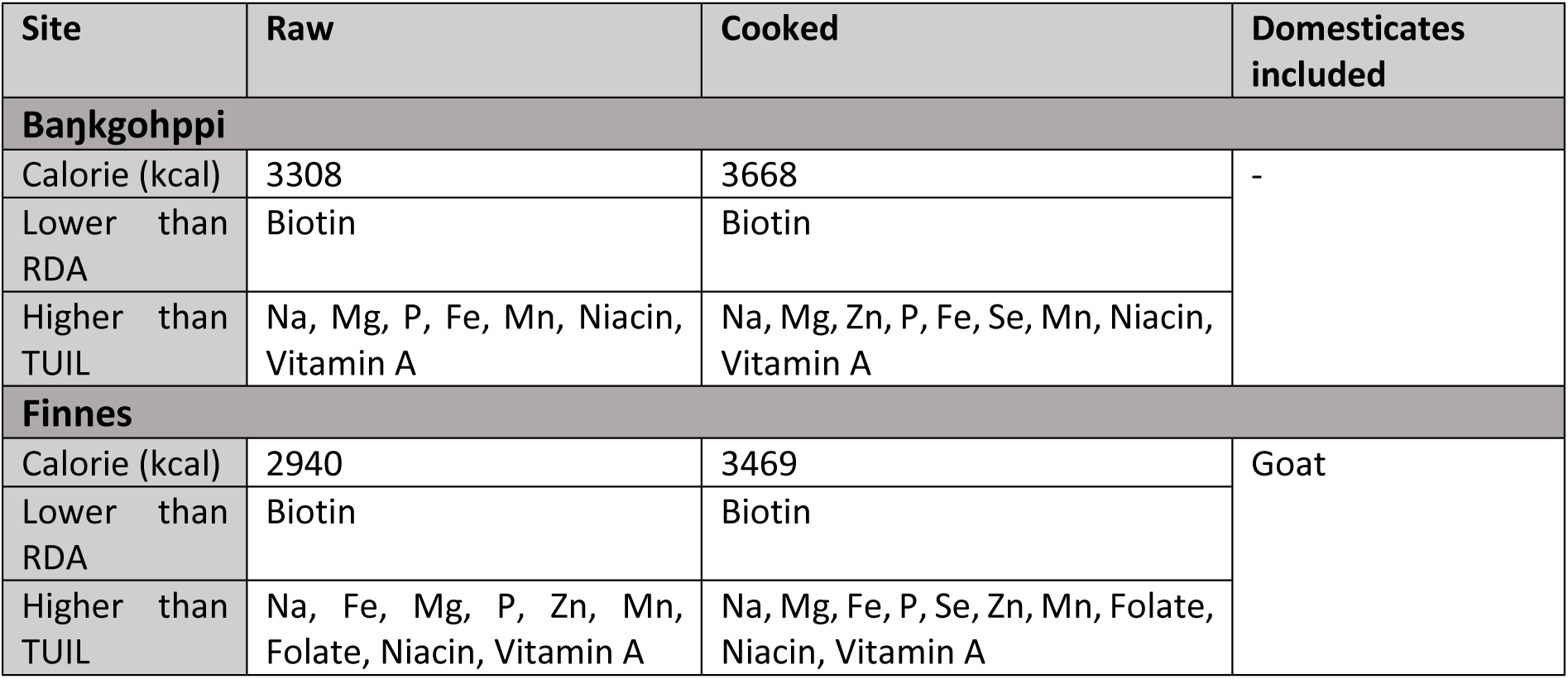
The summary of the nutritional analysis for Baŋkgohppi and Finnes.

However, biotin might be underrepresented as we assume 100g of each food item was consumed as well as lack of nutritional information for most of the bird species found. Biotin (vitamin B7) works as a cofactor to help enzymes break down macronutrients such as fats, carbohydrates, and proteins as well as help regulate cell signaling (NIH 2022). Hence, the residents of both sites must have compensated by eating higher amounts or other food items that are rich in biotin. Biotin is rich in eggs and livers (NIH 2022), which raises the question of the exploitation of bird-cliffs and eggs. The abundance of birds and particularly seabirds that nest on coastal cliffs such as the common guillemot, great auk (*Pinguinis impennis*), puffin (*Fratercula arctica*), herring gull (*Larus argentatus*), and black kittiwake (*Rissa tridactyla*) strongly implies the utilization of bird cliffs for both meat and eggs. There was a bird-cliff only 1100 m to the east of the Finnes site, so the residents might have consumed bird eggs. Also, the removal of cattle from the analyses is significant since if it were included, medieval Finnes people would have satisfied the biotin required for survival as beef especially liver is high in biotin (NIH 2022; Schweigert et al. 1943). Regardless, biotin can be synthesized by bacteria in the human body (Said 2009); hence, biotin deficiency is less likely for the residents of both sites. Residents at both sites seem to have had a well-balanced diet seasonally over the year and are unlikely to have resulted in any significant nutritional diseases or conditions.

Although both Baŋkgohppi and Finnes occupants had potentially nutrient-rich broad-spectrum diets, there would have been seasonal variation based on the availability of resources and particularly the plant species since most of them are harvested between May and September (S36). At both sites, several micronutrients were supplied by mostly plants. Those nutrients that were supplied over 50% by plants were vitamin K, vitamin C, Copper, Manganese, and Calcium in Baŋkgohppi whereas they were vitamin K, for Finnes (detail in S32-35). However, these nutrients can be supplied by reindeer, cod, and beaver if viscera are eaten in sufficient quantity, and especially from reindeer and cod in winter (Hassan et al. 2012; Norwegian Food Safety Authority 2026). Vitamin C and K can be supplied by seaweed, and well-known source of vitamin C is the skin of some marine mammals (Aubin and Geraci 1980).

## Conclusion

In the first study comparing sedaDNA and macrofossils from middens in northern Eurasia, we show that adding taxa identified from sedaDNA doubles the edible foods that were consumed in two important periods (Gressbakken and Medieval Periods) within the last 3500 years. Even though traditional macrofossil analyses can provide insight into past foodways, sedaDNA can add significantly more and reduce the effect of preservation bias. When macrofossils and sedaDNA are compared separately, they reveal different aspects of foodways which, when considered together, indicate an increase in dietary breadth, with implications for fishing, hunting, and gathering activities. Notwithstanding that our reconstructed diets could still be incomplete, the nutritional analysis points to a deficit in micronutrients that could have been satisfied by systematic gathering of eggs from coastal bird cliffs, for which there is strong circumstantial and ethnographic evidence. The sedaDNA has also provided evidence that in the last Stone Age Gressbakken culture deep-sea fishing was likely practiced. The incorporation of sedaDNA data may also have implications for other important aspects of subarctic life such as the seasonality of settlement and reindeer utilisation and sedentism (year-round occupation).

## Methods

After cleaning/removing the surface of the sediment, 50 ml sterile tubes were inserted to collect sediment samples (Figure 1d). There were 13 and 18 samples collected from Baŋkgohppi and Finnes, respectively. The details of the methods for sedaDNA extraction, amplification, and library preparation are described in Rijal et al. (2021) for plants and Garcés-Pastor et al. (2022) for mammals.

For sedaDNA extraction, the DNeasy PowerLyzer PowerSoil kit (Qiagen: 12855-100) was used. 0.25-0.35 g of sediments from the 50 ml tubes were sub-sampled for DNA extraction. Using 750 μL of PowerBead solution, two rounds of bead beating (20 seconds at 4.0 m/s) were performed. After adding 87 μL of lysis solution containing Solution C1, DTT, and proteinase K, the sub-samples were vortexed and incubated at 56 ^◦^C overnight. Adding 250 μL of Solution C2, the samples were vortexed and incubated in a fridge for 10 minutes. Transferring 800 μL of the supernatant to 250 μL of Solution C3, the same vortex and incubation steps were taken followed by the addition of 880 μL of supernatant to 1400 μL of Solution C4 and vortex. 650 μL of the Solution C4 and the supernatant mix was loaded onto a MB spin column and centrifuged at 10,000x g for one minute. After repeating the centrifugation steps, 500 μL of Solution C5 was added to the column and centrifuged, followed by addition of 65 μL elution butter to the column, incubating for 10 minutes at the room temperature, and centrifugation at 10,000x g for one minute to recover 120-125 μL of DNA extract for each sample.

For PCR amplification, 32 μL of master mix solution containing 10x Gold buffer, MgCl2, BSA, dNTPs, and AmpliTaq Gold, 4 μL of forward and reverse primer mix, and 4 μL of DNA extract were added to PCR plates. In total, there were 31 samples, 5 negative controls, and 1 positive control on the plates. Two different sets of primers were separately used: one set of “gh” primers targeting the vascular plant trnL p6-loop (Taberlet et al. 2007) and primers targeting the mitochondrial 12S (batra) (Hudson et al. 2023) and 16S gene (MamP007F and MamP007R), (Giguet-Covex et al. 2014). Negative controls included those at sedaDNA extraction steps for each site, one at the DNA extract loading step to the plates, one in the ancient DNA laboratory, and one in the post-PCR laboratory. For PCR, activation of the enzyme was at 95 ^◦^C for 10 minutes, followed by 45 cycles of denaturing at 95 ^◦^C for 30 seconds, annealing at 50 ^◦^C for 30 seconds, and extension at 72 ^◦^C for 1 minute with the final extension at 72 ^◦^C for 7 minutes.

PCR products were pooled separately for plant primers and mammal primers following the Illumina TruSeq DNA PCR-Free protocol (Illumina, Inc. 2017), and the libraries were cleaned with the Qiagen MinElute PCR purification kit (Qiagen GmbH, Hilden, Germany). Sequencing of the libraries was performed by 2x 150-cycle mid-output flow cells on the Illumina NextSeq platform at the Genomics Support Centre Tromsø (GSCT) at the University Hospital of North Norway for plants and mammals separately. The full detail of the bioinformatics pipeline is described by Rijal et al. (2021), but in summary, in order to determine taxa, sedaDNA sequences were compared with 4 reference libraries: ArcBorBryo (Soininen et al. 2015; Sønstebø et al. 2010; Willerslev et al. 2014), EMBL 143 (European Molecular Biology Laboratory database release 143), PhyloAlps (Garcés-Pastor et al. 2022), and PhyloNorway (Alsos et al. 2020), and only Taxa that have 100% match were kept. Duplicates after the comparison with reference libraries were removed by adding total reads and keeping the highest weighted repeats and total repeats. R version 4.2.3 (R Core Team 2023) and RStudio version 2023.03.1+446 (Posit team 2023) were used for the analysis of the generated data. The sedaDNA is shown as percentages of weighted repeats. The taxa that were in negative controls as well as cultivars that were not considered present at the time of the site occupancy were removed from the analysis as they were regarded as contamination (detail listed in S37). Also, taxa above the species level and non-edible taxa were removed or species were identified from distributions or using BLAST (NCBI 2023) (Figure 2b, 3b, S37).

Aliquots of the same DNA extracts used for metabarcoding were used to prepare single-stranded shotgun libraries at the Alfred Wegener Iinstitute (Potsdam, Germany). Three samples (GB_6_31, GB_5_46, and IG_4B_72), a library blank control, and 19 other sedaDNA extracts from northern Norway not included in the study were prepared with the single-stranded library method specified for degraded ancient DNA (Gansauge and Meyer 2013; Gansauge et al. 2017) with modifications described in Schulte et al. (2021). DNA extracts were quantified with Qubit 4.0 fluorometer (Invitrogen) and 15 ng of DNA were used as the starting material. After adapter ligation, libraries underwent an indexing PCR with 10 to 13 PCR cycles and unique combinations of P5 and P7 Illumina primers. Then, PCR products were purified with the MinElute PCR Purification Kit (Qiagen, Germany) and libraries’ concentrations were measured with qPCR (Gansauge and Meyer 2013) containing 1x Maxima™ SYBR™ Green (Thermo Scientific, Germany). The library blank was amplified with the maximum number of PCR cycles used for the samples. The libraries’ fragment sizes were checked on the Agilent TapeStation using the D1000 ScreenTape (Agilent Technologies USA) and samples were pooled equimolarly. Pooled samples were sequenced with a customized sequencing primer CL72 on a NextSeq2000 Illumina device using paired-end mode (2×100 bp) at AWI Bremerhaven (Germany).

After sequencing, the quality of the raw sequencing data was checked using FastQC v0.12.0 (Andrews 2010) and reads were then merged, adapter sequences were removed, and reads less than 35 bp were discarded using *fastp* v0.23.4 (Chen 2023). Taxonomic annotation was performed by the multi-mapping the reads to three different databases using bowtie2 (Langmead et al. 2019); the bacterial genome taxonomy database (GTDB, release 232), the 1,541 assembled genome skims of arctic vascular plants from PhyloNorway (Alsos et al. 2020), and a custom database of 42 metazoan genomes retrieved from EMBL of probable fauna and contaminants (S24). Only mappings with ≥97% identity were retained and reads were individually assigned to the lowest common ancestor shared among all mappings using a custom python script. Taxonomic assignments were collapsed to the genus and family-levels with a minimum of 200 reads per sample. Bacterial assignments, as well as reads assigned at higher than family-level, were not considered for downstream analysis. Cytosine deamination patterns were characterized for each taxon identification to support their ancient origin using pyDamage (Borry et al. 2021) grouping all reference contigs together (-g option). A taxonomic assignment was considered ancient if its assigned reads fit the GLM model of pyDamage with a p-value ≤ 0.01, a residual standard mean error ≤ 0.01, and a magnitude of damage ≥ 0.02. A taxonomic identification was discarded if it composed < 1% of a family’s total read count within a sample to avoid possible phantom alignment issues of closely related taxa.

For nutritional analyses, taxa identified by both metabarcoding and macro/microfossils were aggregated. Taxa that could not be assigned to specific food items, including those identified only at higher taxonomic levels, contaminants, and taxa for which nutritional information was unavailable, were excluded (S37-39). Although *Pinus* was also present in a negative control for Baŋkgohppi, C001 (when sampling in the ancient DNA lab), it was kept for the analysis as the presence of *Pinus* in the region during the late Holocene is known (Alsos et al. 2024; Clarke et al. 2019). In the dietary analysis, only those species with an edible rating of above 3 (PFAF 2022) and those considered edible in northern Norway from ethnographic evidence were included in the analysis. To assess effects of the method on the dietary and nutritional estimation, we compare 1) taxa identified based on different primers and macrofossil and 2) nutritional calculation based on taxa identified at each site. For nutritional analysis, if the food items satisfied recommended daily allowance (RDA) (S40) as well as what food items were contributing to each nutrient were calculated using R package NutrienTrackeR (Rodriguez-Martinez et al. 2023) based on the Canadian Nutrient File (Health Canada 2015). For food items for which nutritional values are available, nutritional values for 100g of each item were used except for organs, fat/oil, leafy vegetables, and seeds (one portion for these items were used and adjusted them to add up to 100g for each food species). For those food items whose nutritional values were available for both raw and cooked, they were calculated separately as well as including and excluding wheat (raw with wheat, cooked with wheat, raw without wheat, cooked without wheat).

