## Supplementary materials for "First Broad-spectrum Sámi and Norse diets in northern Norway over the last 4000 years revealed by sedaDNA"

S1. A plan of the Baŋkgohppi site (modified from Martens et al. 2017).

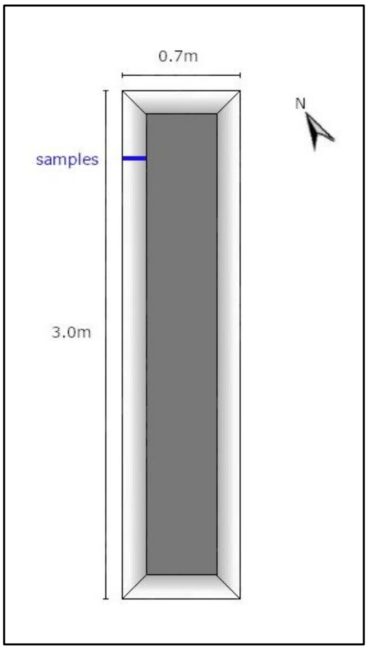

S2. List of flora and fauna excavated at the Baŋkgohppi site in the *InSituFarms* project (Martens et al. 2017; Calibration: Bronk Ramsey 2009, 2021; Reimer et al. 2020). ● indicates presence. * indicates only presence without indication of layers.

<table>
<tr><th rowspan="2">Category</th><th rowspan="2">Common name</th><th rowspan="2">Scientific name</th><th colspan="11">Layer & $^{14}C$ dates (Cal. BCE)</th></tr>
<tr><th>1</th><th>2</th><th>3</th><th>4<br>2130-<br>1896</th><th>5a<br>2277-<br>2035</th><th>5b<br>2297-<br>2051</th><th>6<br>2456-<br>2204</th><th>7<br>2287-<br>2041</th><th>8<br>2453-<br>2148</th><th>9</th><th>10</th></tr>
<tr><td>Plants</td><td>Alder</td><td>Alnus</td><td></td><td>●</td><td>●</td><td></td><td colspan="2"></td><td>●</td><td></td><td></td><td></td><td></td></tr>
<tr><td></td><td>Birch</td><td>Betula</td><td>●</td><td>●</td><td>●</td><td>●</td><td colspan="2">●</td><td>●</td><td>●</td><td>●</td><td>●</td><td></td></tr>
<tr><td></td><td>Spruce</td><td>Picea</td><td></td><td>●</td><td></td><td></td><td colspan="2"></td><td></td><td></td><td></td><td></td><td></td></tr>
<tr><td></td><td>Pine</td><td>Pinus</td><td>●</td><td>●</td><td>●</td><td></td><td colspan="2">●</td><td></td><td>●</td><td>●</td><td>●</td><td></td></tr>
<tr><td></td><td>Larch</td><td>Larix *</td><td></td><td></td><td></td><td></td><td colspan="2"></td><td></td><td></td><td></td><td></td><td></td></tr>
<tr><td></td><td>Cedar</td><td>Juniperus</td><td></td><td></td><td></td><td></td><td colspan="2">●</td><td></td><td></td><td></td><td></td><td></td></tr>
<tr><td></td><td>Willow</td><td>Salix</td><td></td><td></td><td></td><td></td><td colspan="2"></td><td>●</td><td></td><td></td><td></td><td></td></tr>
<tr><td></td><td></td><td>Ericales</td><td>●</td><td>●</td><td></td><td></td><td colspan="2">●</td><td></td><td>●</td><td></td><td></td><td></td></tr>
<tr><td></td><td>Black crowberry</td><td>Empetrum nigrum</td><td></td><td></td><td>●</td><td>●</td><td colspan="2"></td><td></td><td></td><td></td><td></td><td></td></tr>
<tr><td></td><td>Umbellifer</td><td>Apiaceae</td><td></td><td></td><td>●</td><td>●</td><td colspan="2">●</td><td></td><td></td><td></td><td></td><td></td></tr>
<tr><td></td><td>Sagebrush</td><td>Artemisia</td><td></td><td></td><td></td><td></td><td colspan="2"></td><td>●</td><td>●</td><td></td><td></td><td></td></tr>
<tr><td></td><td>Alpine Saw-wort</td><td>Saussurea alpina</td><td></td><td></td><td></td><td></td><td colspan="2">●</td><td></td><td></td><td></td><td></td><td></td></tr>
<tr><td></td><td></td><td>Aster.</td><td></td><td>●</td><td>●</td><td></td><td colspan="2">●</td><td>●</td><td>●</td><td>●</td><td></td><td></td></tr>
<tr><td></td><td></td><td>Cicor.</td><td></td><td>●</td><td></td><td></td><td colspan="2">●</td><td>●</td><td>●</td><td></td><td></td><td></td></tr>
<tr><td></td><td>Pinks</td><td>Caryophyllaceae</td><td></td><td></td><td>●</td><td>●</td><td colspan="2">●</td><td>●</td><td>●</td><td></td><td></td><td></td></tr>
<tr><td></td><td>Buttercup</td><td>Ranunculus</td><td></td><td>●</td><td>●</td><td></td><td colspan="2">●</td><td>●</td><td>●</td><td>●</td><td></td><td></td></tr>
<tr><td></td><td></td><td>Chamerion</td><td></td><td></td><td></td><td></td><td colspan="2"></td><td></td><td></td><td>●</td><td>●</td><td></td></tr>
<tr><td></td><td>Sedge</td><td>Cyperaceae</td><td>●</td><td>●</td><td>●</td><td>●</td><td colspan="2">●</td><td>●</td><td>●</td><td>●</td><td></td><td></td></tr>
<tr><td></td><td>Sedge</td><td>Carex *</td><td></td><td></td><td></td><td></td><td colspan="2"></td><td></td><td></td><td></td><td></td><td></td></tr>
<tr><td></td><td>Catch-fly</td><td>Drosera</td><td>●</td><td>●</td><td></td><td></td><td colspan="2"></td><td></td><td></td><td></td><td></td><td></td></tr>
<tr><td></td><td>Legume</td><td>Fabaceae</td><td></td><td></td><td></td><td>●</td><td colspan="2"></td><td></td><td></td><td></td><td></td><td></td></tr>
<tr><td></td><td>Grass</td><td>Poaceae</td><td>●</td><td>●</td><td>●</td><td>●</td><td colspan="2">●</td><td>●</td><td>●</td><td>●</td><td></td><td></td></tr>
<tr><td></td><td>Rose</td><td>Rosaceae</td><td></td><td></td><td></td><td></td><td colspan="2">●</td><td></td><td>●</td><td></td><td></td><td></td></tr>
<tr><td></td><td>Queen</td><td>Filipendula</td><td></td><td></td><td></td><td>●</td><td colspan="2">●</td><td></td><td></td><td></td><td></td><td></td></tr>
<tr><td></td><td>Dock</td><td>Rumex *</td><td></td><td></td><td></td><td></td><td colspan="2"></td><td></td><td></td><td></td><td></td><td></td></tr>
<tr><td></td><td>Polypody</td><td>Polypodium</td><td>●</td><td>●</td><td>●</td><td>●</td><td colspan="2"></td><td>●</td><td>●</td><td>●</td><td></td><td>●</td></tr>
</table>

| | | | | | | | | | | | | |
|---|---|---|---|---|---|---|---|---|---|---|---|---|
| | Club spikemoss | *Selaginella selaginoides* | ● | | ● | | | | | | ● | |
| | Sphagnum moss | *Sphagnum* | ● | ● | ● | | | | | | | |
| | | *Cenococcum geophilum* | ● | ● | | ● | | | ● | ● | ● | |
| Mammals | Seal | Phocidae | | | ● | ● | ● | ● | ● | ● | | ● |
| | Ringed seal | *Phoca hispida* | | | ● | | ● | ● | | ● | | |
| | Harp seal | *Phoca groenlandica* | | | ● | | | ● | | ● | | |
| | Common seal | *Phoca vitulina* | | | | | ● | ● | ● | | | ● |
| | Atlantic seal | *Halichoerus grypus* | | | | | | ● | | ● | ● | |
| | Bartrobbe | *Erignathus barbatus* | | | | | | | | ● | | |
| | Cetacean | Cetacea | | | | | | ● | | | | |
| | Rodent | Rodentia | | | | | | | | ● | | |
| Fish | Atlantic cod | *Gadus morthua* | | | ● | ● | ● | ● | ● | ● | | |
| | American pollack | *Pollachius virens* | | | | ● | ● | ● | | ● | ● | |
| | Cornish salmon | *Merluccius merluccius* | | | | | ● | | | | | |
| | Callagh | *Pollachius pollachius* | | | | | | ● | | | | |
| | Common ling | *Molva molva* | | | | | | | | ● | | |
| | Arbroath smokie | *Melanogrammus aeglefinus* | | | | | | | | ● | | |
| Shellfish | | *Littorina* | | | ● | ● | | | | | ● | |
| | Common periwinkle | *Littorina littorea* | | ● | | ● | ● | | ● | ● | ● | |
| | Bay mussel | *Mytilus edulis* | | | ● | ● | ● | | ● | | ● | |
| | Iceland cyprina | *Arctica islandica* | | | | | ● | | ● | ● | | |
| | Sea urchin | *Echinus* | | | ● | | ● | | ● | ● | ● | ● |
| Birds | Common Murre | *Uria aalge* | | | ● | ● | ● | ● | | ● | ● | ● |
| | Great auk | *Pinguinus impennis* | | | | | ● | | ● | ● | | |
| | Atlantic puffin | *Fratercula arctica* | | | | | | | | ● | | |
| | Grouse | Phasianidae | | | | | ● | | | ● | | ● |
| | Duck | Anatidae | | | | | ● | ● | | ● | | |
| | Common snipe | *Gallinago gallinago* | | | | | | ● | | | | |
| | Black-legged kittiwake | *Rissa tridactyla* | | | | | | ● | | | | |
| | European herring gull | *Larus argentatus* | | | | | | | | ● | | |
| | Gannet | *Morus bassanus* | | | | | | | | ● | | |

S3. Descriptions of stratigraphy and chronology at Baŋkgohppi (Martens et al. 2017; calibration: OxCal v4.4.4 (Bronk Ramsey 2009); r:5: atmospheric data (Reimer et al. 2020)).

| Layer | Description | $^{14}$C Age (BP) | Calibrated Age (cal. BCE) (95.4% probability) | | | Percentage of Contents | | | |
|---|---|---|---|---|---|---|---|---|---|
| | | | | | | Botanical | Mineral | Zoological | Objects |
| 1 | Top soil, forest turf | - | - | - | - | - | - | - | - |
| 2 | Midden, part of the manure, gray-brown cultural layer | - | - | - | - | 20 | 75 | 3 | 2 |
| 3 | Midden, black-brown cultural layer in the manure | - | - | - | - | 80 | 10 | 10 | - |
| 4 | Midden, gray-brown culture layer, bones and shells well-preserved due to high lime content of the manure | 3630±30 | 2130 (10.9%) 2089 | 2045 (84.5%) 1896 | - | 30 | 60 | 10 | - |
| 5 | Midden, most marked layer in the manure, outside the wall embankment. | 3740±30 | 2277 (4.7%) 2255 | 2207 (90.7%) 2035 | - | 30 | 55 | 14 | 1 |
| 6 | Sand lens in layer 5 | 3850±30 | 2456 (72.9%) 2270 | 2260 (22.6%) 2204 | - | 20 | 70 | 10 | - |
| 7 | Part of the wall of the house | 3760±30 | 2287 (15.4%) 2246 | 2238 (65.2%) 2127 | 2092 (14.9%) 2041 | 10 | 59 | 30 | 1 |
| 8 | Subsoil, beach wall, sand, medium brown layer, part of the beach embankment under the house and the manure | 3830±30 | 2453 (4.6%) 2419 | 2408 (7.2%) 2374 | 2355 (80.6%) 2197 | 20 | 75 | 5 | - |

S4. A plan of Finnes site (modified from Wickler 2016, 2026).

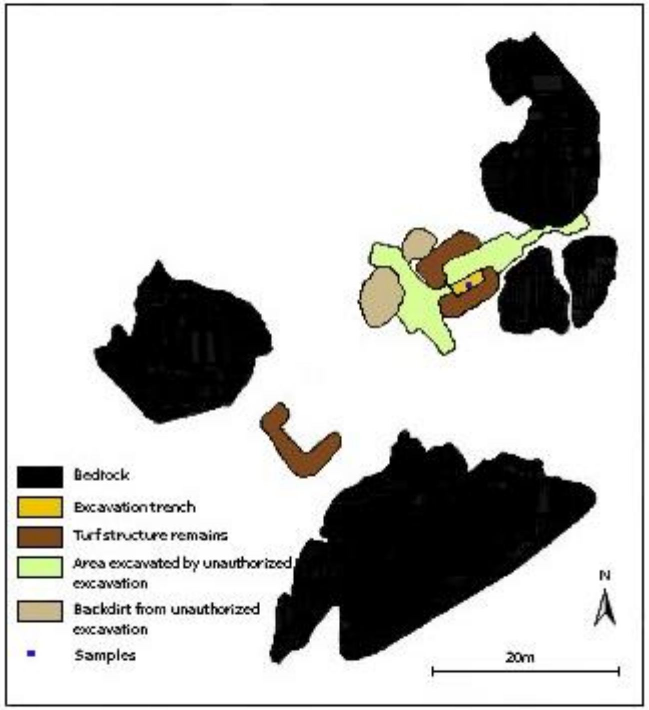

S5. A table of the original stratigraphy (Wickler 2016), the updated stratigraphy, and chronology (calibration: OxCal v4.4.4 (Bronk Ramsey 2009, 2021); r:5: atmospheric data (Reimer et al. 2020)).

| Original layer | Revised layer | Context | $^{14}$C age (BP) | Calibrated age (95.4% probability) | |
|---|---|---|---|---|---|
| 1A | 1 | Turf | - | - | - |
| 1B | 2 | Disturbed cultural deposit representing late medieval and post-medieval settlement mound site | - | - | - |
| 3 | 3 | Truncated late medieval cultural deposit (disturbed by layer 2) 1299-1400 cal. CE | 616±27 | 1299 (95.4%)<br>1400 cal. CE | |
| 4A<br>4B | 4A<br>4B | Upper (4A) and lower (4B) high medieval shell midden deposit dominated by horse mussel with remains of a stone wall (shells more heavily fragmented and slightly darker color in 4B) 1279-1321 cal. CE | 669±28<br>667±27 | 1279 (52.6%)<br>1321 cal. CE<br>1280 (51.3%)<br>1321 cal. CE | 1358 (44.1%)<br>1390 cal. CE<br>1358 (44.1%)<br>1390 cal. CE |
| 7 | 5 | Upper Early Metal Age cultural deposit representing seasonal marine exploitation site<br>ca. 400-200 cal. BCE | 2275±30<br>2243±29<br>2237±29 | 400 (47.5%)<br>351 cal. BCE<br>390 (27.1%)<br>346 cal. BCE<br>389 (24.8%)<br>343 cal. BCE | 302 (48.0%)<br>208 cal. BCE<br>317 (68.4%)<br>204 cal. BCE<br>320 (70.6%)<br>202 cal. BCE |
| 8 | 6 | Lower Early Metal Age cultural deposit representing seasonal marine exploitation site | - | - | - |

S6 List of macro- and microfossils at Finnes.

a. The list of fauna excavated at the Finnes site in 2011 (Wickler 2016, 2026; Calibration: Bronk Ramsey 2009, 2021; Reimer et al. 2020).

| Category | Common name | Scientific name | Layer | | Radiocarbon date (2011) |
|---|---|---|---|---|---|
| | | | 2011 | 2020 | |
| Mammals | Common seal | *Phoca vitulina* | 7 & 6 | 5 | Layer 6 1277-1390 (cal. CE) |
| | Atlantic/Gray seal | *Halichoerus grypus* | | | |
| Birds | European/Common shag | *Phalacrocorax aristotelis* | | | |
| | Black-legged Kittiwake | *Rissa tridactyla* | | | |
| | (Northern) fulmar | *Fulmarus glacialis* | | | Layer 7 400-208 (cal. BCE) |
| Fish | Atlantic cod | *Gadus morhua* | | | |
| | Atlantic halibut | *Hippoglossus hippoglossus* | | | |
| | Brismak/Tusk | *Brosme brosme* | | | |
| | Common ling | *Molva molva* | | | |
| Shellfish | Northern horse mussel | *Modiolus modiolus* | 4A | 4A | |

b. Detail of fauna excavated in 2011 and number of identified specimens at Finnes. Revised layer designations are in parentheses (see Figure 1 and S5).

| Category | Taxa | Norwegian name | Layer: 8 (7) | 10(4) | 12(6) | Grand Total |
|---|---|---|---|---|---|---|
| Amphibian | Amphibia | Amphibian | | 2 | | 2 |
| | Amphibia Total | | | 2 | | 2 |
| Amphibian Total | | | | 2 | | 2 |
| Bird | Alcidae | Auk | 1 | | | 1 |
| | Alcidae Total | | 1 | | | 1 |
| | Anseriformes | Waterfowl | 1 | | | 1 |
| | Anseriformes Total | | 1 | | | 1 |
| | Anser sp | Bean Geese | | 1 | | 1 |
| | Anser sp Total | | | 1 | | 1 |
| | Cepphus grylle | Black Guillemot | 1 | 1 | | 2 |
| | Cepphus grylle Total | | 1 | 1 | | 2 |
| | Charadriiformes | Alcid | | 3 | | 3 |
| | Charadriiformes Total | | | 3 | | 3 |
| | Corvus corax | Common Raven | 1 | | | 1 |
| | Corvus corax Total | | 1 | | | 1 |
| | Fulmarus glacialis | Fulmar | 18 | | | 18 |
| | Fulmarus glacialis Total | | 18 | | | 18 |
| | Laridae | Gull | | 1 | 2 | 3 |
| | Laridae Total | | | 1 | 2 | 3 |
| | Larus argentatus | European Herring Gull | 2 | 2 | | 4 |

| | | | | | | |
|---|---|---|---|---|---|---|
| | Larus argentatus Total | | 2 | 2 | | 4 |
| | Larus marinus | Great Black-backed Gull | 1 | | 2 | 3 |
| | Larus marinus Total | | 1 | | 2 | 3 |
| | Larus sp | Gull | 2 | | | 2 |
| | Larus sp Total | | 2 | | | 2 |
| | Mergus serrator | Red-breasted Merganser | | 1 | | 1 |
| | Mergus serrator Total | | | 1 | | 1 |
| | Mergus merganser | Common Merganser | 1 | | | 1 |
| | Mergus merganser Total | | 1 | | | 1 |
| | Morus bassanus | Gannett | | 2 | | 2 |
| | Morus bassanus Total | | | 2 | | 2 |
| | Phalacrocorax aristotelis | European Shag | 79 | 23 | 16 | 118 |
| | Phalacrocorax aristotelis Total | | 79 | 23 | 16 | 118 |
| | Phalacrocorax sp. | Cormorant | 1 | | | 1 |
| | Phalacrocorax sp. Total | | 1 | | | 1 |
| | Rissa tridactyla | Black-legged Kittiwake | 10 | 20 | | 30 |
| | Rissa tridactyla Total | | 10 | 20 | | 30 |
| | Unidentified | Bird | 65 | 19 | 5 | 89 |
| | Unidentified Total | | 65 | 19 | 5 | 89 |
| Bird Total | | | 183 | 73 | 25 | 281 |
| Mammal | Artiodactyla | Artiodactyl | 2 | | | 2 |
| | Artiodactyla Total | | 2 | | | 2 |
| | Halichoerus grypus | Atlantic Seal | 3 | | | 3 |
| | Halichoerus grypus Total | | 3 | | | 3 |
| | Lutra lutra | Common Otter | 1 | 1 | | 2 |
| | Lutra lutra Total | | 1 | 1 | | 2 |
| | Phoca vitulina | Common Seal | 23 | 27 | | 50 |
| | Phoca vitulina Total | | 23 | 27 | | 50 |
| | Phoca vitulina (?) | Common Seal | | 1 | | 1 |
| | Phoca vitulina (?) Total | | | 1 | | 1 |
| | Phocidae | Earless seal | 12 | 2 | | 14 |
| | Phocidae Total | | 12 | 2 | | 14 |
| | Rangifer tarandus | Reindeer | 4 | | | 4 |
| | Rangifer tarandus Total | | 4 | | | 4 |
| | Unidentified | Mammal | 62 | | | 62 |
| | Unidentified Total | | 62 | | | 62 |
| Mammal Total | | | 107 | 31 | | 138 |
| Fish | Brosme brosme | Cusk | 36 | 44 | 7 | 87 |
| | Brosme brosme Total | | 36 | 44 | 7 | 87 |
| | Gadidae | Codfish | 37 | 10 | 4 | 51 |
| | Gadidae Total | | 37 | 10 | 4 | 51 |

| | | | | | | |
|---|---|---|---|---|---|---|
| | Gadus morhua | Cod | 113 | 54 | 41 | 208 |
| | Gadus morhua Total | | 113 | 54 | 41 | 208 |
| | Pleuronectiformes | Flatfish | 2 | 4 | | 6 |
| | Pleuronectiformes Total | | 2 | 4 | | 6 |
| | Hippoglossus hippoglossus | Atlantic Halibut | 22 | 57 | 24 | 103 |
| | Hippoglossus hippoglossus Total | | 22 | 57 | 24 | 103 |
| | Molva molva | Common ling | | | 14 | 14 |
| | Molva molva Total | | | | 14 | 14 |
| | Pollachius sp | Pollock | 8 | | | 8 |
| | Pollachius sp Total | | 8 | | | 8 |
| | Unidentified | Fish | 451 | 549 | 269 | 1269 |
| | Unidentified Total | | 451 | 549 | 269 | 1269 |
| Fish Total | | | 669 | 718 | 359 | 1746 |
| Grand Total | | | 959 | 824 | 384 | 2167 |

c. Identification of shellfish from bulk samples of layer 4A and 4B (samples take in 2020).

| Type | Sample | Taxa identified |
|---|---|---|
| DNA | IG_4A_51 | Buccinum undatum<br>Littorina obtusata |
| DNA | IG_4A_54 | Littorina obtusata<br>Erginus rubellus |
| DNA | IG_4A_57 | Littorina saxatilis<br>Steromphala cineraria<br>Gastropod |
| Bulk | Ingøy_4A | Boreotrophon clathratus<br>Littorina saxatilis<br>Nucella lapillus<br>Hiatella arctica<br>Red seaweed unid.<br>Ostrea edulis<br>Cirripiola<br>Erginus rubellus<br>Buccinum undatum<br>Littorina obtusata<br>Modiolus modiolus |
| Bulk | Ingøy_4B | Nucella lapillus<br>Boreotrophon clathratus<br>Steromphala cineraria<br>Astarte borealis<br>Chlamys islandica<br>Hiatella arctica<br>Rhodolith<br>Red seaweed unid.<br>Erginus rubellus<br>Littorina obtusata<br>Buccinum undatum<br>Mytilus edulis |

d. Pollen analysis (samples taken in 2020).

200 of the added Lycopodium spores were counted in each sample and other taxa seen were identified and counted.

| Sample | Layer_Depth | Taxa and counts |
|---|---|---|
| 4 | 1A_11 | 375 of the added Lycopodium spores<br>6 Betula<br>57 Pinus<br>3 Alnus<br>3 Salix<br>54 Ericales<br>36 Empetrum<br>6 Calluna<br>27 Carex<br>18 Poaceae<br>3 Rumex<br>3 Chenopodiaceae<br>12 Hordeum<br>3 Rosaceae<br>6 Filipendula<br>9 Apiaceae<br>6 Caryophyllaceae<br>9 Apiaceae<br>6 Caryophyllaceae<br>30 Asteraceae sect. cichorioideae<br>3 unidentified<br>Charcoal |
| 5 | 1A_18 | 200 of the added Lycopodium spores<br>3 Pinus<br>5 Betula<br>2 Filipendula<br>35 Ericales<br>Asteraceae sect. cichorioideae<br>Carex<br>1 Brassicaceae<br>1 Sordaria |
| 6 | 3_38 | 200 of the added Lycopodium spores<br>3 Pinus<br>2 Ericales<br>Charcoal |
| 7 | 4A_43 | 200 of the added Lycopodium spores<br>16 charcoal particles |
| 8 | 4A_46 | 200 of the added Lycopodium spores<br>3 Pinus<br>1 Filipendula<br>Fungal spores<br>A few Sordaria<br>44 charcoal particles |
| 9 | 4A_51 | 200 of the added Lycopodium spores<br>1 Alnus<br>Fungal spores |

| | | |
|---|---|---|
| | | 128 charcoal particles |
| 10 | 4A_54 | 200 of the added Lycopodium spores<br>3 Pinus<br>Fungal spores (not Sordaria)<br>108 charcoal particles |
| 11 | 4A_57 | 200 of the added Lycopodium spores<br>8 Sordaria<br>20 charcoal particles |
| 12 | 4B_62 | 200 of the added Lycopodium spores<br>1 Pinus<br>1 Filipendula<br>Charcoal |
| 13 | 4B_64 | 200 of the added Lycopodium spores<br>1 Pinus<br>Charcoal |
| 14 | 4B_73 | 200 of the added Lycopodium spores<br>7 Pinus<br>84 charcoal particles |

S7. Percentage of edible plant taxa identified by P6-Loop at Baŋkgohppi.

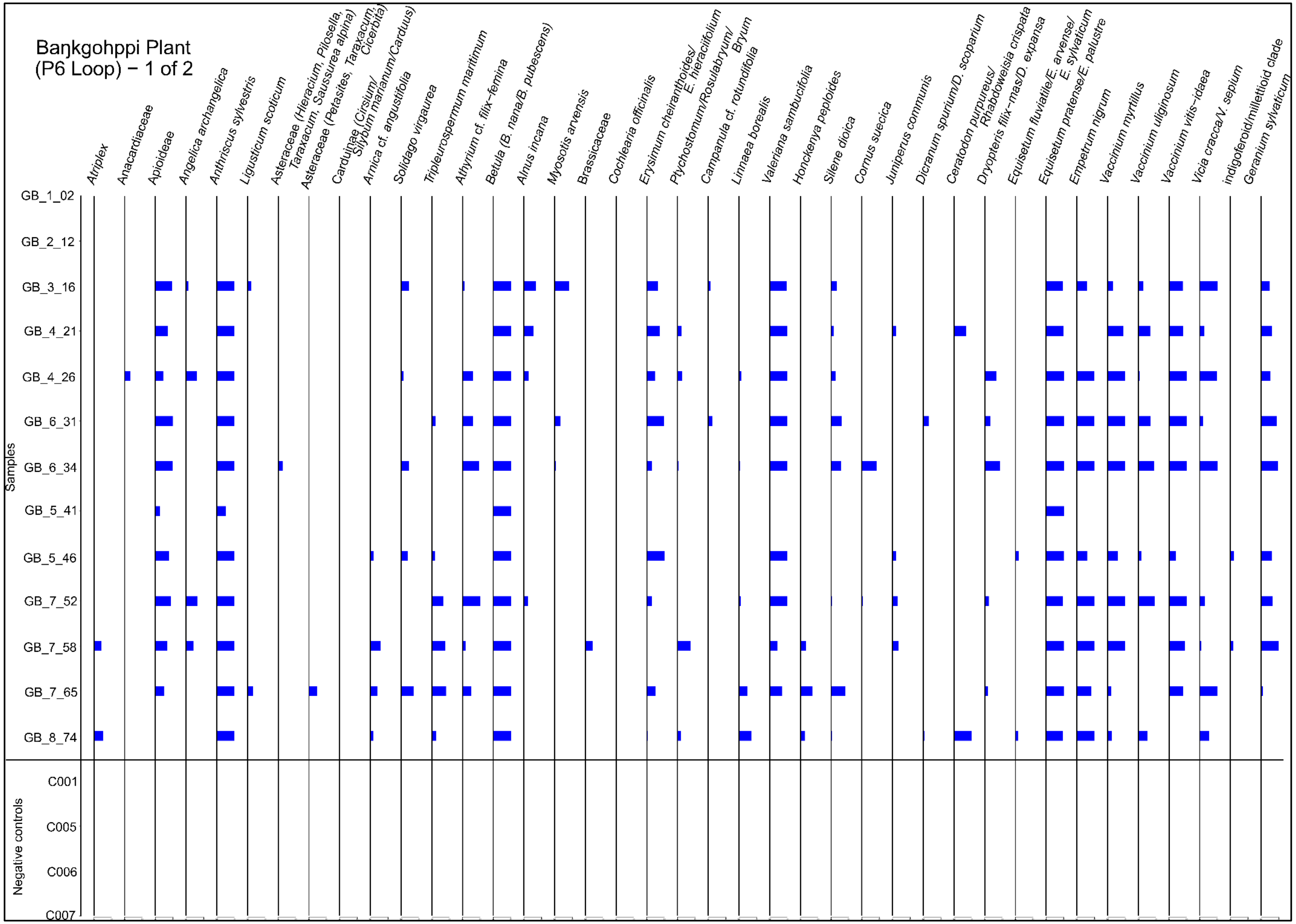

S7. Cont.

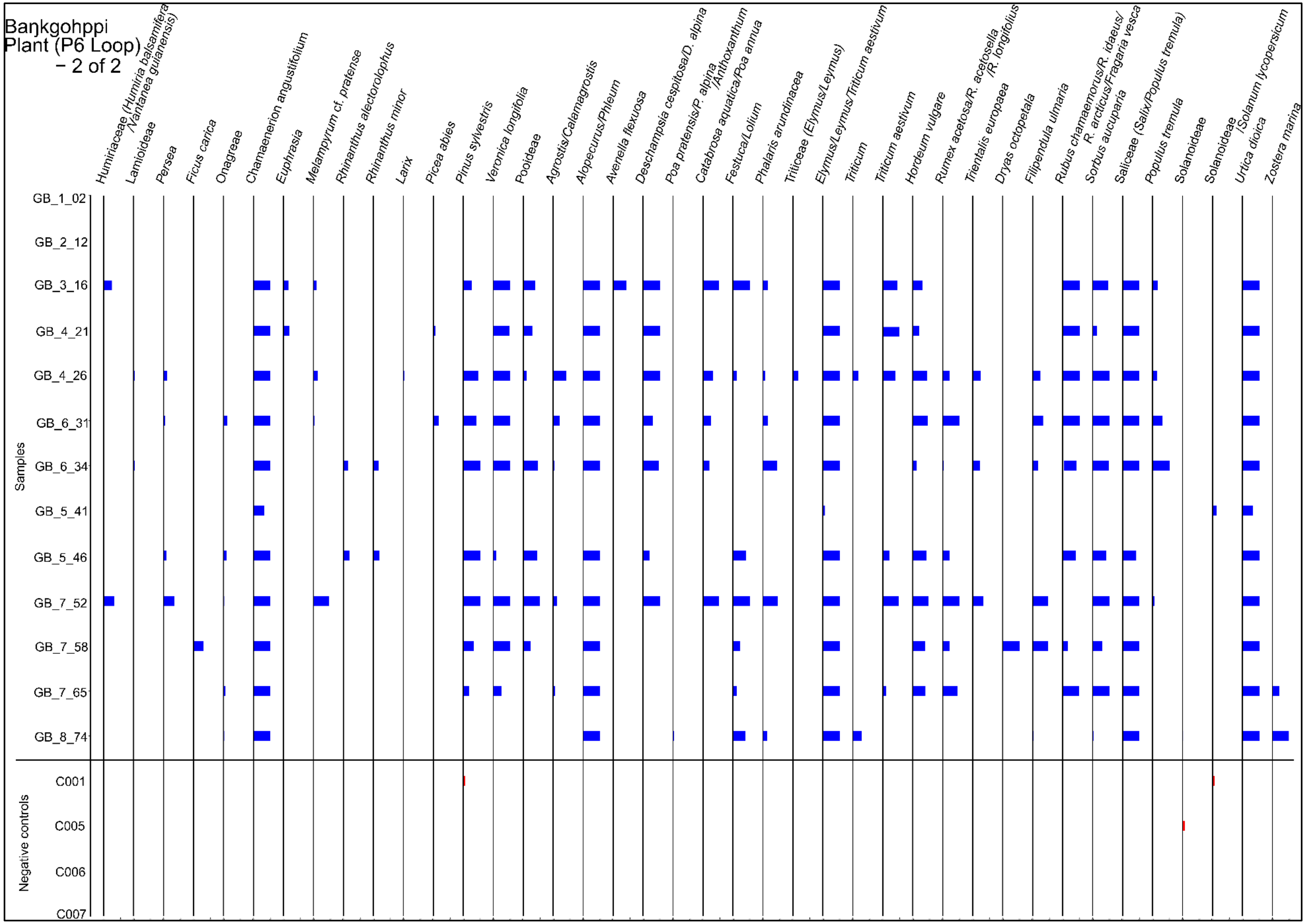
Baŋkgohppi
Plant (P6 Loop)
– 2 of 2
Humiriaceae (Humiria balsamifera /Vantanea guianensis)
Lamioideae
Persea
Ficus carica
Onagreae
Chamaenerion angustifolium
Euphrasia
Melampyrum cf. pratense
Rhinanthus alectorolophus
Rhinanthus minor
Larix
Picea abies
Pinus sylvestris
Veronica longifolia
Pooideae
Agrostis/Calamagrostis
Alopecurus/Phleum
Avenella flexuosa
Deschampsia cespitosa/D. alpina
Poa pratensis/P. alpina /Anthoxanthum
Catabrosa aquatica/Poa annua
Festuca/Lolium
Phalaris arundinacea
Triticeae (Elymus/Leymus)
Elymus/Leymus/Triticum aestivum
Triticum
Triticum aestivum
Hordeum vulgare
Rumex acetosa/R. acetosella /R. longifolius
Trientalis europaea
Dryas octopetala
Filipendula ulmaria
Rubus chamaemorus/R. idaeus/ R. arcticus/Fragaria vesca
Sorbus aucuparia
Saliceae (Salix/Populus tremula)
Populus tremula
Solanoideae
Solanoideae /Solanum lycopersicum
Urtica dioica
Zostera marina
Samples
GB_1_02
GB_2_12
GB_3_16
GB_4_21
GB_4_26
GB_6_31
GB_6_34
GB_5_41
GB_5_46
GB_7_52
GB_7_58
GB_7_65
GB_8_74
Negative controls
C001
C005
C006
C007

S8. Percentage of edible taxa identified at Baŋkgohppi by sedaDNA.

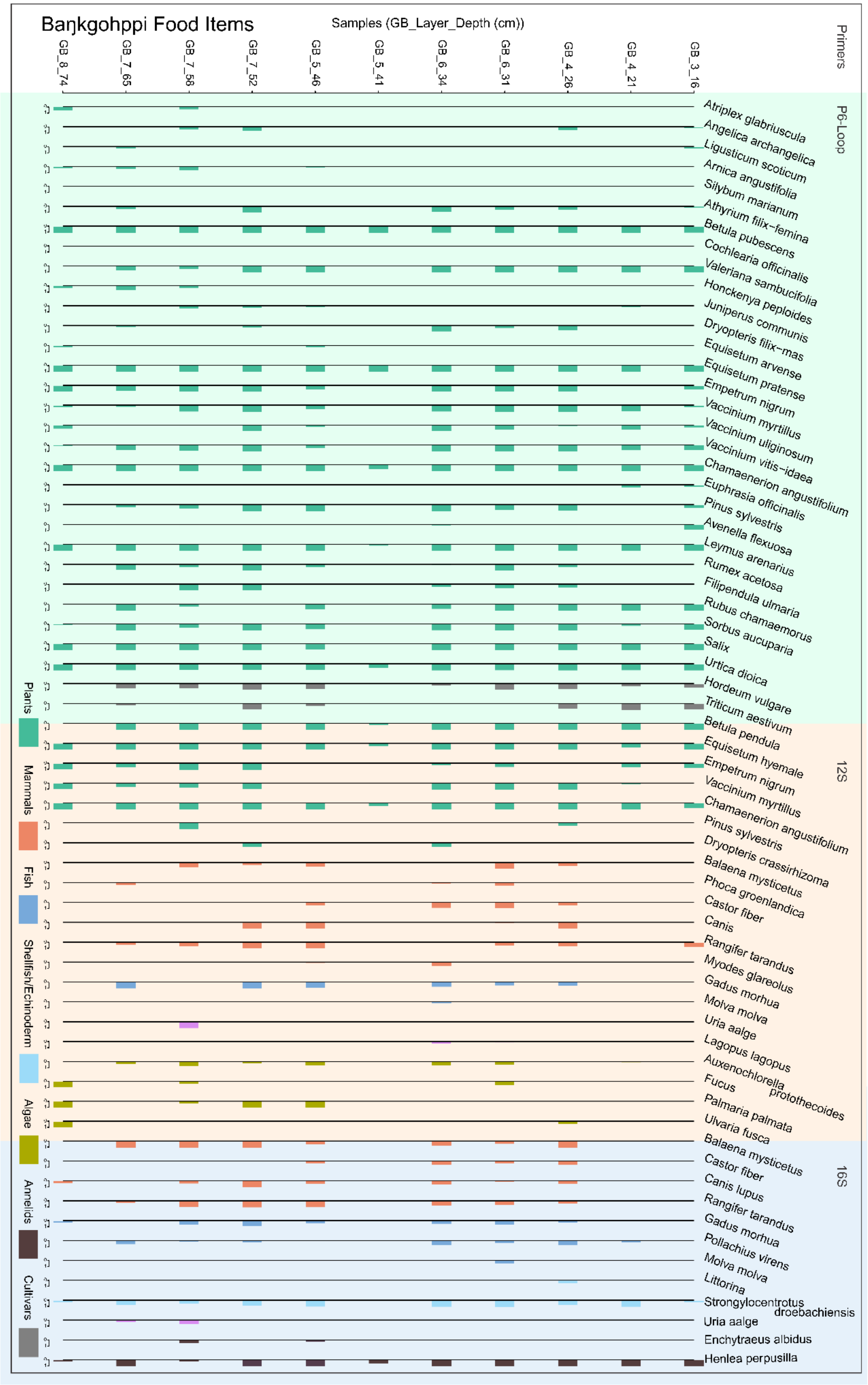

S9. Percentage of edible taxonomic units identified by 12S primer at Baŋkgohppi.

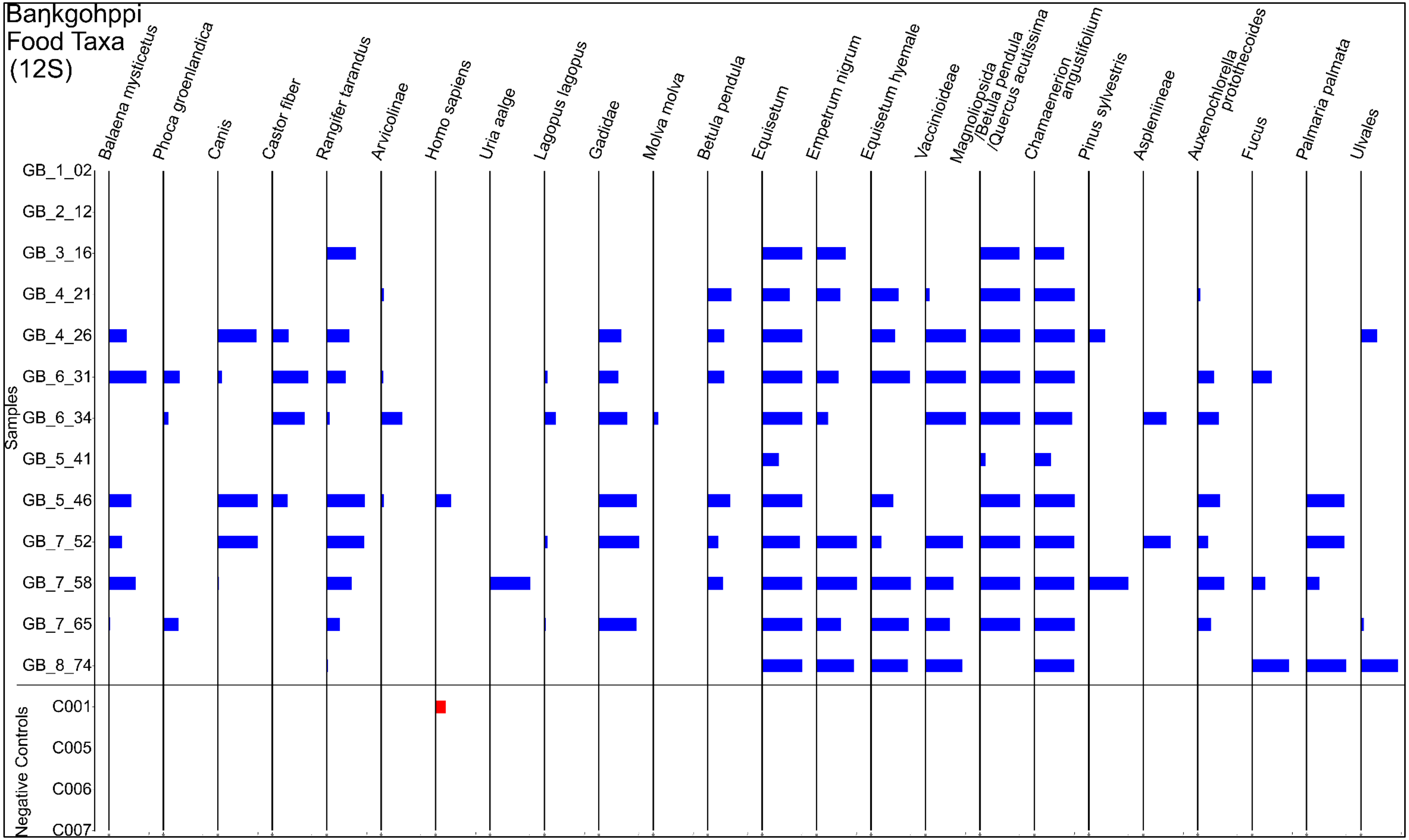

S10. Percentage of edible taxonomic units identified by 16S primer at Baŋkgohppi.

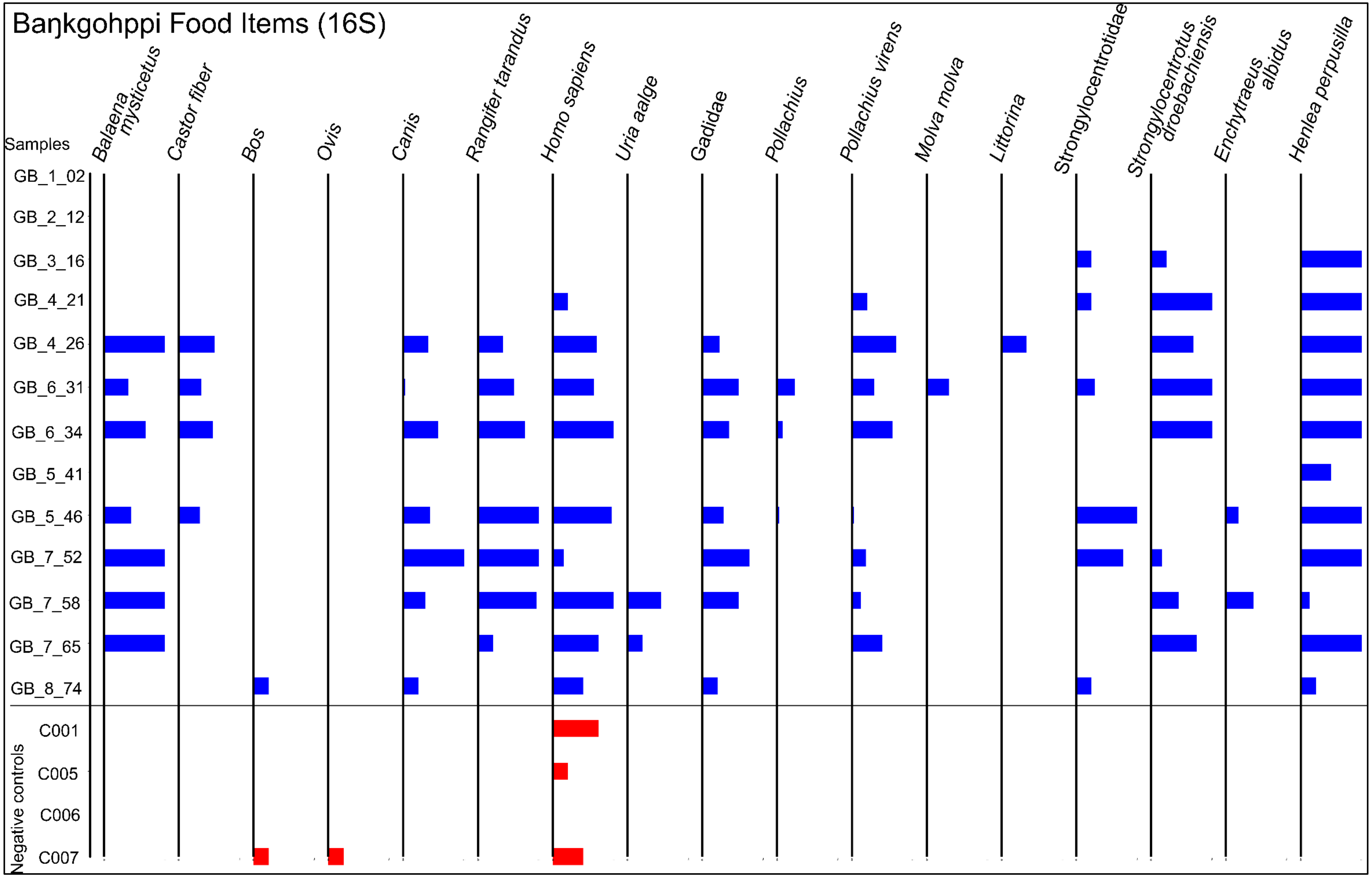

S11. Top 10 plant families and their relative abundance at Baŋkgohppi based on the total reads.

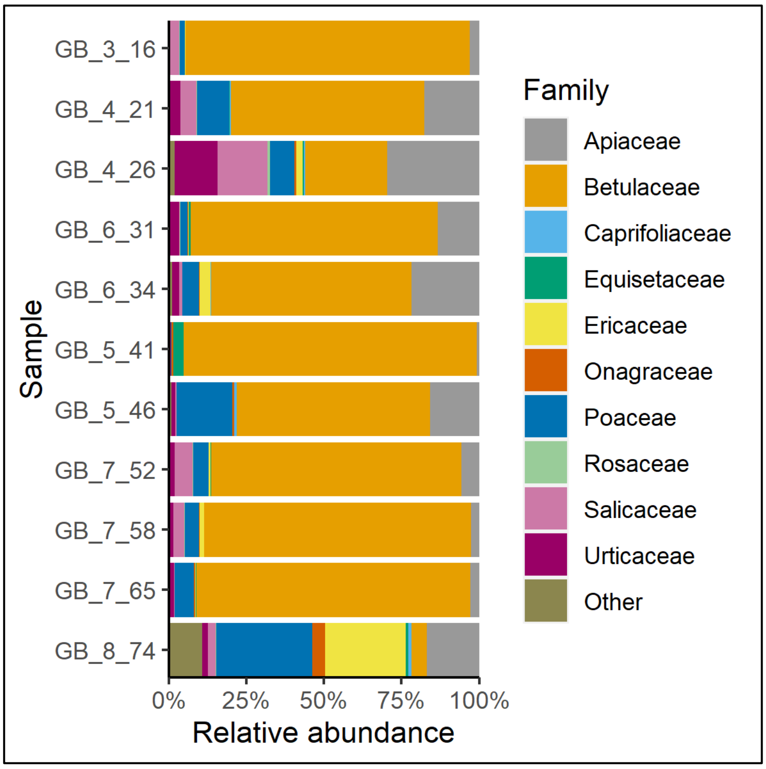

S12. Percentage of edible plant taxa identified by P6-Loop at Finnes.

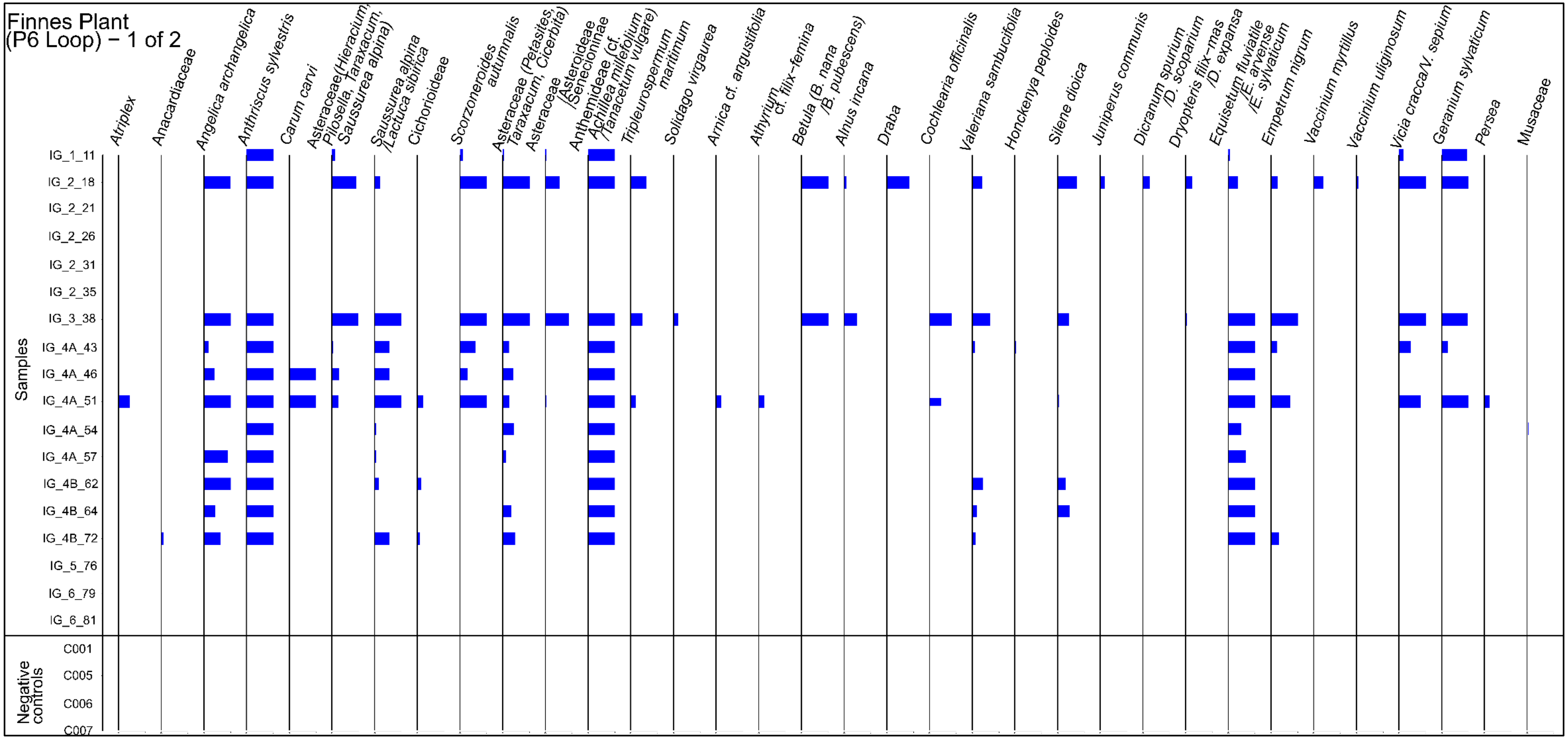

S12. Cont.

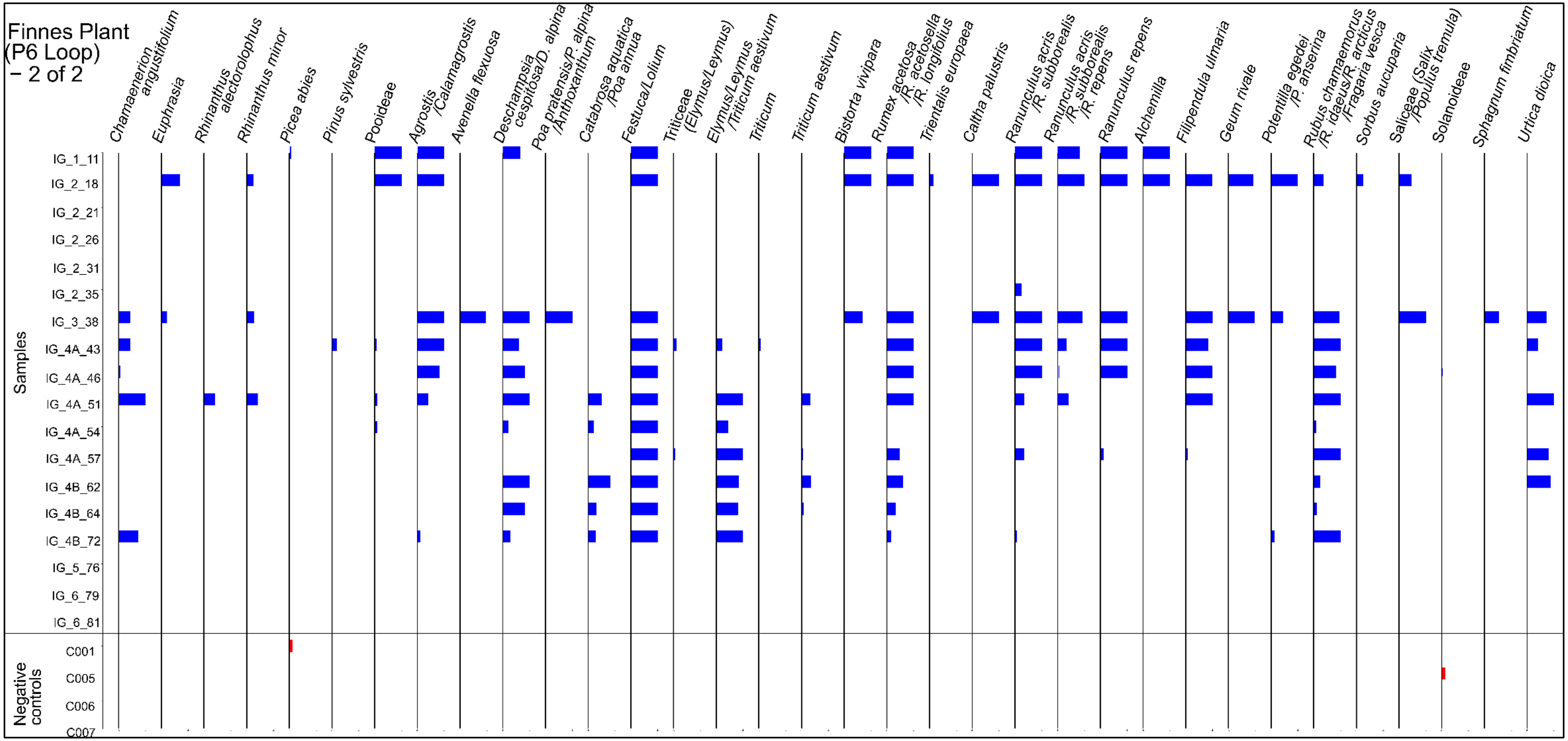
Finnes Plant (P6 Loop) – 2 of 2
Chamaenerion angustifolium
Euphrasia
Rhinanthus alectorolophus
Rhinanthus minor
Picea abies
Pinus sylvestris
Pooideae
Agrostis /Calamagrostis
Avenella flexuosa
Deschampsia cespitosa/D. alpina
Poa pratensis/P. alpina /Anthoxanthum
Catabrosa aquatica /Poa annua
Festuca/Lolium
Triticeae (Elymus/Leymus)
Elymus/Leymus /Triticum aestivum
Triticum
Triticum aestivum
Bistorta vivipara
Rumex acetosa /R. acetosella /R. longifolius
Trientalis europaea
Caltha palustris
Ranunculus acris /R. subborealis
Ranunculus acris /R. subborealis /R. repens
Ranunculus repens
Alchemilla
Filipendula ulmaria
Geum rivale
Potentilla egedei /P. anserina
Rubus chamaemorus /R. idaeus/R. arcticus /Fragaria vesca
Sorbus aucuparia
Saliceae (Salix /Populus tremula)
Solanoideae
Sphagnum fimbriatum
Urtica dioica
Samples
IG_1_11
IG_2_18
IG_2_21
IG_2_26
IG_2_31
IG_2_35
IG_3_38
IG_4A_43
IG_4A_46
IG_4A_51
IG_4A_54
IG_4A_57
IG_4B_62
IG_4B_64
IG_4B_72
IG_5_76
IG_6_79
IG_6_81
Negative controls
C001
C005
C006
C007

S13. Percentage of edible taxonomic units identified by 12S primer at Finnes.

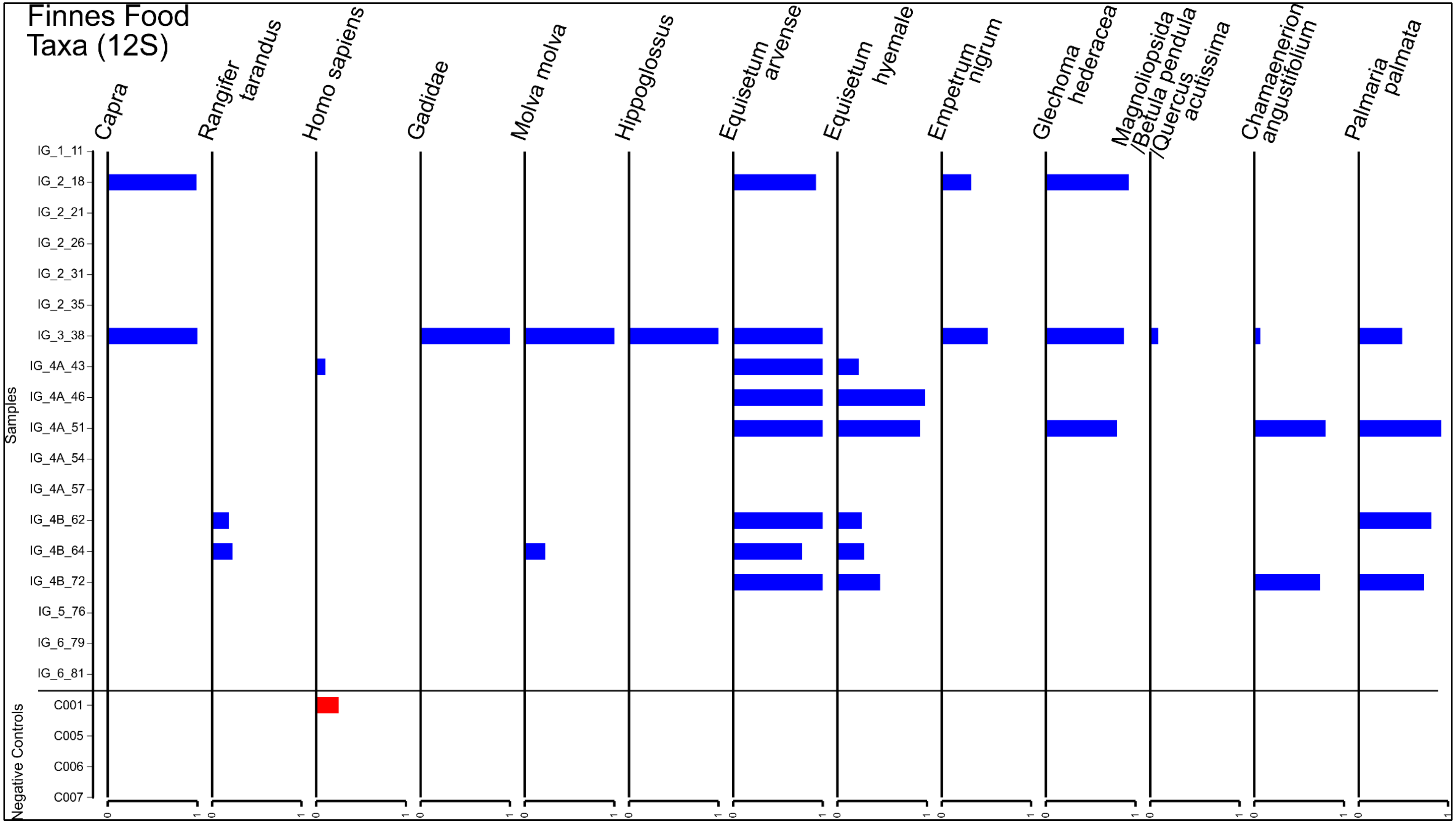

S14. Percentage of edible taxonomic units identified by 16S primer at Finnes.

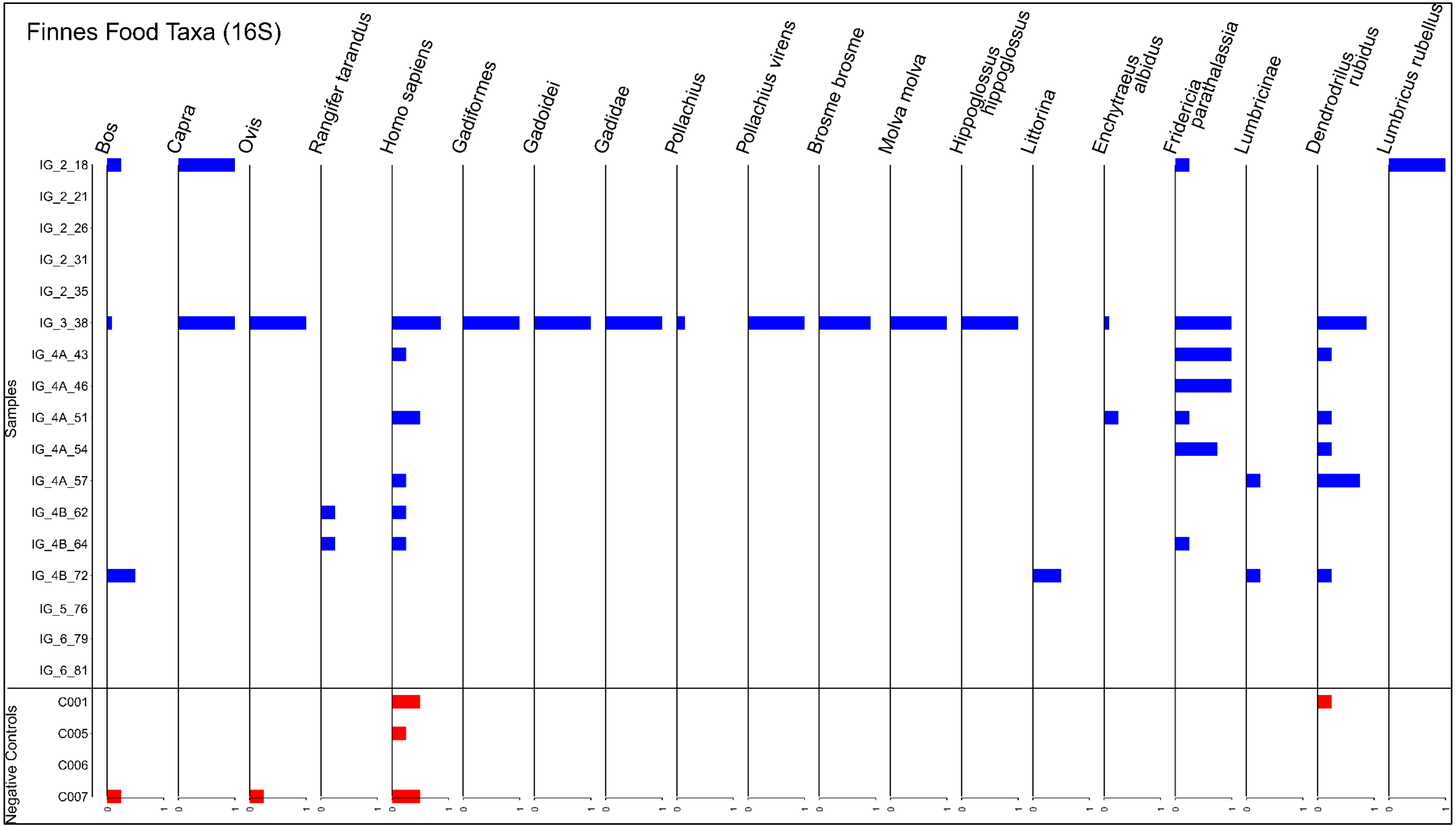

S15. Percentage of edible taxa identified at Finnes by sedaDNA.

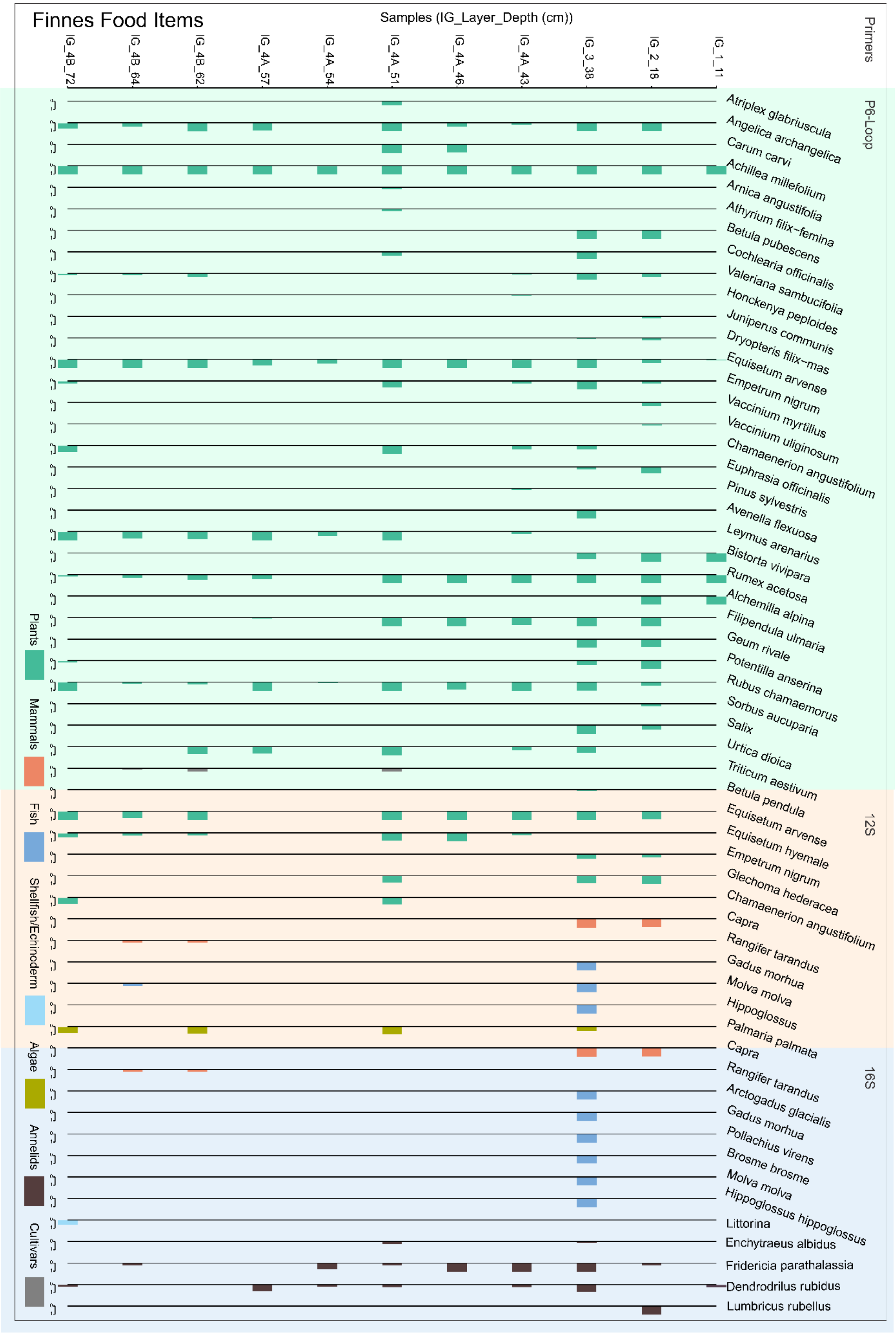

S16. Top 10 plant families and their relative abundance at Finnes based on the total reads of metabarcoding data.

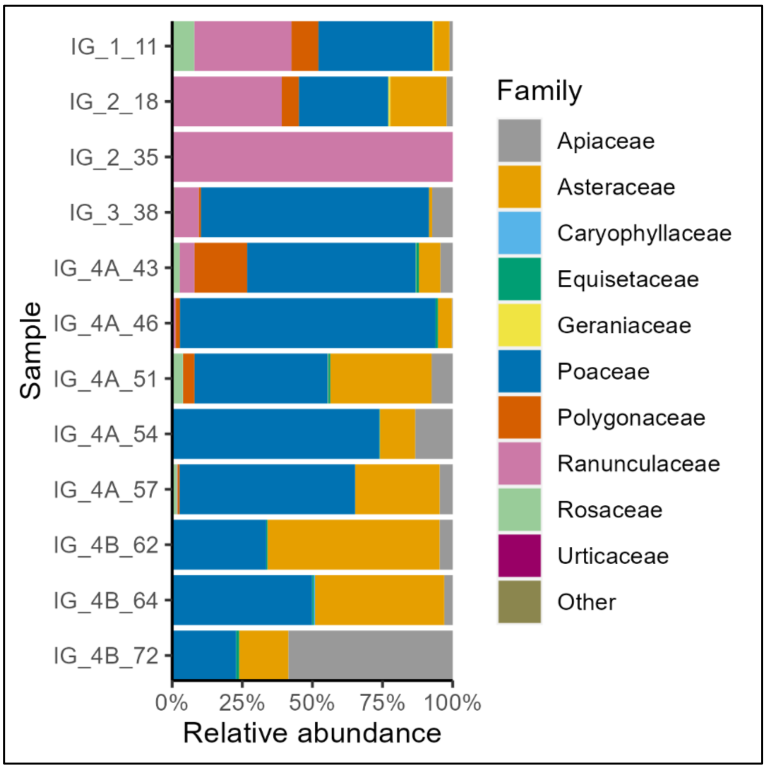

S17. Table of retained taxa identified through shotgun sequencing demonstrating a damage pattern.

| count | avg.length | sample | uniq_kmer_prop | uniq_kmer | tot_kmer | conf | param | paramstd | pvalue | RSME | dam_mag | family | genus | family_prop | taxa_name |
|---|---|---|---|---|---|---|---|---|---|---|---|---|---|---|---|
| 109330 | 76.3821001 | IG_4B_72 | 0.46925145 | 2379553 | 5070955 | 1 | 0.61 | 0.059 | 0 | 0.001 | 0.02691667 | Poaceae | NA | 0.65478044 | Poaceae |
| 38400 | 85.6707292 | IG_4B_72 | 0.53611451 | 1146082 | 2137756 | 1 | 0.854 | 0.068 | 0 | 0.001 | 0.02366667 | Asteraceae | Achillea | 0.96898736 | Achillea |
| 35883 | 82.0452861 | IG_4B_72 | 0.62887937 | 1174458 | 1867541 | 1 | 0.708 | 0.069 | 0 | 0.002 | 0.03541667 | Poaceae | Leymus | 0.2149043 | Leymus |
| 23026 | 82.3670199 | IG_4B_72 | 0.645666 | 778546 | 1205803 | 1 | 0.749 | 0.129 | 0 | 0.003 | 0.02158333 | Rosaceae | Rubus | 0.96508655 | Rubus |
| 11613 | 83.7730991 | IG_4B_72 | 0.66189887 | 413334 | 624467 | 1 | 0.751 | 0.108 | 0 | 0.002 | 0.02508333 | Poaceae | Festuca | 0.06955058 | Festuca |
| 5833 | 85.9691411 | IG_4B_72 | 0.6559479 | 214146 | 326468 | 1 | 0.516 | 0.134 | 0 | 0.004 | 0.03766667 | Poaceae | Elymus | 0.034934 | Elymus |
| 2835 | 74.6582011 | IG_4B_72 | 0.68059176 | 86167 | 126606 | 1 | 0.69 | 0.119 | 0 | 0.006 | 0.05125 | Poaceae | Hordelymus | 0.01697889 | Hordelymus |
| 196919 | 83.39695 | GB_5_46 | 0.65687387 | 6906946 | 10514874 | 1 | 0.717 | 0.041 | 0 | 0.001 | 0.03125 | Betulaceae | Betula | 0.94657578 | Betula |
| 173550 | 75.286949 | GB_5_46 | 0.47659014 | 3745784 | 7859550 | 1 | 0.7 | 0.054 | 0 | 0.001 | 0.02816667 | Poaceae | NA | 0.54608786 | Poaceae |
| 103584 | 72.4740307 | GB_5_46 | 0.53384739 | 2348731 | 4399630 | 1 | 0.673 | 0.054 | 0 | 0.002 | 0.03883333 | Caprifoliaceae | Valeriana | 1 | Valeriana |
| 59021 | 82.108724 | GB_5_46 | 0.62398029 | 1919057 | 3075509 | 1 | 0.576 | 0.058 | 0 | 0.002 | 0.03875 | Poaceae | Elymus | 0.18571393 | Elymus |
| 44601 | 77.607094 | GB_5_46 | 0.51085185 | 1084704 | 2123324 | 1 | 0.795 | 0.05 | 0 | 0.002 | 0.03108333 | Apiaceae | NA | 0.78888162 | Apiaceae |
| 39633 | 82.8220422 | GB_5_46 | 0.64890594 | 1358482 | 2093496 | 1 | 0.75 | 0.068 | 0 | 0.002 | 0.03766667 | Poaceae | Leymus | 0.12470816 | Leymus |
| 21316 | 78.0424095 | GB_5_46 | 0.69241777 | 728231 | 1051722 | 1 | 0.766 | 0.037 | 0 | 0.001 | 0.03683333 | Poaceae | Poa | 0.06707236 | Poa |
| 18258 | 78.1391719 | GB_5_46 | 0.64261137 | 568116 | 884074 | 1 | 0.753 | 0.095 | 0 | 0.003 | 0.03566667 | Poaceae | Alopecurus | 0.05745014 | Alopecurus |
| 13878 | 80.3463756 | GB_5_46 | 0.64179978 | 448430 | 698707 | 1 | 0.797 | 0.112 | 0 | 0.003 | 0.03225 | Urticaceae | Urtica | 1 | Urtica |
| 10918 | 76.9020883 | GB_5_46 | 0.68722868 | 351914 | 512077 | 1 | 0.889 | 0.083 | 0 | 0.003 | 0.03141667 | Canidae | NA | 0.54028108 | Canidae |
| 9290 | 80.4231432 | GB_5_46 | 0.69845506 | 327178 | 468431 | 1 | 0.673 | 0.055 | 0 | 0.004 | 0.07341667 | Canidae | Canis | 0.45971892 | Canis |
| 7841 | 71.3609234 | GB_5_46 | 0.65533084 | 212531 | 324311 | 1 | 0.904 | 0.075 | 0 | 0.004 | 0.04491667 | Onagraceae | Chamaenerion | 0.90542725 | Chamaenerion |
| 7033 | 74.6478032 | GB_5_46 | 0.7030645 | 224627 | 319497 | 1 | 0.79 | 0.154 | 0 | 0.003 | 0.022 | Rosaceae | Rubus | 0.81343974 | Rubus |
| 6716 | 83.3878797 | GB_5_46 | 0.68261317 | 244753 | 358553 | 1 | 0.866 | 0.142 | 0 | 0.006 | 0.04933333 | Apiaceae | Anthriscus | 0.11878947 | Anthriscus |
| 6293 | 73.1597013 | GB_5_46 | 0.68863124 | 187035 | 271604 | 1 | 0.589 | 0.268 | 0 | 0.005 | 0.02391667 | Cricetidae | Microtus | 0.84515176 | Microtus |
| 4929 | 85.7630351 | GB_5_46 | 0.68955744 | 189529 | 274856 | 1 | 0.557 | 0.127 | 0 | 0.007 | 0.05016667 | Apiaceae | Carum | 0.08718185 | Carum |
| 2835 | 73.0828924 | GB_5_46 | 0.68446046 | 83600 | 122140 | 1 | 0.966 | 0.201 | 0 | 0.005 | 0.02658333 | Geraniaceae | Geranium | 1 | Geranium |
| 2220 | 73.3711712 | GB_5_46 | 0.70137302 | 67531 | 96284 | 1 | 0.687 | 0.482 | 0.008 | 0.009 | 0.03466667 | Pinaceae | Pinus | 1 | Pinus |

| | | | | | | | | | | | | | | | |
|---|---|---|---|---|---|---|---|---|---|---|---|---|---|---|---|
| 1908 | 76.1284067 | GB_5_46 | 0.69349982 | 61037 | 88013 | 1 | 0.795 | 0.293 | 0.001 | 0.008 | 0.03741667 | Gadidae | Gadus | 1 | Gadus |
| 1206 | 67.8897181 | GB_5_46 | 0.70898348 | 32397 | 45695 | 1 | 0.759 | 0.179 | 0 | 0.006 | 0.028 | Rosaceae | NA | 0.13948647 | Rosaceae |
| 819 | 71.4859585 | GB_5_46 | 0.60590988 | 20587 | 33977 | 1 | 0.99 | 0.24 | 0 | 0.009 | 0.0255 | Onagraceae | NA | 0.09457275 | Onagraceae |
| 407 | 72.9041769 | GB_5_46 | 0.69356317 | 12111 | 17462 | 0.994 | 0.99 | 0.15 | 0 | 0.008 | 0.03175 | Rosaceae | Sorbus | 0.04707379 | Sorbus |
| 377495 | 85.8297355 | GB_6_31 | 0.63528288 | 13388870 | 21075446 | 1 | 0.732 | 0.029 | 0 | 0.001 | 0.0265 | Betulaceae | Betula | 0.95084746 | Betula |
| 48829 | 77.7066088 | GB_6_31 | 0.50568199 | 1177969 | 2329466 | 1 | 0.697 | 0.053 | 0 | 0.002 | 0.03191667 | Apiaceae | NA | 0.7894873 | Apiaceae |
| 43276 | 74.7398327 | GB_6_31 | 0.59847864 | 1158751 | 1936161 | 1 | 0.706 | 0.049 | 0 | 0.002 | 0.03908333 | Caprifoliaceae | Valeriana | 1 | Valeriana |
| 26790 | 74.2377006 | GB_6_31 | 0.58981224 | 699003 | 1185128 | 1 | 0.646 | 0.049 | 0 | 0.002 | 0.03183333 | Poaceae | NA | 0.48045194 | Poaceae |
| 22471 | 80.8242624 | GB_6_31 | 0.65742703 | 750829 | 1142072 | 1 | 0.792 | 0.052 | 0 | 0.002 | 0.0455 | Poaceae | Elymus | 0.40299498 | Elymus |
| 18588 | 78.557564 | GB_6_31 | 0.65220233 | 588670 | 902588 | 1 | 0.777 | 0.049 | 0 | 0.002 | 0.04241667 | Urticaceae | Urtica | 1 | Urtica |
| 18285 | 77.789773 | GB_6_31 | 0.5961336 | 520923 | 873836 | 1 | 0.877 | 0.078 | 0 | 0.001 | 0.02066667 | Betulaceae | NA | 0.04605689 | Betulaceae |
| 14188 | 76.6205244 | GB_6_31 | 0.70056179 | 463388 | 661452 | 1 | 0.729 | 0.048 | 0 | 0.002 | 0.0325 | Rosaceae | Rubus | 0.75871658 | Rubus |
| 7218 | 84.5581879 | GB_6_31 | 0.68567627 | 270020 | 393801 | 1 | 0.762 | 0.106 | 0 | 0.005 | 0.04641667 | Apiaceae | Anthriscus | 0.11670358 | Anthriscus |
| 5879 | 70.2874639 | GB_6_31 | 0.69878826 | 165508 | 236850 | 1 | 0.705 | 0.088 | 0 | 0.004 | 0.04341667 | Ericaceae | Vaccinium | 0.63916069 | Vaccinium |
| 5509 | 87.3283718 | GB_6_31 | 0.69011975 | 217955 | 315822 | 1 | 0.645 | 0.05 | 0 | 0.004 | 0.07083333 | Apiaceae | Carum | 0.08907177 | Carum |
| 3801 | 78.0157853 | GB_6_31 | 0.69158612 | 126220 | 182508 | 1 | 0.752 | 0.094 | 0 | 0.004 | 0.03916667 | Poaceae | Alopecurus | 0.06816714 | Alopecurus |
| 3702 | 72.9397623 | GB_6_31 | 0.67205576 | 106832 | 158963 | 1 | 0.762 | 0.109 | 0 | 0.005 | 0.03866667 | Plantaginaceae | Veronica | 1 | Veronica |
| 3469 | 73.7247045 | GB_6_31 | 0.68391559 | 103737 | 151681 | 1 | 0.533 | 0.163 | 0 | 0.007 | 0.03641667 | Onagraceae | Chamaenerion | 0.91506199 | Chamaenerion |
| 3129 | 71.3876638 | GB_6_31 | 0.6974873 | 90326 | 129502 | 1 | 0.592 | 0.163 | 0 | 0.004 | 0.03225 | Rosaceae | NA | 0.1673262 | Rosaceae |
| 2131 | 70.370718 | GB_6_31 | 0.7007323 | 60284 | 86030 | 1 | 0.99 | 0.183 | 0 | 0.006 | 0.03116667 | Ericaceae | Empetrum | 0.2316808 | Empetrum |
| 2107 | 77.752729 | GB_6_31 | 0.70072057 | 70503 | 100615 | 1 | 0.5 | 0.197 | 0 | 0.007 | 0.02625 | Canidae | NA | 0.52478207 | Canidae |
| 1908 | 80.6477987 | GB_6_31 | 0.70587566 | 68213 | 96636 | 1 | 0.55 | 0.075 | 0 | 0.006 | 0.06366667 | Canidae | Canis | 0.47521793 | Canis |
| 1612 | 73.558933 | GB_6_31 | 0.72519475 | 50921 | 70217 | 1 | 0.703 | 0.162 | 0 | 0.006 | 0.0355 | Salicaceae | Salix | 1 | Salix |
| 1483 | 74.6979096 | GB_6_31 | 0.70641302 | 46826 | 66287 | 1 | 0.706 | 0.132 | 0 | 0.005 | 0.03316667 | Geraniaceae | Geranium | 1 | Geranium |
| 1196 | 78.9013378 | GB_6_31 | 0.67222925 | 39316 | 58486 | 1 | 0.324 | 0.127 | 0 | 0.008 | 0.03883333 | Poaceae | Poa | 0.02144907 | Poa |
| 1129 | 72.8246236 | GB_6_31 | 0.70226892 | 33954 | 48349 | 1 | 0.806 | 0.132 | 0 | 0.006 | 0.04808333 | Polygonaceae | Rumex | 1 | Rumex |
| 1070 | 76.0476636 | GB_6_31 | 0.71015405 | 34990 | 49271 | 1 | 0.515 | 0.111 | 0 | 0.007 | 0.04575 | Rosaceae | Sorbus | 0.05721925 | Sorbus |
| 337 | 78.6735905 | GB_6_31 | 0.71931964 | 11799 | 16403 | 0.999 | 0.799 | 0.057 | 0 | 0.006 | 0.04983333 | Dryopteridaceae | Dryopteris | 1 | Dryopteris |

| | | | | | | | | | | | | | | | |
|---|---|---|---|---|---|---|---|---|---|---|---|---|---|---|---|
| 236 | 61.7372881 | GB_6_31 | 0.68771696 | 5151 | 7490 | 1 | 0.99 | 0.098 | 0 | 0.007 | 0.04575 | Cricetidae | NA | 0.20629371 | Cricetidae |

S18. Cytosine deamination plot of *Canis* reads in GB_5_46.

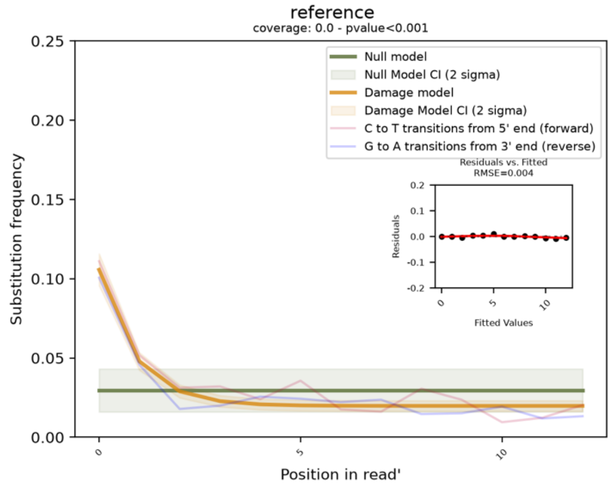

S19. Cytosine deamination plot of *Betula* reads in GB_5_46.

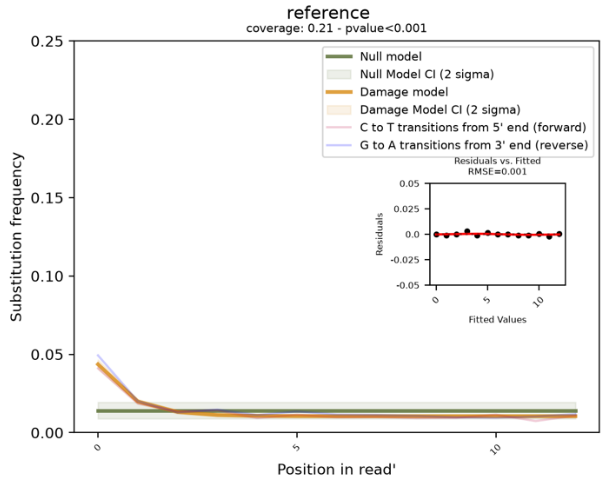

S20. Cytosine deamination plot of *Rubus* reads in GB_6_31.

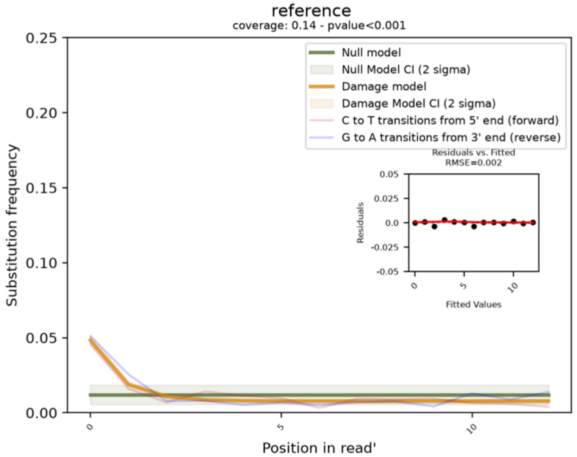

S21. Cytosine deamination plot of *Valeriana* reads in GB_6_31.

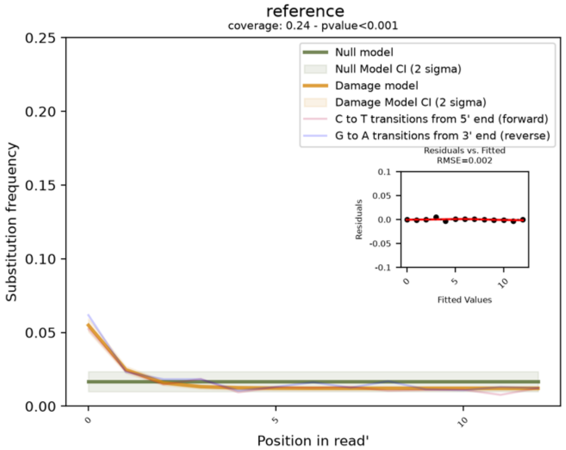

S22. Cytosine deamination plot of *Rubus* reads in IB_4B_72.

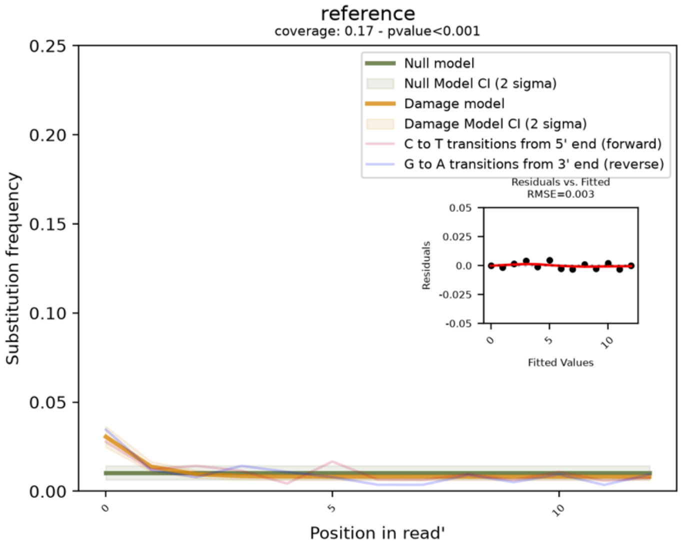

S23. Cytosine deamination plot of *Elymus* reads in IB_4B_72.

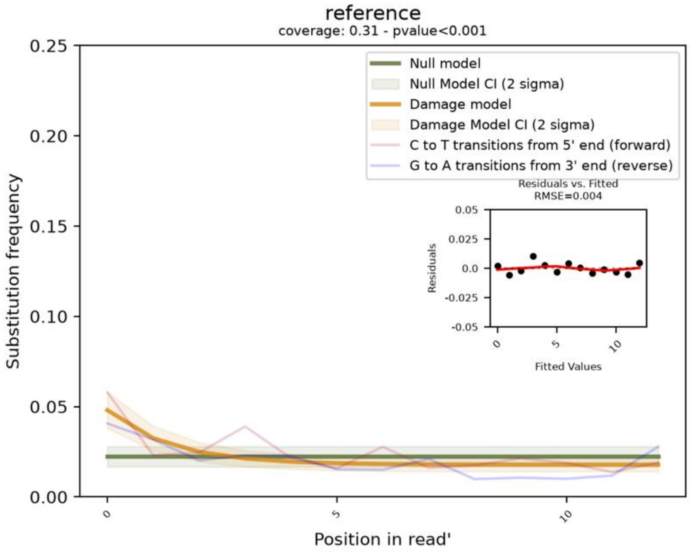

S24. Table of fauna genomes used in shotgun reference database.

| species | taxon_id | ncbi_accession |
|---|---|---|
| Bos taurus | 9913 | GCF_002263795.3 |
| Capra hircus | 9925 | GCA_044587945.1 |
| Capra aegagrus | 9923 | GCA_000978405.1 |
| Ovis aries | 9940 | GCA_033439445.1 |
| Sus scrofa | 9823 | GCF_000003025.6 |
| Equus caballus | 9796 | GCF_041296265.1 |
| Equus asinus | 9793 | GCF_041296235.1 |
| Microtus agrestis | 29092 | GCA_902806775.1 |
| Canis lupus | 9615 | GCA_012044875.1 |
| Felis catus | 9685 | GCF_018350175.1 |
| Rangifer tarandus | 9870 | GCA_949782905.1 |
| Ovibos moschatus | 37176 | GCA_041156055.1 |
| Alces alces | 9852 | GCA_964270355.1 |
| Cervus elephus | 9860 | GCF_910594005.1 |
| Rattus rattus | 10117 | GCF_011064425.1 |
| Megaptera novaeangliae | 9773 | GCA_041834305.1 |
| Phoco vitulina | 9720 | GCF_004348235.1 |
| Phocoena phocoena | 9742 | GCF_963924675.1 |
| Gallus gallus | 9031 | GCF_016699485.2 |
| Bufo bufo | 8384 | GCF_905171765.1 |
| Dicrostonyx torquatus | 85952 | GCA_028555095.1 |
| Lumbirculus variegatus | 61662 | GCA_056247715.1 |
| Lepus timidus | 62621 | GCA_040893245.2 |
| Lagopus muta | 64668 | GCF_023343835.1 |
| Rana temporaria | 8407 | GCF_905171775.1 |
| Anas platyrhynchos | 8839 | GCF_047663525.1 |
| Camelus dromedarius | 9838 | GCF_036321535.1 |
| Vulpes vulpes | 9627 | GCF_048418805.1 |
| Meleagris gallopavo | 9103 | GCA_905368555.5 |
| Homo sapiens | 9606 | GCA_000001405.29 |
| Gadus morhua | 8049 | GCF_902167405.1 |
| Salmo salar | 8030 | GCF_965601325.1 |
| Betta splendens | 158456 | GCF_900634795.4 |
| Lagenorhynchus albirostris | 27610 | GCF_949774975.1 |
| Hippoglossus hippoglossus | 8267 | GCF_009819705.1 |
| Clupea harengus | 7950 | GCA_040183275.2 |
| Caenorhabditis elegans | 6239 | GCF_000002985.6 |
| Drosophila melanogaster | 7227 | GCA_056747195.1 |
| Callosobruchus maculatus | 64391 | GCA_040182625.1 |
| Anolis carolinensis | 28377 | GCF_035594765.1 |
| Lamna ditropis | 38578 | KF962053 |
| Xenopus laevis | 8355 | GCF_017654675.1 |

S25. Total reads and total repeats of *Triticum* at Baŋkgohppi.

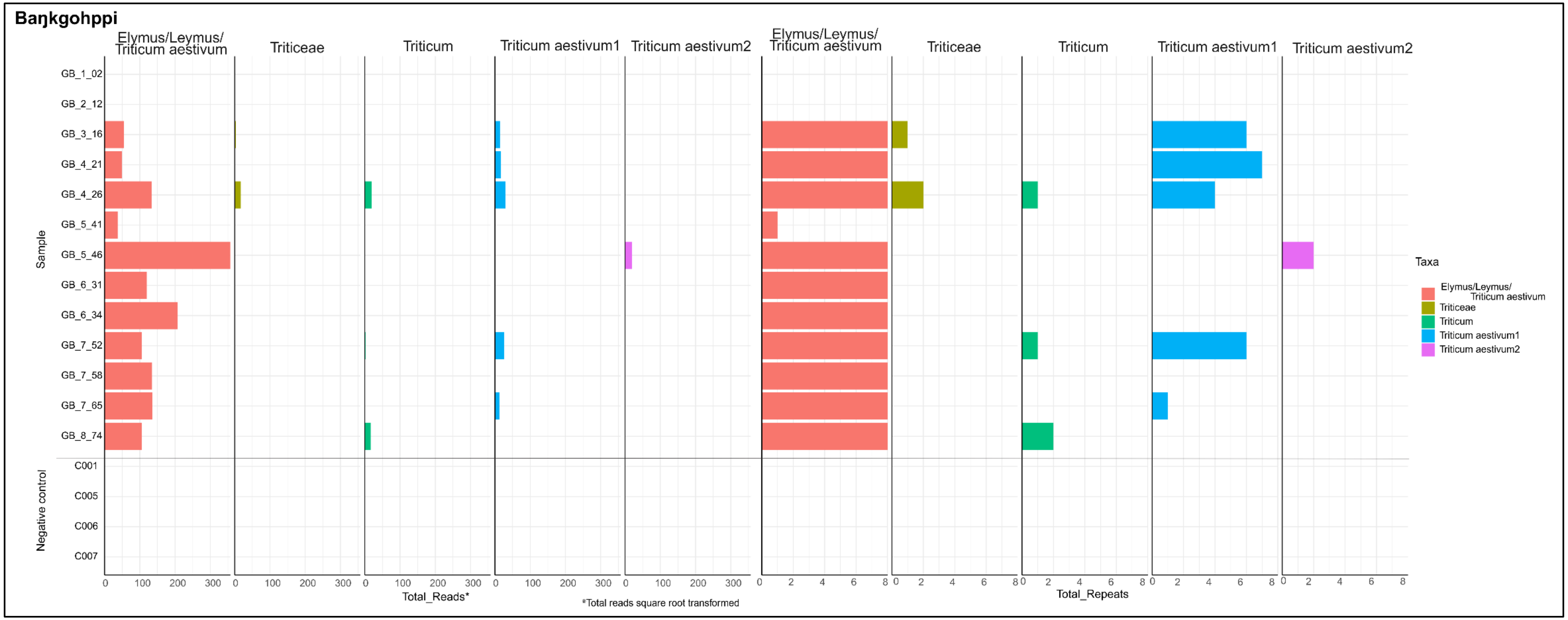

S26. Total reads and total repeats of *Triticum* at Finnes (P6-Loop).

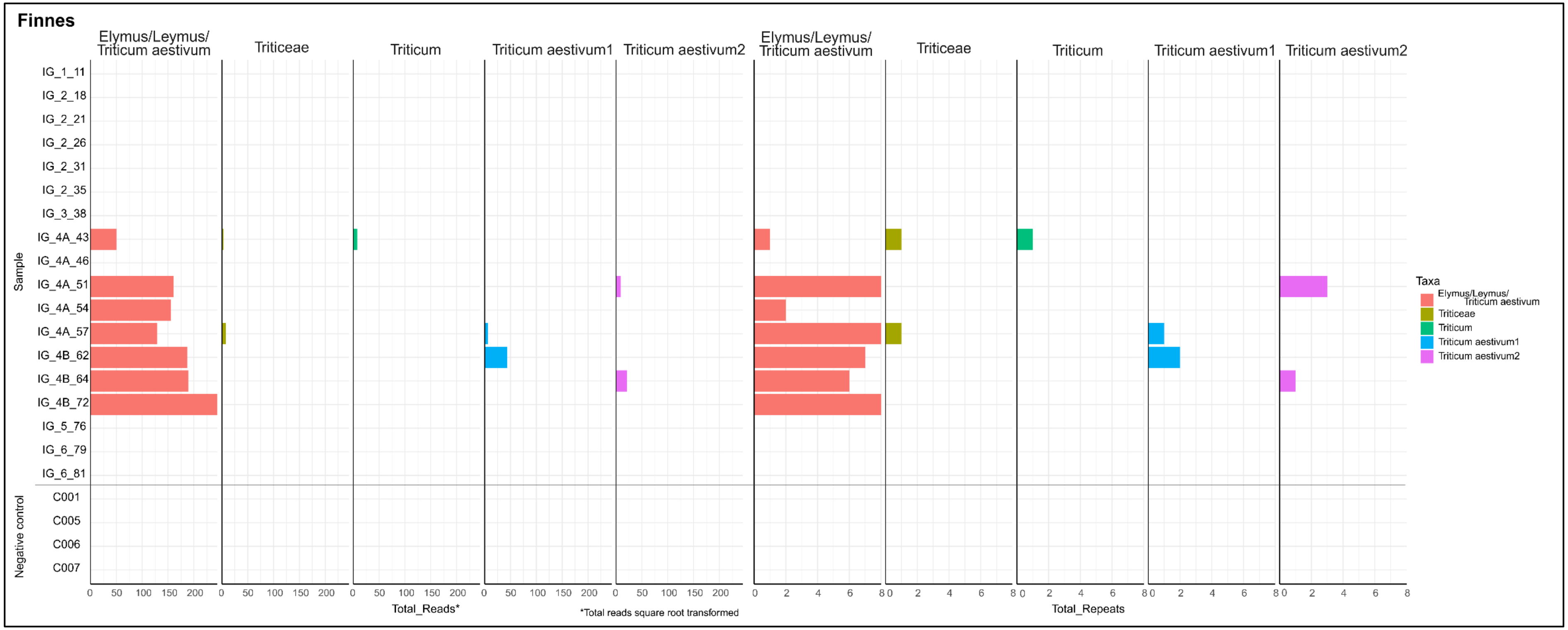

S27. Total reads and total repeats of *Pinus sylvestris* at Baŋkgohppi (P6-Loop).

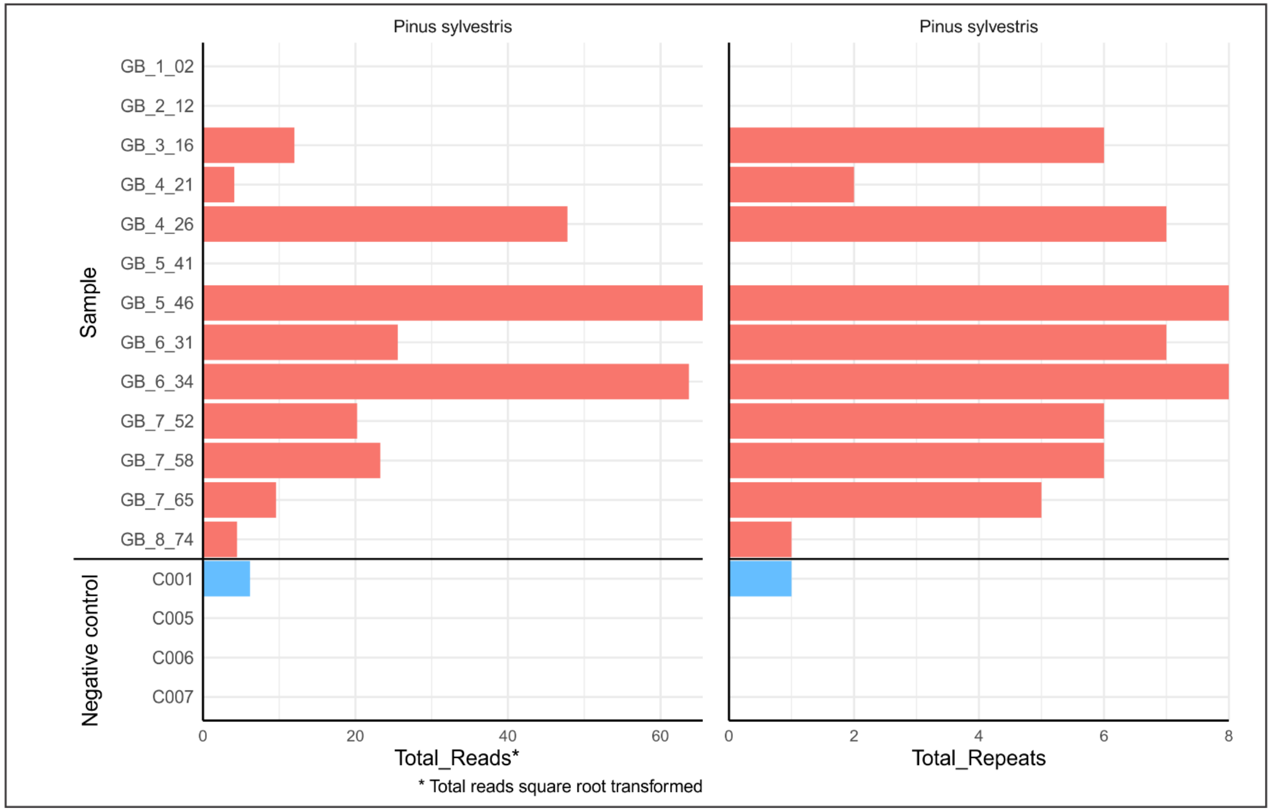

S28. Baŋkgohppi: food list for the nutritional analysis.

**Baŋkgohppi**

| Food_List | Quantity (1 unit = 100g) | |
|---|---|---|
| | GB_Raw | GB_Cooked |
| Egg, duck, whole, fresh, raw | 0.10 | 0.10 |
| Duck, wild, native, meat and skin, raw | 0.80 | 0.00 |
| Duck, wild, native, cooked | 0.00 | 0.80 |
| Animal fat, duck | 0.10 | 0.10 |
| Celery, raw | 1.00 | 0.00 |
| Celery, boiled, drained | 0.00 | 1.00 |
| Fiddlehead greens (ferns), raw | 1.00 | 0.00 |
| Fiddlehead greens (ferns), frozen, boiled, drained | 0.00 | 1.00 |
| Game meat, whale, raw | 1.00 | 1.00 |
| Game meat, native, beaver, raw | 0.50 | 0.00 |
| Game meat, native, beaver, roasted | 0.00 | 0.50 |
| Game meat, native, beaver, meat, dried | 0.30 | 0.30 |
| Game meat, native, beaver, feet, roasted | 0.05 | 0.05 |
| Game meat, native, beaver, tail, roasted | 0.05 | 0.05 |
| Game meat, native, beaver, liver, raw | 0.10 | 0.10 |
| Fireweed, native, leaves, raw | 0.50 | 0.50 |
| Fireweed, native, young leaves, raw | 0.50 | 0.50 |
| Black crowberry (curlewberry), native | 1.00 | 1.00 |
| Game meat, native, bearded seal meat, raw | 0.50 | 0.00 |
| Game meat, native, bearded seal meat, boiled | 0.00 | 0.50 |
| Game meat, native, bearded seal, intestine, raw | 0.10 | 0.00 |
| Game meat, native, bearded seal, intestine, boiled | 0.00 | 0.10 |
| Game meat, native, bearded seal, blubber, raw | 0.10 | 0.10 |
| Game meat, native, bearded seal, (oogruk), air−dried | 0.20 | 0.20 |
| Animal fat, native, bearded seal, oil (oogruk oil) | 0.10 | 0.10 |
| Seaweed, kelp (kombu, tangle), raw | 1.00 | 1.00 |
| Fish, cod (scrod), atlantic, raw | 0.40 | 0.00 |
| Fish, cod (scrod), atlantic, baked or broiled | 0.00 | 0.50 |
| Fish, cod (scrod), atlantic, smoked | 0.00 | 0.10 |
| Fish, cod (scrod), Atlantic, dried and salted, soaked in water | 0.20 | 0.10 |
| Fish, cod (scrod), atlantic, dried and salted | 0.10 | 0.10 |
| Fish, cod (scrod), atlantic, tongues and cheeks, raw | 0.20 | 0.10 |
| Fish oil, cod liver | 0.10 | 0.10 |
| Purslane, raw | 1.00 | 0.00 |
| Purslane, boiled, drained | 0.00 | 1.00 |
| Ptarmigan, native, meat, raw | 1.00 | 0.00 |
| Ptarmigan, native, meat, cooked | 0.00 | 1.00 |
| Fish, haddock, raw | 1.00 | 0.00 |
| Fish, haddock (finnan haddie), smoked | 0.00 | 0.50 |
| Fish, haddock, baked or broiled | 0.00 | 0.50 |
| Fish, hake (whiting), mixed species, raw | 1.00 | 0.00 |
| Fish, hake (whiting), mixed species, baked or broiled | 0.00 | 1.00 |
| Fish, ling, raw | 1.00 | 0.00 |
| Fish, ling, baked or broiled | 0.00 | 1.00 |
| Mollusks, mussel, blue, raw | 1.00 | 0.00 |
| Mollusks, mussel, blue, boiled or steamed | 0.00 | 1.00 |
| Seaweed, dulse (laver, nori), raw | 1.00 | 1.00 |
| Game meat, native, ringed seal, meat, raw | 0.50 | 0.00 |
| Game meat, native, ringed seal, meat, boiled | 0.00 | 0.50 |
| Game meat, native, ringed seal, meat, aged | 0.41 | 0.41 |
| Game meat, native, ringed seal, blubber, raw | 0.01 | 0.00 |
| Game meat, native, ringed seal, blubber, boiled | 0.00 | 0.01 |
| Game meat, native, ringed seal, blubber, aged, liquid | 0.01 | 0.01 |
| Game meat, native, ringed seal, blubber, aged, solid | 0.01 | 0.01 |
| Game meat, native, ringed seal, heart, raw | 0.01 | 0.01 |
| Game meat, native, ringed seal, brain, raw | 0.01 | 0.01 |
| Game meat, native, ringed seal, eyes, raw | 0.01 | 0.01 |
| Game meat, native, ringed seal, liver, raw | 0.01 | 0.01 |
| Game meat, native, ringed seal, pup, raw | 0.01 | 0.00 |
| Game meat, native, ringed seal, pup, boiled | 0.00 | 0.01 |
| Game meat, native, ringed seal, flippers, mattak, aged | 0.01 | 0.01 |
| Nuts, pine nuts, pignolia, dried | 1.00 | 1.00 |
| Fish, pollock, (alaska), raw | 1.00 | 0.00 |
| Fish, pollock (alaska), baked or broiled | 0.00 | 1.00 |
| Fish, pollock, atlantic (boston blue), raw | 1.00 | 0.00 |
| Fish, pollock, atlantic (boston blue), baked or broiled | 0.00 | 1.00 |
| Game meat, native, caribou (reindeer), meat, raw | 0.50 | 0.00 |
| Game meat, native, caribou (reindeer), meat, cooked | 0.00 | 0.50 |
| Game meat, native, caribou (reindeer), meat, dried | 0.21 | 0.21 |
| Game meat, native, caribou (reindeer), meat, aged | 0.20 | 0.20 |
| Game meat, native, caribou (reindeer), bone marrow, raw | 0.01 | 0.00 |
| Game meat, native, caribou (reindeer), bone marrow, cooked | 0.00 | 0.01 |
| Game meat, native, caribou (reindeer), fat, raw | 0.01 | 0.01 |
| Game meat, native, caribou (reindeer), brain, raw | 0.01 | 0.00 |
| Game meat, native, caribou (reindeer), brain, boiled | 0.00 | 0.01 |
| Game meat, native, caribou (reindeer), heart, raw | 0.01 | 0.00 |
| Game meat, native, caribou (reindeer), heart, boiled | 0.00 | 0.01 |
| Game meat, native, caribou (reindeer), lungs, raw | 0.01 | 0.01 |
| Game meat, native, caribou (reindeer), liver, raw | 0.01 | 0.00 |
| Game meat, native, caribou (reindeer), liver, baked | 0.00 | 0.01 |
| Game meat, native, caribou (reindeer), stomach, raw | 0.01 | 0.01 |
| Game meat, native, caribou (reindeer), stomach, contents | 0.01 | 0.01 |
| Game meat, native, caribou (reindeer), tongue, raw | 0.01 | 0.00 |
| Game meat, native, caribou (reindeer), tongue, cooked | 0.00 | 0.01 |
| Cloudberry (bakeapple), native | 1.00 | 1.00 |
| Dock (sorrel), raw | 1.00 | 0.00 |
| Dock (sorrel), boiled, drained | 0.00 | 1.00 |
| Willow, native, young leaves, chopped | 0.50 | 0.50 |
| Netted willow, native, leaves, raw | 0.50 | 0.50 |
| Stinging Nettles, native, blanched | 1.00 | 1.00 |
| Blueberry, wild, raw | 1.00 | 1.00 |
| Cranberry (lingonberry), wild, raw | 1.00 | 1.00 |

S29. Finnes: food list for the nutritional analysis.

### Finnes

| Food_List | Quantity (1 unit = 100g) | |
|---|---|---|
| | IG_Raw | IG_Cooked |
| Celery, raw | 1.000 | 0.000 |
| Celery, boiled, drained | 0.000 | 1.000 |
| Egg, goose, whole, fresh, raw | 0.100 | 0.100 |
| Animal fat, goose | 0.100 | 0.100 |
| Goose, wild (Canada goose), native, meat only, raw | 0.200 | 0.000 |
| Goose, wild (Canada goose), native, meat only, roasted | 0.000 | 0.100 |
| Goose, wild (Canada goose), native, meat only, boiled | 0.000 | 0.100 |
| Goose, wild (Canada goose), native, breast, meat and skin, raw | 0.200 | 0.000 |
| Goose, wild (Canada goose), native, breast, meat and skin, fire roasted | 0.000 | 0.100 |
| Goose, wild (Canada goose), native, breast, meat and skin, boiled | 0.000 | 0.100 |
| Goose, wild (Canada goose), native, breast, skinless, raw | 0.150 | 0.050 |
| Goose, wild (Canada goose), native, leg, meat and skin, raw | 0.050 | 0.000 |
| Goose, wild (Canada goose), native, leg, meat and skin, fire roasted | 0.000 | 0.050 |
| Goose, wild (Canada goose), native, leg, meat and skin, boiled | 0.000 | 0.050 |
| Goose, wild (Canada goose), native, collar, meat and skin, roasted | 0.050 | 0.050 |
| Goose, wild (Canada goose), native, collar, meat and skin, fire roasted | 0.050 | 0.050 |
| Goose, wild (Canada goose), native, liver, raw | 0.050 | 0.000 |
| Goose, wild (Canada goose), native, liver, roasted | 0.000 | 0.050 |
| Goose, wild (Canada goose), native, lungs, roasted | 0.050 | 0.050 |
| Goose, wild (Canada goose), native, meat and skin, dried, boiled | 0.000 | 0.050 |
| Fiddlehead greens (ferns), raw | 1.000 | 0.000 |
| Fiddlehead greens (ferns), frozen, boiled, drained | 0.000 | 1.000 |
| Fish, cusk, raw | 1.000 | 0.000 |
| Fish, cusk, baked or broiled | 0.000 | 1.000 |
| Mollusks, whelk, unspecified, raw | 1.000 | 0.000 |
| Mollusks, whelk, unspecified, boiled or steamed | 0.000 | 1.000 |
| Game meat, goat, raw | 0.500 | 0.000 |
| Game meat, goat, roasted | 0.000 | 0.500 |
| Milk, fluid, goat, unenriched, whole | 0.300 | 0.300 |
| Yogourt, goat, plain | 0.100 | 0.100 |
| Cheese, goat, soft, (more than 55% water, 21% M.F.) | 0.100 | 0.100 |
| Spices, caraway seed | 0.067 | 0.067 |
| Fireweed, native, leaves, raw | 0.500 | 0.500 |
| Fireweed, native, young leaves, raw | 0.500 | 0.500 |
| Mollusks, scallop, mixed species, raw | 1.000 | 0.000 |
| Mollusks, scallop (bay and sea), cooked, steamed | 0.000 | 1.000 |
| Black crowberry (curlewberry), native | 1.000 | 1.000 |
| Fish oil, cod liver | 0.100 | 0.100 |
| Fish, cod (scrod), atlantic, raw | 0.500 | 0.000 |
| Fish, cod (scrod), atlantic, baked or broiled | 0.000 | 0.250 |
| Fish, cod (scrod), atlantic, smoked | 0.000 | 0.250 |
| Fish, cod (scrod), atlantic, tongues and cheeks, raw | 0.200 | 0.200 |
| Fish, cod (scrod), Atlantic, dried and salted, soaked in water | 0.100 | 0.100 |
| Fish, cod (scrod), atlantic, dried and salted | 0.100 | 0.100 |
| Mollusks, clam, mixed species, raw | 1.000 | 0.000 |
| Mollusks, clam, mixed species, boiled or steamed | 0.000 | 1.000 |
| Fish, halibut, Atlantic or Pacific, raw | 1.000 | 0.000 |
| Fish, halibut, Atlantic or Pacific, baked or broiled | 0.000 | 1.000 |
| Purslane, raw | 1.000 | 0.000 |
| Purslane, boiled, drained | 0.000 | 1.000 |
| Fish, ling, raw | 1.000 | 0.000 |
| Fish, ling, baked or broiled | 0.000 | 1.000 |
| Mollusks, mussel, blue, raw | 1.000 | 0.000 |
| Mollusks, mussel, blue, boiled or steamed | 0.000 | 1.000 |
| Mollusks, oyster, eastern (blue point), wild, raw | 1.000 | 0.000 |
| Mollusks, oyster, eastern (blue point), wild, boiled or steamed | 0.000 | 0.500 |
| Mollusks, oyster, eastern (blue point), wild, baked or broiled | 0.000 | 0.500 |
| Seaweed, dulse (laver, nori), raw | 1.000 | 1.000 |
| Seaweed, dulse (laver, nori), dried | 1.000 | 1.000 |
| Nuts, pine nuts, pignolia, dried | 1.000 | 1.000 |
| Fish, pollock, atlantic (boston blue), raw | 1.000 | 0.000 |
| Fish, pollock, atlantic (boston blue), baked or broiled | 0.000 | 1.000 |
| Game meat, native, caribou (reindeer), meat, raw | 0.500 | 0.000 |
| Game meat, native, caribou (reindeer), meat, cooked | 0.000 | 0.500 |
| Game meat, native, caribou (reindeer), meat, dried | 0.210 | 0.210 |
| Game meat, native, caribou (reindeer), meat, aged | 0.200 | 0.200 |
| Game meat, native, caribou (reindeer), bone marrow, raw | 0.010 | 0.000 |
| Game meat, native, caribou (reindeer), bone marrow, cooked | 0.000 | 0.010 |
| Game meat, native, caribou (reindeer), fat, raw | 0.010 | 0.010 |
| Game meat, native, caribou (reindeer), brain, raw | 0.010 | 0.000 |
| Game meat, native, caribou (reindeer), brain, boiled | 0.000 | 0.010 |
| Game meat, native, caribou (reindeer), heart, raw | 0.010 | 0.000 |
| Game meat, native, caribou (reindeer), heart, boiled | 0.000 | 0.010 |
| Game meat, native, caribou (reindeer), lungs, raw | 0.010 | 0.010 |
| Game meat, native, caribou (reindeer), stomach, raw | 0.010 | 0.010 |
| Game meat, native, caribou (reindeer), stomach, contents | 0.010 | 0.010 |
| Game meat, native, caribou (reindeer), liver, raw | 0.010 | 0.000 |
| Game meat, native, caribou (reindeer), liver, baked | 0.000 | 0.010 |
| Game meat, native, caribou (reindeer), tongue, raw | 0.010 | 0.000 |
| Game meat, native, caribou (reindeer), tongue, cooked | 0.000 | 0.010 |
| Cloudberry (bakeapple), native | 1.000 | 1.000 |
| Dock (sorrel), raw | 1.000 | 0.000 |
| Game meat, native, caribou (reindeer), liver, raw | 0.010 | 0.000 |
| Willow, native, young leaves, chopped | 0.000 | 0.010 |
| Game meat, native, caribou (reindeer), tongue, raw | 0.010 | 0.000 |
| Stinging Nettles, native, blanched | 0.000 | 0.010 |
| Cloudberry (bakeapple), native | 1.000 | 1.000 |

S30. Percentages of RDA for each nutrient for Baŋkgohppi.

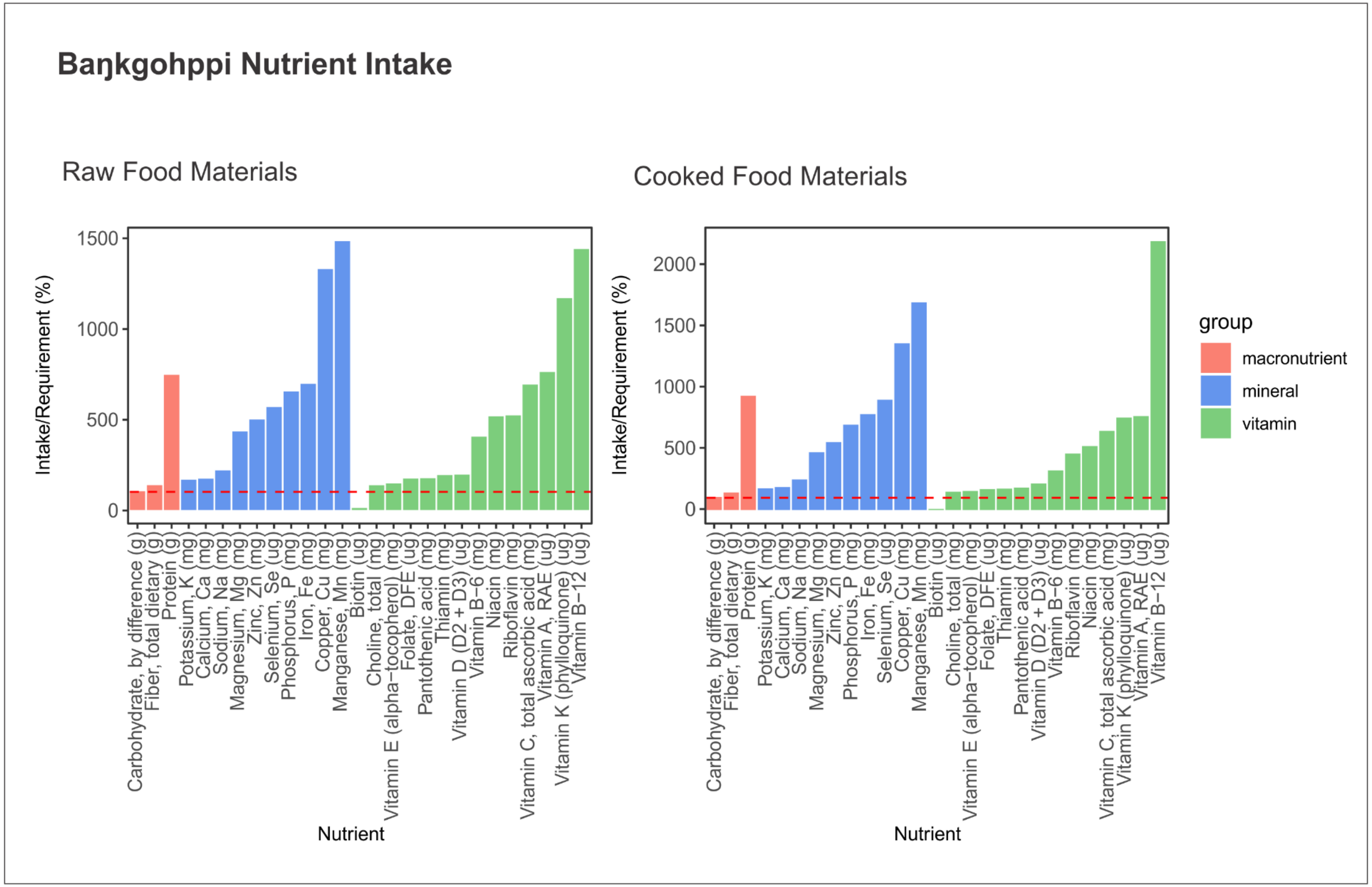

S31. Percentages of RDA for each nutrient for Finnes.

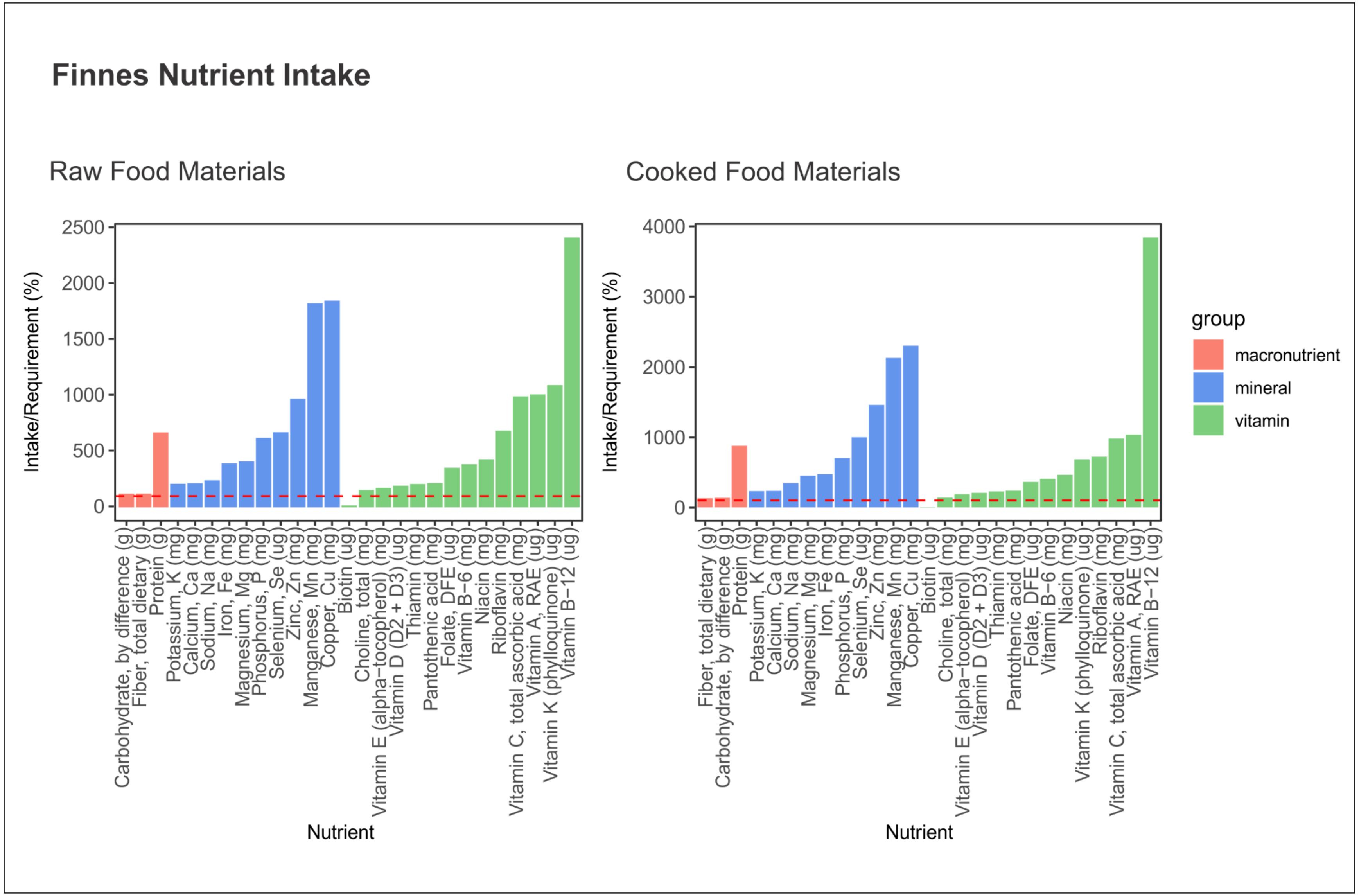

S32. Contribution of food items for each nutrient for Baŋkgohppi (raw materials).

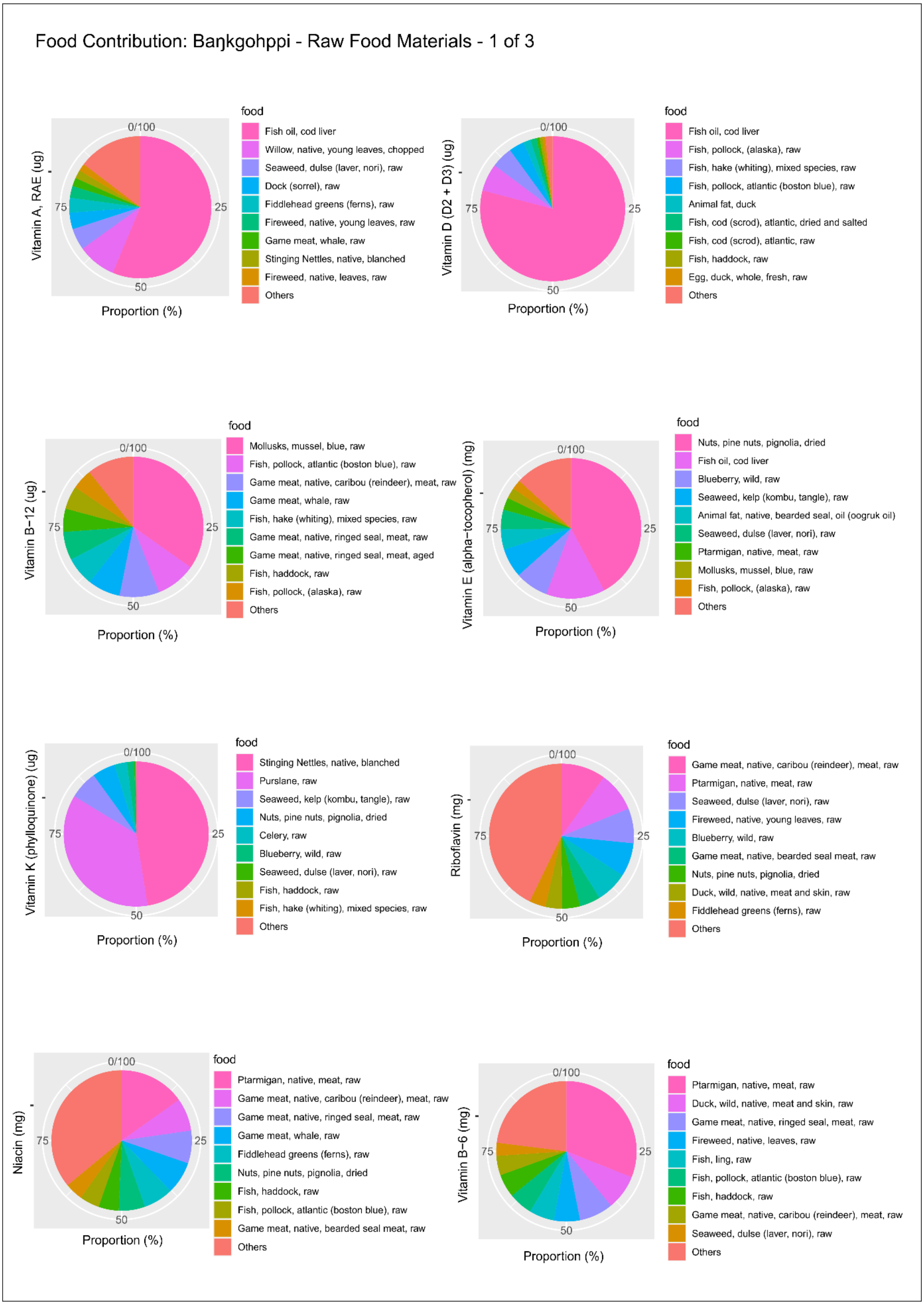

S32. Cont.

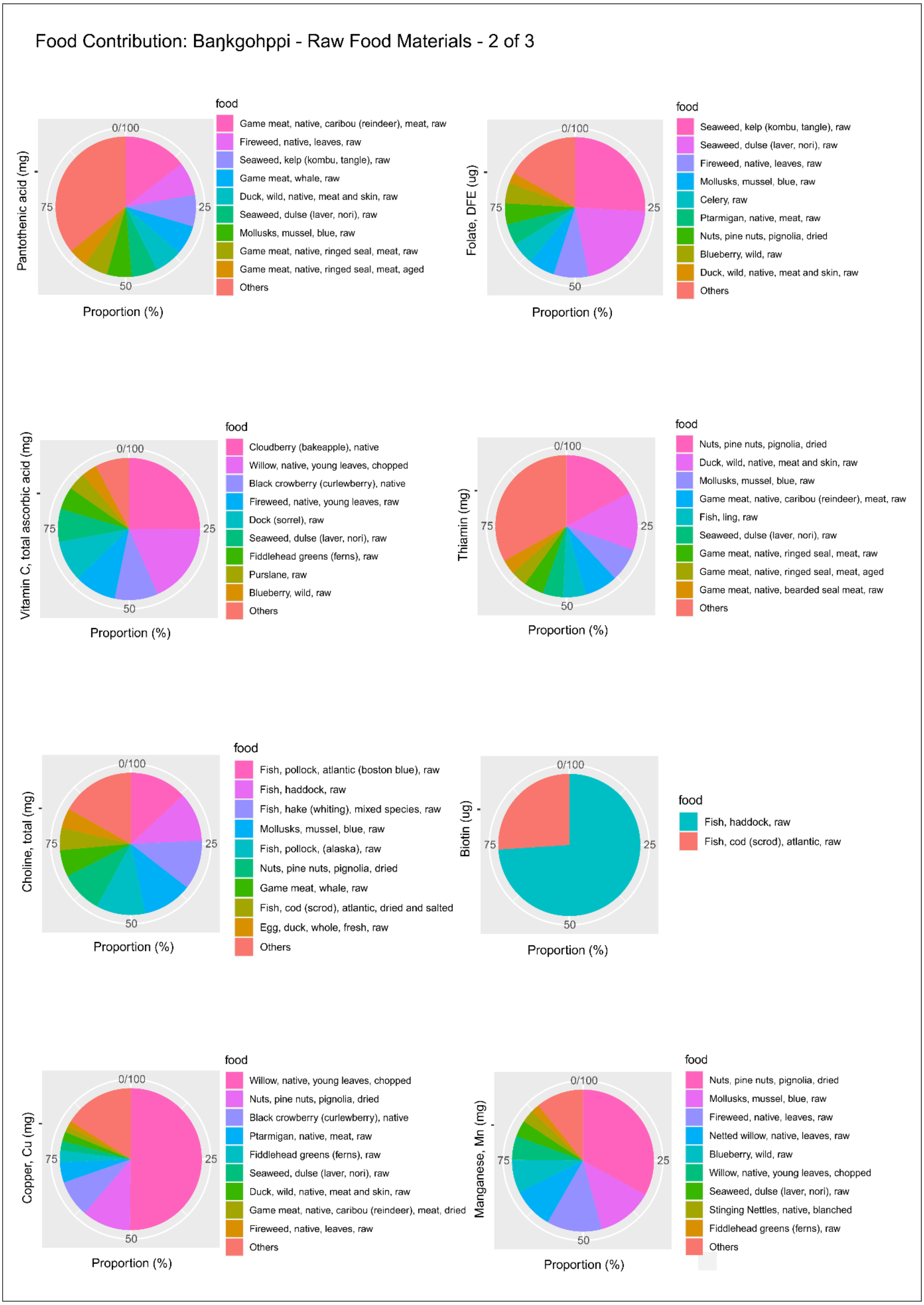
Food Contribution: Baŋkgohppi - Raw Food Materials - 2 of 3
Pantothenic acid (mg)
Proportion (%)
food
Game meat, native, caribou (reindeer), meat, raw
Fireweed, native, leaves, raw
Seaweed, kelp (kombu, tangle), raw
Game meat, whale, raw
Duck, wild, native, meat and skin, raw
Seaweed, dulse (laver, nori), raw
Mollusks, mussel, blue, raw
Game meat, native, ringed seal, meat, raw
Game meat, native, ringed seal, meat, aged
Others
Folate, DFE (ug)
Proportion (%)
food
Seaweed, kelp (kombu, tangle), raw
Seaweed, dulse (laver, nori), raw
Fireweed, native, leaves, raw
Mollusks, mussel, blue, raw
Celery, raw
Ptarmigan, native, meat, raw
Nuts, pine nuts, pignolia, dried
Blueberry, wild, raw
Duck, wild, native, meat and skin, raw
Others
Vitamin C, total ascorbic acid (mg)
Proportion (%)
food
Cloudberry (bakeapple), native
Willow, native, young leaves, chopped
Black crowberry (curlewberry), native
Fireweed, native, young leaves, raw
Dock (sorrel), raw
Seaweed, dulse (laver, nori), raw
Fiddlehead greens (ferns), raw
Purslane, raw
Blueberry, wild, raw
Others
Thiamin (mg)
Proportion (%)
food
Nuts, pine nuts, pignolia, dried
Duck, wild, native, meat and skin, raw
Mollusks, mussel, blue, raw
Game meat, native, caribou (reindeer), meat, raw
Fish, ling, raw
Seaweed, dulse (laver, nori), raw
Game meat, native, ringed seal, meat, raw
Game meat, native, ringed seal, meat, aged
Game meat, native, bearded seal meat, raw
Others
Choline, total (mg)
Proportion (%)
food
Fish, pollock, atlantic (boston blue), raw
Fish, haddock, raw
Fish, hake (whiting), mixed species, raw
Mollusks, mussel, blue, raw
Fish, pollock, (alaska), raw
Nuts, pine nuts, pignolia, dried
Game meat, whale, raw
Fish, cod (scrod), atlantic, dried and salted
Egg, duck, whole, fresh, raw
Others
Biotin (ug)
Proportion (%)
food
Fish, haddock, raw
Fish, cod (scrod), atlantic, raw
Copper, Cu (mg)
Proportion (%)
food
Willow, native, young leaves, chopped
Nuts, pine nuts, pignolia, dried
Black crowberry (curlewberry), native
Ptarmigan, native, meat, raw
Fiddlehead greens (ferns), raw
Seaweed, dulse (laver, nori), raw
Duck, wild, native, meat and skin, raw
Game meat, native, caribou (reindeer), meat, dried
Fireweed, native, leaves, raw
Others
Manganese, Mn (mg)
Proportion (%)
food
Nuts, pine nuts, pignolia, dried
Mollusks, mussel, blue, raw
Fireweed, native, leaves, raw
Netted willow, native, leaves, raw
Blueberry, wild, raw
Willow, native, young leaves, chopped
Seaweed, dulse (laver, nori), raw
Stinging Nettles, native, blanched
Fiddlehead greens (ferns), raw
Others

S32. Cont.

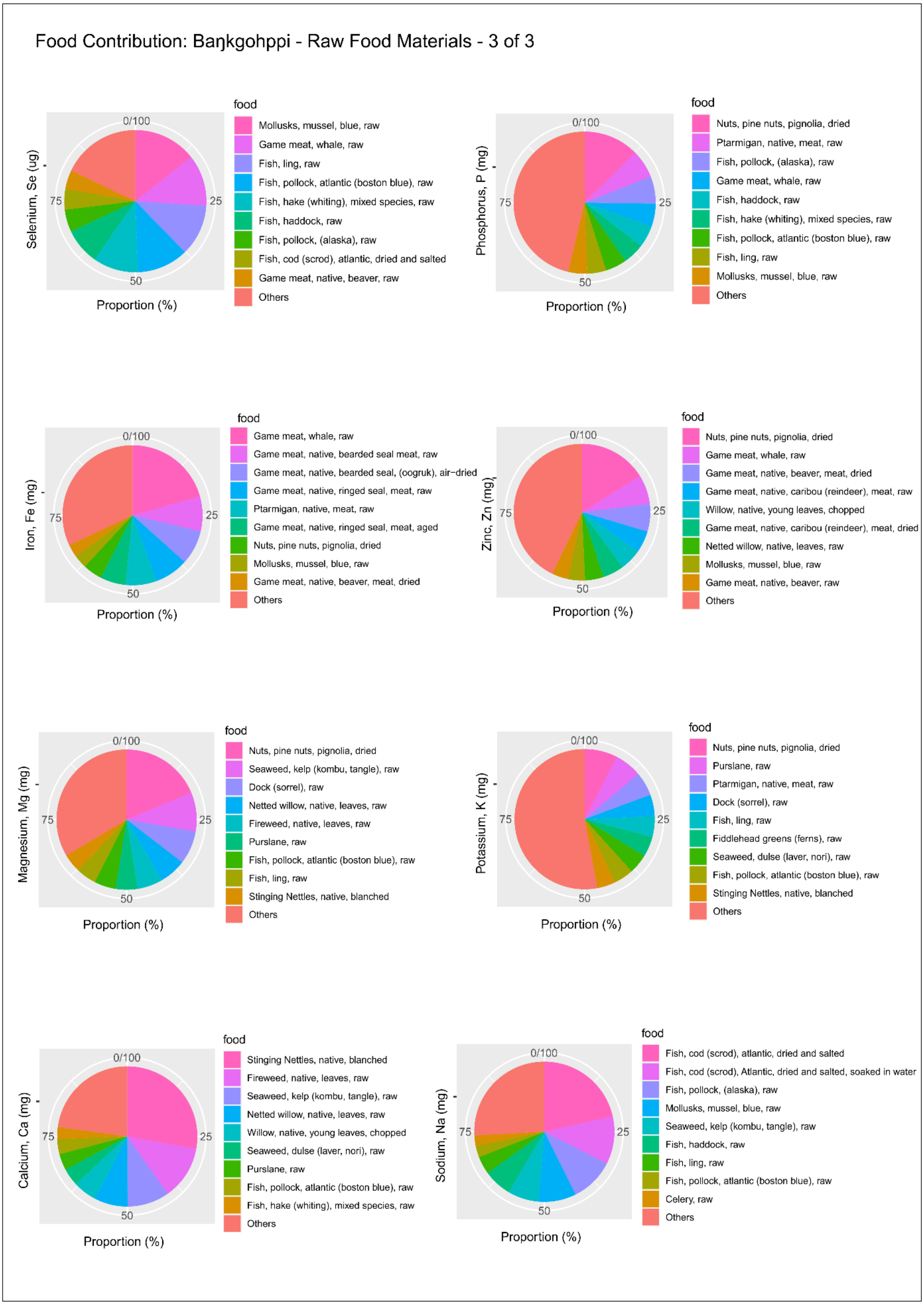
Food Contribution: Baŋkgohppi - Raw Food Materials - 3 of 3
Selenium, Se (ug)
0/100
25
50
75
Proportion (%)
food
Mollusks, mussel, blue, raw
Game meat, whale, raw
Fish, ling, raw
Fish, pollock, atlantic (boston blue), raw
Fish, hake (whiting), mixed species, raw
Fish, haddock, raw
Fish, pollock, (alaska), raw
Fish, cod (scrod), atlantic, dried and salted
Game meat, native, beaver, raw
Others
Phosphorus, P (mg)
0/100
25
50
75
Proportion (%)
food
Nuts, pine nuts, pignolia, dried
Ptarmigan, native, meat, raw
Fish, pollock, (alaska), raw
Game meat, whale, raw
Fish, haddock, raw
Fish, hake (whiting), mixed species, raw
Fish, pollock, atlantic (boston blue), raw
Fish, ling, raw
Mollusks, mussel, blue, raw
Others
Iron, Fe (mg)
0/100
25
50
75
Proportion (%)
food
Game meat, whale, raw
Game meat, native, bearded seal meat, raw
Game meat, native, bearded seal, (oogruk), air−dried
Game meat, native, ringed seal, meat, raw
Ptarmigan, native, meat, raw
Game meat, native, ringed seal, meat, aged
Nuts, pine nuts, pignolia, dried
Mollusks, mussel, blue, raw
Game meat, native, beaver, meat, dried
Others
Zinc, Zn (mg)
0/100
25
50
75
Proportion (%)
food
Nuts, pine nuts, pignolia, dried
Game meat, whale, raw
Game meat, native, beaver, meat, dried
Game meat, native, caribou (reindeer), meat, raw
Willow, native, young leaves, chopped
Game meat, native, caribou (reindeer), meat, dried
Netted willow, native, leaves, raw
Mollusks, mussel, blue, raw
Game meat, native, beaver, raw
Others
Magnesium, Mg (mg)
0/100
25
50
75
Proportion (%)
food
Nuts, pine nuts, pignolia, dried
Seaweed, kelp (kombu, tangle), raw
Dock (sorrel), raw
Netted willow, native, leaves, raw
Fireweed, native, leaves, raw
Purslane, raw
Fish, pollock, atlantic (boston blue), raw
Fish, ling, raw
Stinging Nettles, native, blanched
Others
Potassium, K (mg)
0/100
25
50
75
Proportion (%)
food
Nuts, pine nuts, pignolia, dried
Purslane, raw
Ptarmigan, native, meat, raw
Dock (sorrel), raw
Fish, ling, raw
Fiddlehead greens (ferns), raw
Seaweed, dulse (laver, nori), raw
Fish, pollock, atlantic (boston blue), raw
Stinging Nettles, native, blanched
Others
Calcium, Ca (mg)
0/100
25
50
75
Proportion (%)
food
Stinging Nettles, native, blanched
Fireweed, native, leaves, raw
Seaweed, kelp (kombu, tangle), raw
Netted willow, native, leaves, raw
Willow, native, young leaves, chopped
Seaweed, dulse (laver, nori), raw
Purslane, raw
Fish, pollock, atlantic (boston blue), raw
Fish, hake (whiting), mixed species, raw
Others
Sodium, Na (mg)
0/100
25
50
75
Proportion (%)
food
Fish, cod (scrod), atlantic, dried and salted
Fish, cod (scrod), Atlantic, dried and salted, soaked in water
Fish, pollock, (alaska), raw
Mollusks, mussel, blue, raw
Seaweed, kelp (kombu, tangle), raw
Fish, haddock, raw
Fish, ling, raw
Fish, pollock, atlantic (boston blue), raw
Celery, raw
Others

S33. Contribution of food items for each nutrient for Baŋkgohppi (cooked materials).

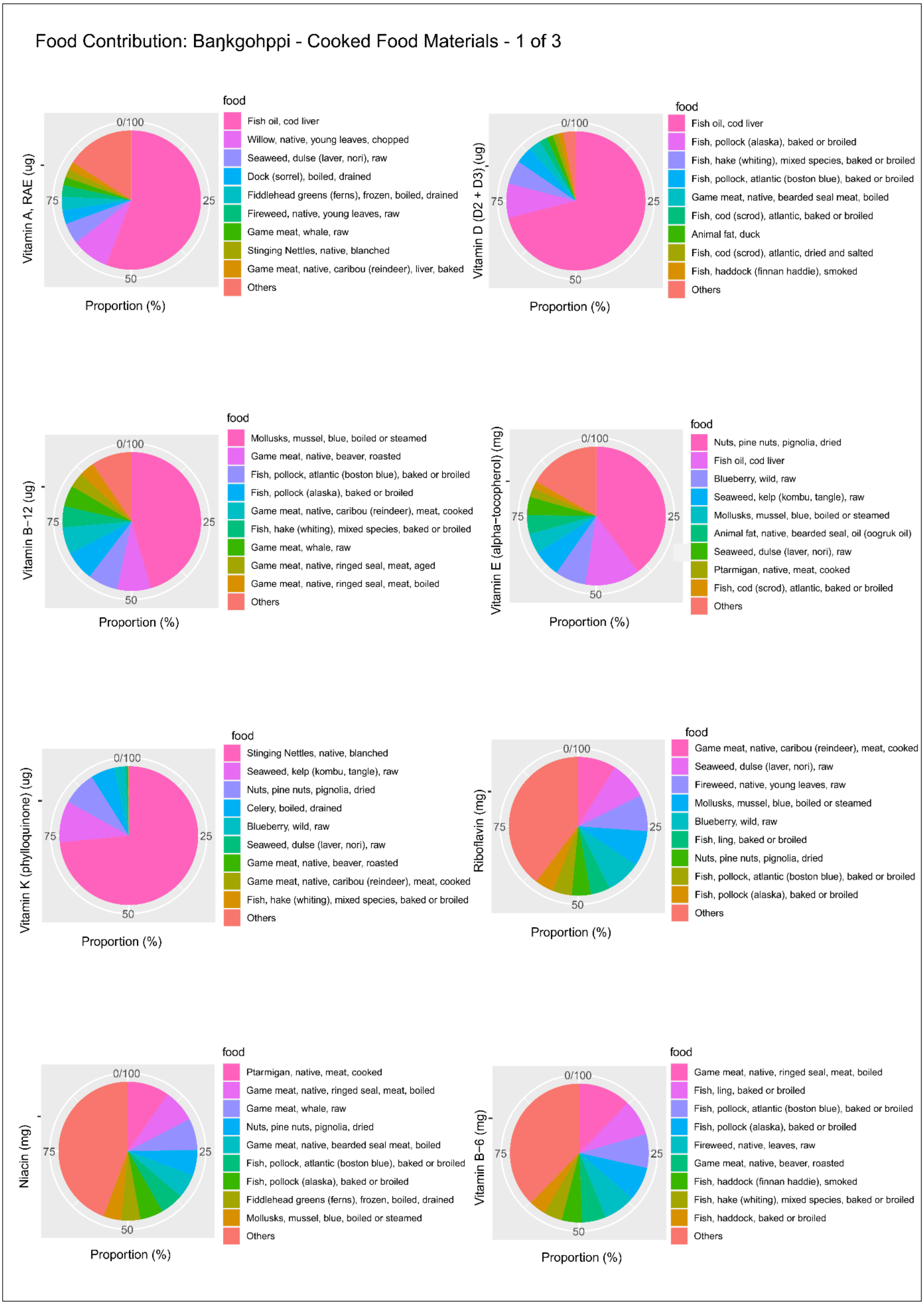

S33. Cont.

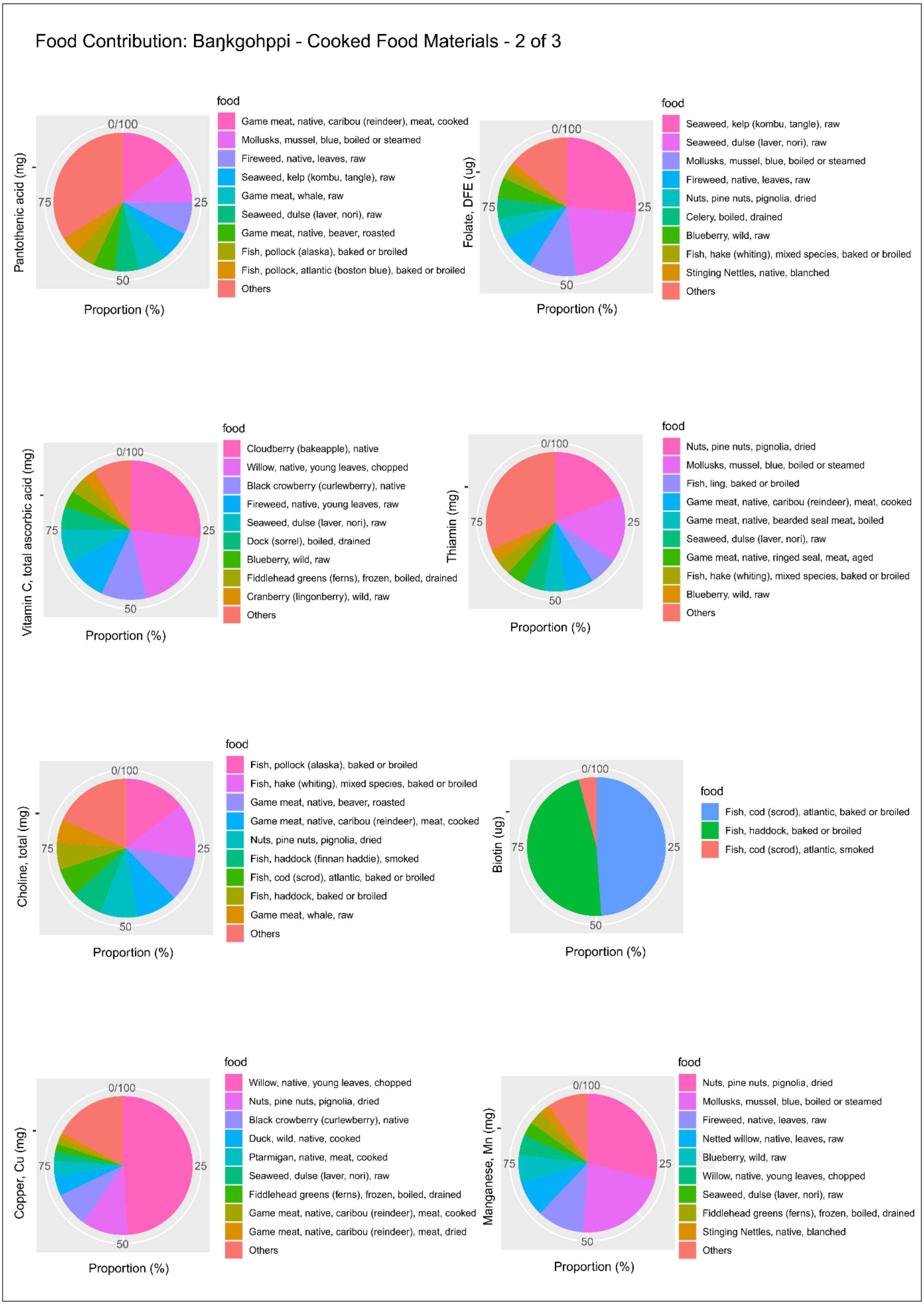
Food Contribution: Baŋkgohppi - Cooked Food Materials - 2 of 3
Pantothenic acid (mg)
0/100
25
50
75
Proportion (%)
food
Game meat, native, caribou (reindeer), meat, cooked
Mollusks, mussel, blue, boiled or steamed
Fireweed, native, leaves, raw
Seaweed, kelp (kombu, tangle), raw
Game meat, whale, raw
Seaweed, dulse (laver, nori), raw
Game meat, native, beaver, roasted
Fish, pollock (alaska), baked or broiled
Fish, pollock, atlantic (boston blue), baked or broiled
Others
Folate, DFE (ug)
0/100
25
50
75
Proportion (%)
food
Seaweed, kelp (kombu, tangle), raw
Seaweed, dulse (laver, nori), raw
Mollusks, mussel, blue, boiled or steamed
Fireweed, native, leaves, raw
Nuts, pine nuts, pignolia, dried
Celery, boiled, drained
Blueberry, wild, raw
Fish, hake (whiting), mixed species, baked or broiled
Stinging Nettles, native, blanched
Others
Vitamin C, total ascorbic acid (mg)
0/100
25
50
75
Proportion (%)
food
Cloudberry (bakeapple), native
Willow, native, young leaves, chopped
Black crowberry (curlewberry), native
Fireweed, native, young leaves, raw
Seaweed, dulse (laver, nori), raw
Dock (sorrel), boiled, drained
Blueberry, wild, raw
Fiddlehead greens (ferns), frozen, boiled, drained
Cranberry (lingonberry), wild, raw
Others
Thiamin (mg)
0/100
25
50
75
Proportion (%)
food
Nuts, pine nuts, pignolia, dried
Mollusks, mussel, blue, boiled or steamed
Fish, ling, baked or broiled
Game meat, native, caribou (reindeer), meat, cooked
Game meat, native, bearded seal meat, boiled
Seaweed, dulse (laver, nori), raw
Game meat, native, ringed seal, meat, aged
Fish, hake (whiting), mixed species, baked or broiled
Blueberry, wild, raw
Others
Choline, total (mg)
0/100
25
50
75
Proportion (%)
food
Fish, pollock (alaska), baked or broiled
Fish, hake (whiting), mixed species, baked or broiled
Game meat, native, beaver, roasted
Game meat, native, caribou (reindeer), meat, cooked
Nuts, pine nuts, pignolia, dried
Fish, haddock (finnan haddie), smoked
Fish, cod (scrod), atlantic, baked or broiled
Fish, haddock, baked or broiled
Game meat, whale, raw
Others
Biotin (ug)
0/100
25
50
75
Proportion (%)
food
Fish, cod (scrod), atlantic, baked or broiled
Fish, haddock, baked or broiled
Fish, cod (scrod), atlantic, smoked
Copper, Cu (mg)
0/100
25
50
75
Proportion (%)
food
Willow, native, young leaves, chopped
Nuts, pine nuts, pignolia, dried
Black crowberry (curlewberry), native
Duck, wild, native, cooked
Ptarmigan, native, meat, cooked
Seaweed, dulse (laver, nori), raw
Fiddlehead greens (ferns), frozen, boiled, drained
Game meat, native, caribou (reindeer), meat, cooked
Game meat, native, caribou (reindeer), meat, dried
Others
Manganese, Mn (mg)
0/100
25
50
75
Proportion (%)
food
Nuts, pine nuts, pignolia, dried
Mollusks, mussel, blue, boiled or steamed
Fireweed, native, leaves, raw
Netted willow, native, leaves, raw
Blueberry, wild, raw
Willow, native, young leaves, chopped
Seaweed, dulse (laver, nori), raw
Fiddlehead greens (ferns), frozen, boiled, drained
Stinging Nettles, native, blanched
Others

S33. Cont.

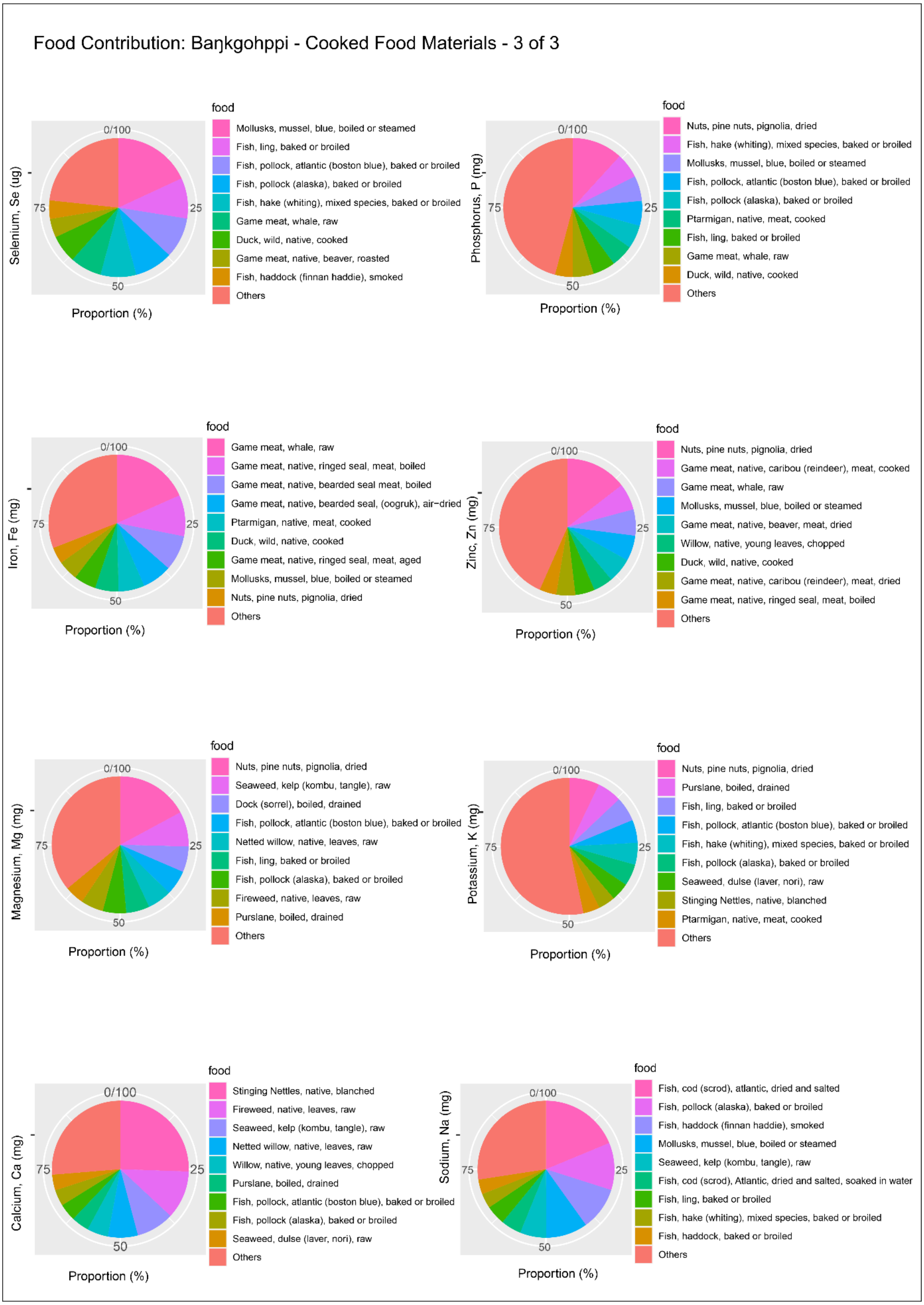

Food Contribution: Baŋkgohppi - Cooked Food Materials - 3 of 3
Selenium, Se (ug)
0/100
25
50
75
Proportion (%)
food
Mollusks, mussel, blue, boiled or steamed
Fish, ling, baked or broiled
Fish, pollock, atlantic (boston blue), baked or broiled
Fish, pollock (alaska), baked or broiled
Fish, hake (whiting), mixed species, baked or broiled
Game meat, whale, raw
Duck, wild, native, cooked
Game meat, native, beaver, roasted
Fish, haddock (finnan haddie), smoked
Others
Phosphorus, P (mg)
0/100
25
50
75
Proportion (%)
food
Nuts, pine nuts, pignolia, dried
Fish, hake (whiting), mixed species, baked or broiled
Mollusks, mussel, blue, boiled or steamed
Fish, pollock, atlantic (boston blue), baked or broiled
Fish, pollock (alaska), baked or broiled
Ptarmigan, native, meat, cooked
Fish, ling, baked or broiled
Game meat, whale, raw
Duck, wild, native, cooked
Others
Iron, Fe (mg)
0/100
25
50
75
Proportion (%)
food
Game meat, whale, raw
Game meat, native, ringed seal, meat, boiled
Game meat, native, bearded seal meat, boiled
Game meat, native, bearded seal, (oogruk), air−dried
Ptarmigan, native, meat, cooked
Duck, wild, native, cooked
Game meat, native, ringed seal, meat, aged
Mollusks, mussel, blue, boiled or steamed
Nuts, pine nuts, pignolia, dried
Others
Zinc, Zn (mg)
0/100
25
50
75
Proportion (%)
food
Nuts, pine nuts, pignolia, dried
Game meat, native, caribou (reindeer), meat, cooked
Game meat, whale, raw
Mollusks, mussel, blue, boiled or steamed
Game meat, native, beaver, meat, dried
Willow, native, young leaves, chopped
Duck, wild, native, cooked
Game meat, native, caribou (reindeer), meat, dried
Game meat, native, ringed seal, meat, boiled
Others
Magnesium, Mg (mg)
0/100
25
50
75
Proportion (%)
food
Nuts, pine nuts, pignolia, dried
Seaweed, kelp (kombu, tangle), raw
Dock (sorrel), boiled, drained
Fish, pollock, atlantic (boston blue), baked or broiled
Netted willow, native, leaves, raw
Fish, ling, baked or broiled
Fish, pollock (alaska), baked or broiled
Fireweed, native, leaves, raw
Purslane, boiled, drained
Others
Potassium, K (mg)
0/100
25
50
75
Proportion (%)
food
Nuts, pine nuts, pignolia, dried
Purslane, boiled, drained
Fish, ling, baked or broiled
Fish, pollock, atlantic (boston blue), baked or broiled
Fish, hake (whiting), mixed species, baked or broiled
Fish, pollock (alaska), baked or broiled
Seaweed, dulse (laver, nori), raw
Stinging Nettles, native, blanched
Ptarmigan, native, meat, cooked
Others
Calcium, Ca (mg)
0/100
25
50
75
Proportion (%)
food
Stinging Nettles, native, blanched
Fireweed, native, leaves, raw
Seaweed, kelp (kombu, tangle), raw
Netted willow, native, leaves, raw
Willow, native, young leaves, chopped
Purslane, boiled, drained
Fish, pollock, atlantic (boston blue), baked or broiled
Fish, pollock (alaska), baked or broiled
Seaweed, dulse (laver, nori), raw
Others
Sodium, Na (mg)
0/100
25
50
75
Proportion (%)
food
Fish, cod (scrod), atlantic, dried and salted
Fish, pollock (alaska), baked or broiled
Fish, haddock (finnan haddie), smoked
Mollusks, mussel, blue, boiled or steamed
Seaweed, kelp (kombu, tangle), raw
Fish, cod (scrod), Atlantic, dried and salted, soaked in water
Fish, ling, baked or broiled
Fish, hake (whiting), mixed species, baked or broiled
Fish, haddock, baked or broiled
Others

S34. Contribution of food items for each nutrient for Finnes (raw materials).

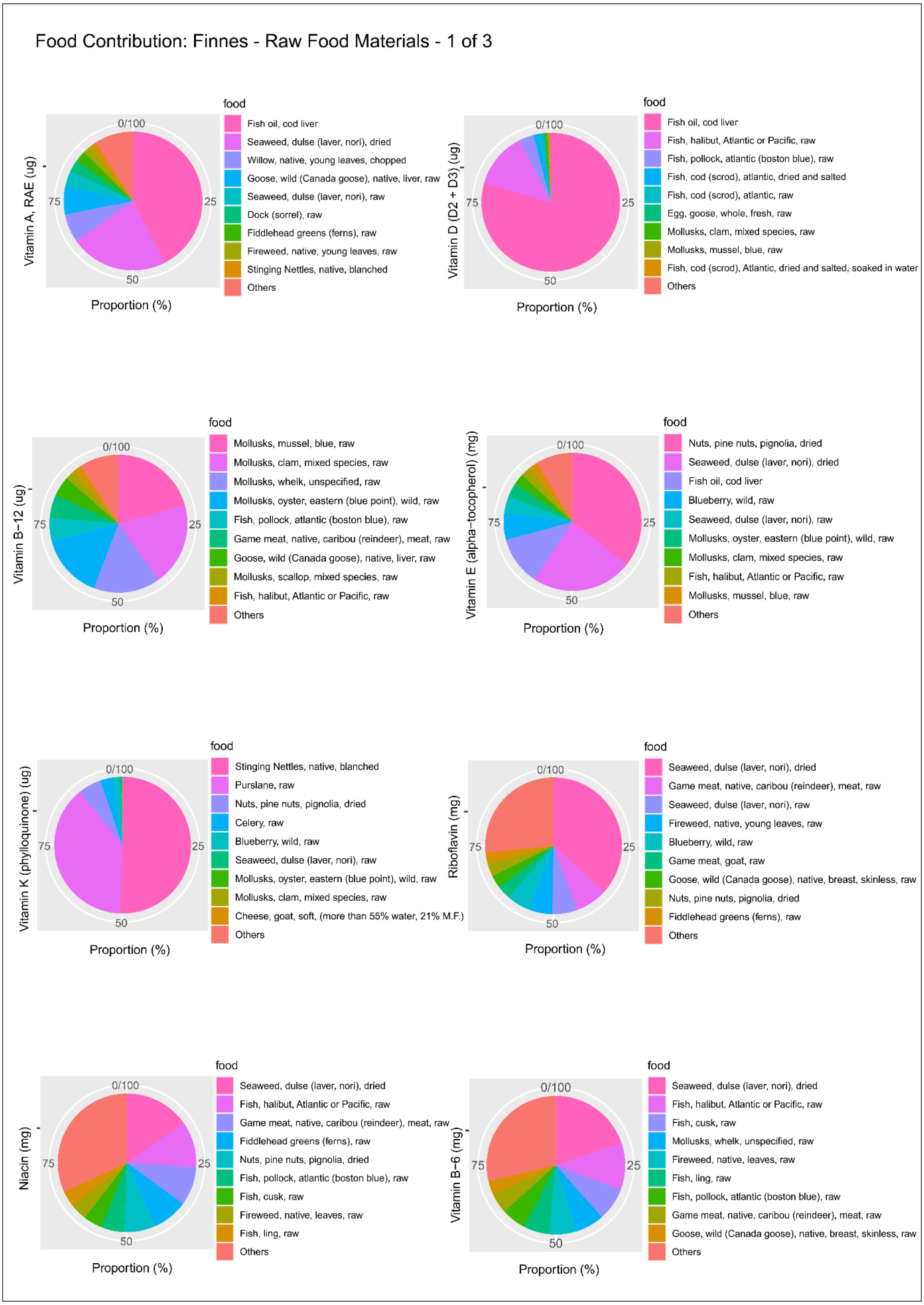

S34. Cont.

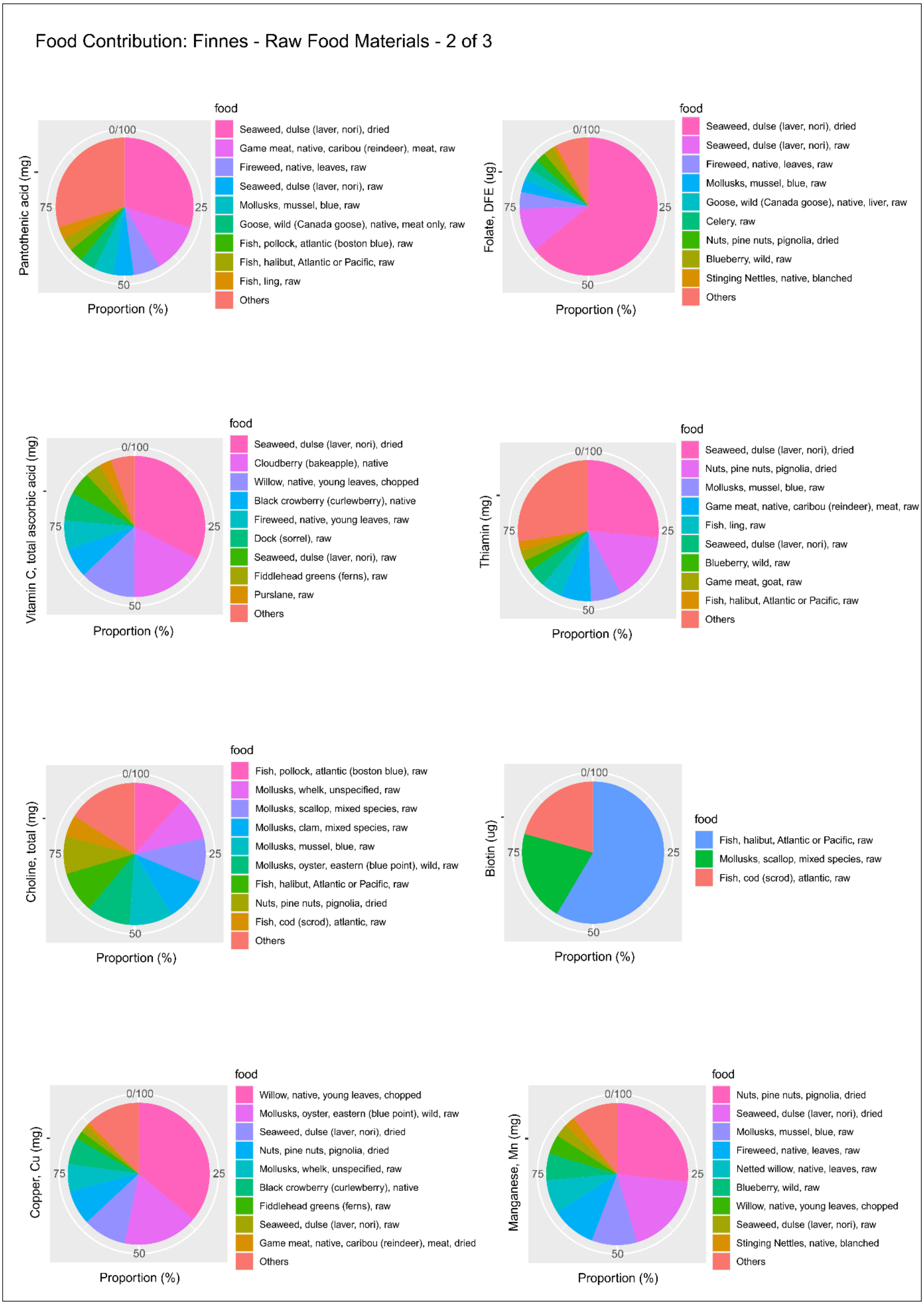
Food Contribution: Finnes - Raw Food Materials - 2 of 3
Pantothenic acid (mg)
0/100
25
50
75
Proportion (%)
food
Seaweed, dulse (laver, nori), dried
Game meat, native, caribou (reindeer), meat, raw
Fireweed, native, leaves, raw
Seaweed, dulse (laver, nori), raw
Mollusks, mussel, blue, raw
Goose, wild (Canada goose), native, meat only, raw
Fish, pollock, atlantic (boston blue), raw
Fish, halibut, Atlantic or Pacific, raw
Fish, ling, raw
Others
Folate, DFE (ug)
0/100
25
50
75
Proportion (%)
food
Seaweed, dulse (laver, nori), dried
Seaweed, dulse (laver, nori), raw
Fireweed, native, leaves, raw
Mollusks, mussel, blue, raw
Goose, wild (Canada goose), native, liver, raw
Celery, raw
Nuts, pine nuts, pignolia, dried
Blueberry, wild, raw
Stinging Nettles, native, blanched
Others
Vitamin C, total ascorbic acid (mg)
0/100
25
50
75
Proportion (%)
food
Seaweed, dulse (laver, nori), dried
Cloudberry (bakeapple), native
Willow, native, young leaves, chopped
Black crowberry (curlewberry), native
Fireweed, native, young leaves, raw
Dock (sorrel), raw
Seaweed, dulse (laver, nori), raw
Fiddlehead greens (ferns), raw
Purslane, raw
Others
Thiamin (mg)
0/100
25
50
75
Proportion (%)
food
Seaweed, dulse (laver, nori), dried
Nuts, pine nuts, pignolia, dried
Mollusks, mussel, blue, raw
Game meat, native, caribou (reindeer), meat, raw
Fish, ling, raw
Seaweed, dulse (laver, nori), raw
Blueberry, wild, raw
Game meat, goat, raw
Fish, halibut, Atlantic or Pacific, raw
Others
Choline, total (mg)
0/100
25
50
75
Proportion (%)
food
Fish, pollock, atlantic (boston blue), raw
Mollusks, whelk, unspecified, raw
Mollusks, scallop, mixed species, raw
Mollusks, clam, mixed species, raw
Mollusks, mussel, blue, raw
Mollusks, oyster, eastern (blue point), wild, raw
Fish, halibut, Atlantic or Pacific, raw
Nuts, pine nuts, pignolia, dried
Fish, cod (scrod), atlantic, raw
Others
Biotin (ug)
0/100
25
50
75
Proportion (%)
food
Fish, halibut, Atlantic or Pacific, raw
Mollusks, scallop, mixed species, raw
Fish, cod (scrod), atlantic, raw
Copper, Cu (mg)
0/100
25
50
75
Proportion (%)
food
Willow, native, young leaves, chopped
Mollusks, oyster, eastern (blue point), wild, raw
Seaweed, dulse (laver, nori), dried
Nuts, pine nuts, pignolia, dried
Mollusks, whelk, unspecified, raw
Black crowberry (curlewberry), native
Fiddlehead greens (ferns), raw
Seaweed, dulse (laver, nori), raw
Game meat, native, caribou (reindeer), meat, dried
Others
Manganese, Mn (mg)
0/100
25
50
75
Proportion (%)
food
Nuts, pine nuts, pignolia, dried
Seaweed, dulse (laver, nori), dried
Mollusks, mussel, blue, raw
Fireweed, native, leaves, raw
Netted willow, native, leaves, raw
Blueberry, wild, raw
Willow, native, young leaves, chopped
Seaweed, dulse (laver, nori), raw
Stinging Nettles, native, blanched
Others

S34. Cont.

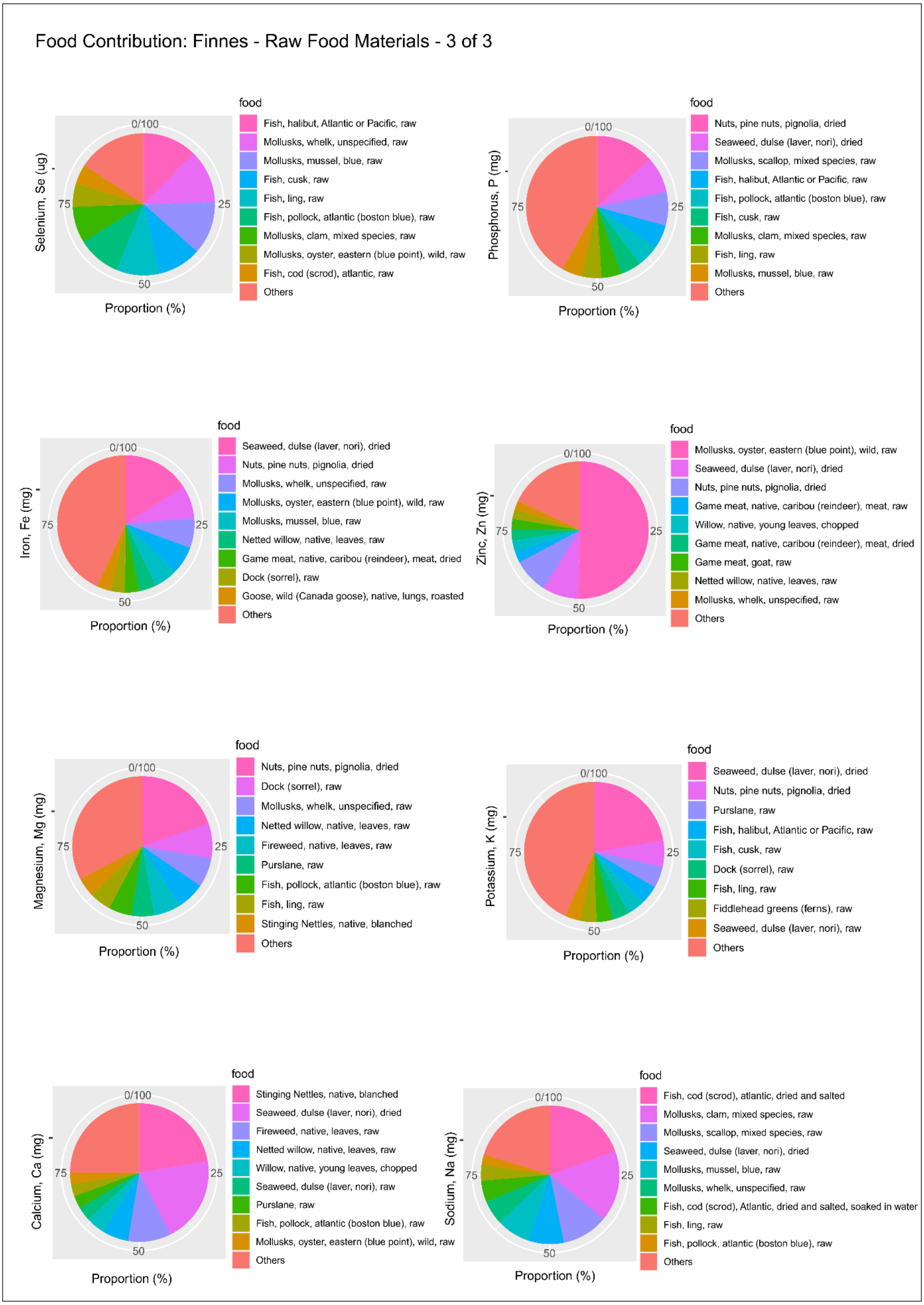
Food Contribution: Finnes - Raw Food Materials - 3 of 3
Selenium, Se (ug)
0/100
25
50
75
Proportion (%)
food
Fish, halibut, Atlantic or Pacific, raw
Mollusks, whelk, unspecified, raw
Mollusks, mussel, blue, raw
Fish, cusk, raw
Fish, ling, raw
Fish, pollock, atlantic (boston blue), raw
Mollusks, clam, mixed species, raw
Mollusks, oyster, eastern (blue point), wild, raw
Fish, cod (scrod), atlantic, raw
Others
Phosphorus, P (mg)
0/100
25
50
75
Proportion (%)
food
Nuts, pine nuts, pignolia, dried
Seaweed, dulse (laver, nori), dried
Mollusks, scallop, mixed species, raw
Fish, halibut, Atlantic or Pacific, raw
Fish, pollock, atlantic (boston blue), raw
Fish, cusk, raw
Mollusks, clam, mixed species, raw
Fish, ling, raw
Mollusks, mussel, blue, raw
Others
Iron, Fe (mg)
0/100
25
50
75
Proportion (%)
food
Seaweed, dulse (laver, nori), dried
Nuts, pine nuts, pignolia, dried
Mollusks, whelk, unspecified, raw
Mollusks, oyster, eastern (blue point), wild, raw
Mollusks, mussel, blue, raw
Netted willow, native, leaves, raw
Game meat, native, caribou (reindeer), meat, dried
Dock (sorrel), raw
Goose, wild (Canada goose), native, lungs, roasted
Others
Zinc, Zn (mg)
0/100
25
50
75
Proportion (%)
food
Mollusks, oyster, eastern (blue point), wild, raw
Seaweed, dulse (laver, nori), dried
Nuts, pine nuts, pignolia, dried
Game meat, native, caribou (reindeer), meat, raw
Willow, native, young leaves, chopped
Game meat, native, caribou (reindeer), meat, dried
Game meat, goat, raw
Netted willow, native, leaves, raw
Mollusks, whelk, unspecified, raw
Others
Magnesium, Mg (mg)
0/100
25
50
75
Proportion (%)
food
Nuts, pine nuts, pignolia, dried
Dock (sorrel), raw
Mollusks, whelk, unspecified, raw
Netted willow, native, leaves, raw
Fireweed, native, leaves, raw
Purslane, raw
Fish, pollock, atlantic (boston blue), raw
Fish, ling, raw
Stinging Nettles, native, blanched
Others
Potassium, K (mg)
0/100
25
50
75
Proportion (%)
food
Seaweed, dulse (laver, nori), dried
Nuts, pine nuts, pignolia, dried
Purslane, raw
Fish, halibut, Atlantic or Pacific, raw
Fish, cusk, raw
Dock (sorrel), raw
Fish, ling, raw
Fiddlehead greens (ferns), raw
Seaweed, dulse (laver, nori), raw
Others
Calcium, Ca (mg)
0/100
25
50
75
Proportion (%)
food
Stinging Nettles, native, blanched
Seaweed, dulse (laver, nori), dried
Fireweed, native, leaves, raw
Netted willow, native, leaves, raw
Willow, native, young leaves, chopped
Seaweed, dulse (laver, nori), raw
Purslane, raw
Fish, pollock, atlantic (boston blue), raw
Mollusks, oyster, eastern (blue point), wild, raw
Others
Sodium, Na (mg)
0/100
25
50
75
Proportion (%)
food
Fish, cod (scrod), atlantic, dried and salted
Mollusks, clam, mixed species, raw
Mollusks, scallop, mixed species, raw
Seaweed, dulse (laver, nori), dried
Mollusks, mussel, blue, raw
Mollusks, whelk, unspecified, raw
Fish, cod (scrod), Atlantic, dried and salted, soaked in water
Fish, ling, raw
Fish, pollock, atlantic (boston blue), raw
Others

S35. Contribution of food items for each nutrient for Finnes (cooked materials).

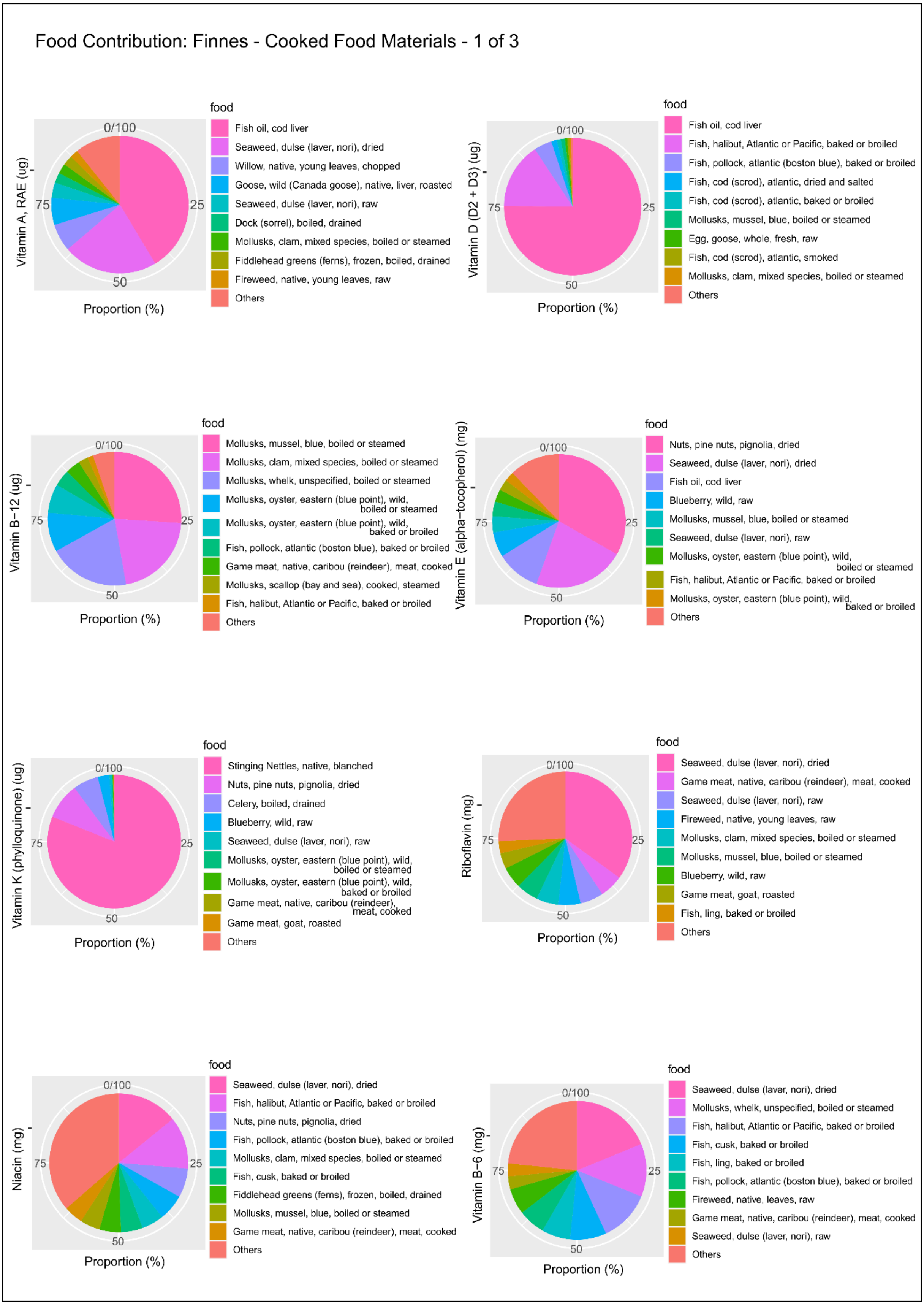

S35. Cont.

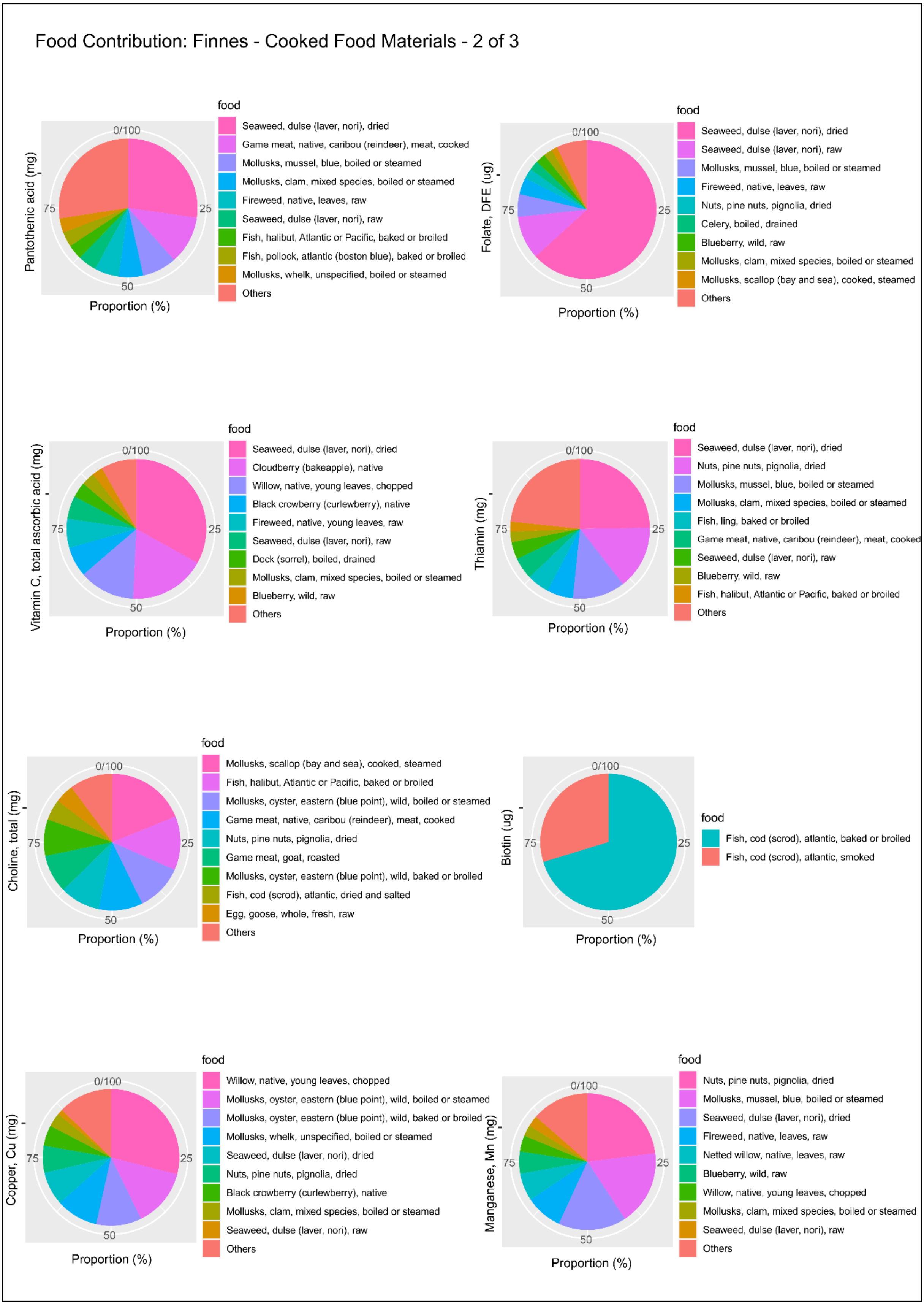
Food Contribution: Finnes - Cooked Food Materials - 2 of 3
Pantothenic acid (mg)
0/100
25
50
75
Proportion (%)
food
Seaweed, dulse (laver, nori), dried
Game meat, native, caribou (reindeer), meat, cooked
Mollusks, mussel, blue, boiled or steamed
Mollusks, clam, mixed species, boiled or steamed
Fireweed, native, leaves, raw
Seaweed, dulse (laver, nori), raw
Fish, halibut, Atlantic or Pacific, baked or broiled
Fish, pollock, atlantic (boston blue), baked or broiled
Mollusks, whelk, unspecified, boiled or steamed
Others
Folate, DFE (ug)
0/100
25
50
75
Proportion (%)
food
Seaweed, dulse (laver, nori), dried
Seaweed, dulse (laver, nori), raw
Mollusks, mussel, blue, boiled or steamed
Fireweed, native, leaves, raw
Nuts, pine nuts, pignolia, dried
Celery, boiled, drained
Blueberry, wild, raw
Mollusks, clam, mixed species, boiled or steamed
Mollusks, scallop (bay and sea), cooked, steamed
Others
Vitamin C, total ascorbic acid (mg)
0/100
25
50
75
Proportion (%)
food
Seaweed, dulse (laver, nori), dried
Cloudberry (bakeapple), native
Willow, native, young leaves, chopped
Black crowberry (curlewberry), native
Fireweed, native, young leaves, raw
Seaweed, dulse (laver, nori), raw
Dock (sorrel), boiled, drained
Mollusks, clam, mixed species, boiled or steamed
Blueberry, wild, raw
Others
Thiamin (mg)
0/100
25
50
75
Proportion (%)
food
Seaweed, dulse (laver, nori), dried
Nuts, pine nuts, pignolia, dried
Mollusks, mussel, blue, boiled or steamed
Mollusks, clam, mixed species, boiled or steamed
Fish, ling, baked or broiled
Game meat, native, caribou (reindeer), meat, cooked
Seaweed, dulse (laver, nori), raw
Blueberry, wild, raw
Fish, halibut, Atlantic or Pacific, baked or broiled
Others
Choline, total (mg)
0/100
25
50
75
Proportion (%)
food
Mollusks, scallop (bay and sea), cooked, steamed
Fish, halibut, Atlantic or Pacific, baked or broiled
Mollusks, oyster, eastern (blue point), wild, boiled or steamed
Game meat, native, caribou (reindeer), meat, cooked
Nuts, pine nuts, pignolia, dried
Game meat, goat, roasted
Mollusks, oyster, eastern (blue point), wild, baked or broiled
Fish, cod (scrod), atlantic, dried and salted
Egg, goose, whole, fresh, raw
Others
Biotin (ug)
0/100
25
50
75
Proportion (%)
food
Fish, cod (scrod), atlantic, baked or broiled
Fish, cod (scrod), atlantic, smoked
Copper, Cu (mg)
0/100
25
50
75
Proportion (%)
food
Willow, native, young leaves, chopped
Mollusks, oyster, eastern (blue point), wild, boiled or steamed
Mollusks, oyster, eastern (blue point), wild, baked or broiled
Mollusks, whelk, unspecified, boiled or steamed
Seaweed, dulse (laver, nori), dried
Nuts, pine nuts, pignolia, dried
Black crowberry (curlewberry), native
Mollusks, clam, mixed species, boiled or steamed
Seaweed, dulse (laver, nori), raw
Others
Manganese, Mn (mg)
0/100
25
50
75
Proportion (%)
food
Nuts, pine nuts, pignolia, dried
Mollusks, mussel, blue, boiled or steamed
Seaweed, dulse (laver, nori), dried
Fireweed, native, leaves, raw
Netted willow, native, leaves, raw
Blueberry, wild, raw
Willow, native, young leaves, chopped
Mollusks, clam, mixed species, boiled or steamed
Seaweed, dulse (laver, nori), raw
Others

S35. Cont.

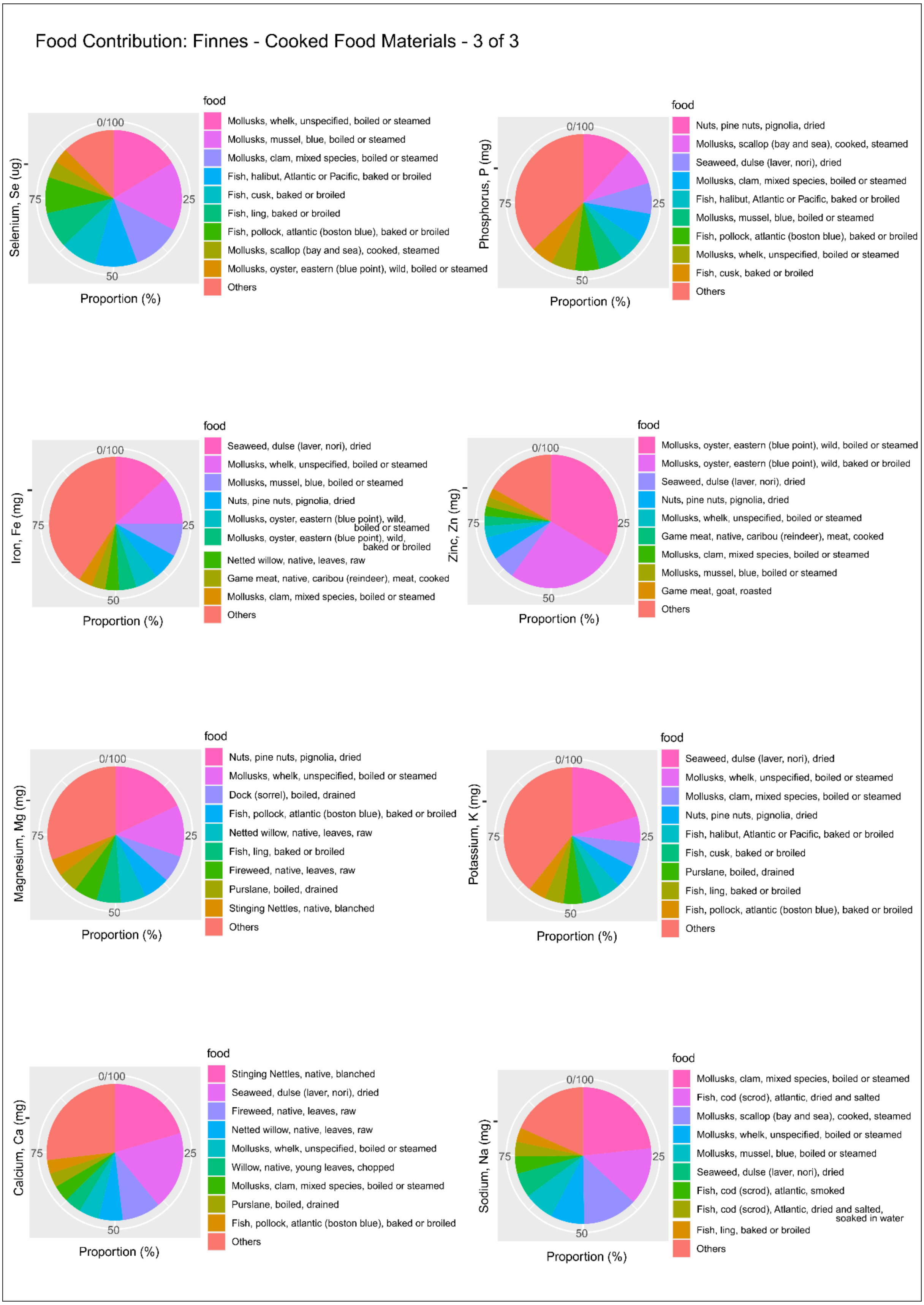

Food Contribution: Finnes - Cooked Food Materials - 3 of 3
Selenium, Se (ug)
0/100
25
50
75
Proportion (%)
food
Mollusks, whelk, unspecified, boiled or steamed
Mollusks, mussel, blue, boiled or steamed
Mollusks, clam, mixed species, boiled or steamed
Fish, halibut, Atlantic or Pacific, baked or broiled
Fish, cusk, baked or broiled
Fish, ling, baked or broiled
Fish, pollock, atlantic (boston blue), baked or broiled
Mollusks, scallop (bay and sea), cooked, steamed
Mollusks, oyster, eastern (blue point), wild, boiled or steamed
Others
Phosphorus, P (mg)
0/100
25
50
75
Proportion (%)
food
Nuts, pine nuts, pignolia, dried
Mollusks, scallop (bay and sea), cooked, steamed
Seaweed, dulse (laver, nori), dried
Mollusks, clam, mixed species, boiled or steamed
Fish, halibut, Atlantic or Pacific, baked or broiled
Mollusks, mussel, blue, boiled or steamed
Fish, pollock, atlantic (boston blue), baked or broiled
Mollusks, whelk, unspecified, boiled or steamed
Fish, cusk, baked or broiled
Others
Iron, Fe (mg)
0/100
25
50
75
Proportion (%)
food
Seaweed, dulse (laver, nori), dried
Mollusks, whelk, unspecified, boiled or steamed
Mollusks, mussel, blue, boiled or steamed
Nuts, pine nuts, pignolia, dried
Mollusks, oyster, eastern (blue point), wild, boiled or steamed
Mollusks, oyster, eastern (blue point), wild, baked or broiled
Netted willow, native, leaves, raw
Game meat, native, caribou (reindeer), meat, cooked
Mollusks, clam, mixed species, boiled or steamed
Others
Zinc, Zn (mg)
0/100
25
50
75
Proportion (%)
food
Mollusks, oyster, eastern (blue point), wild, boiled or steamed
Mollusks, oyster, eastern (blue point), wild, baked or broiled
Seaweed, dulse (laver, nori), dried
Nuts, pine nuts, pignolia, dried
Mollusks, whelk, unspecified, boiled or steamed
Game meat, native, caribou (reindeer), meat, cooked
Mollusks, clam, mixed species, boiled or steamed
Mollusks, mussel, blue, boiled or steamed
Game meat, goat, roasted
Others
Magnesium, Mg (mg)
0/100
25
50
75
Proportion (%)
food
Nuts, pine nuts, pignolia, dried
Mollusks, whelk, unspecified, boiled or steamed
Dock (sorrel), boiled, drained
Fish, pollock, atlantic (boston blue), baked or broiled
Netted willow, native, leaves, raw
Fish, ling, baked or broiled
Fireweed, native, leaves, raw
Purslane, boiled, drained
Stinging Nettles, native, blanched
Others
Potassium, K (mg)
0/100
25
50
75
Proportion (%)
food
Seaweed, dulse (laver, nori), dried
Mollusks, whelk, unspecified, boiled or steamed
Mollusks, clam, mixed species, boiled or steamed
Nuts, pine nuts, pignolia, dried
Fish, halibut, Atlantic or Pacific, baked or broiled
Fish, cusk, baked or broiled
Purslane, boiled, drained
Fish, ling, baked or broiled
Fish, pollock, atlantic (boston blue), baked or broiled
Others
Calcium, Ca (mg)
0/100
25
50
75
Proportion (%)
food
Stinging Nettles, native, blanched
Seaweed, dulse (laver, nori), dried
Fireweed, native, leaves, raw
Netted willow, native, leaves, raw
Mollusks, whelk, unspecified, boiled or steamed
Willow, native, young leaves, chopped
Mollusks, clam, mixed species, boiled or steamed
Purslane, boiled, drained
Fish, pollock, atlantic (boston blue), baked or broiled
Others
Sodium, Na (mg)
0/100
25
50
75
Proportion (%)
food
Mollusks, clam, mixed species, boiled or steamed
Fish, cod (scrod), atlantic, dried and salted
Mollusks, scallop (bay and sea), cooked, steamed
Mollusks, whelk, unspecified, boiled or steamed
Mollusks, mussel, blue, boiled or steamed
Seaweed, dulse (laver, nori), dried
Fish, cod (scrod), atlantic, smoked
Fish, cod (scrod), Atlantic, dried and salted, soaked in water
Fish, ling, baked or broiled
Others

S36. Harvesting season of some food items.

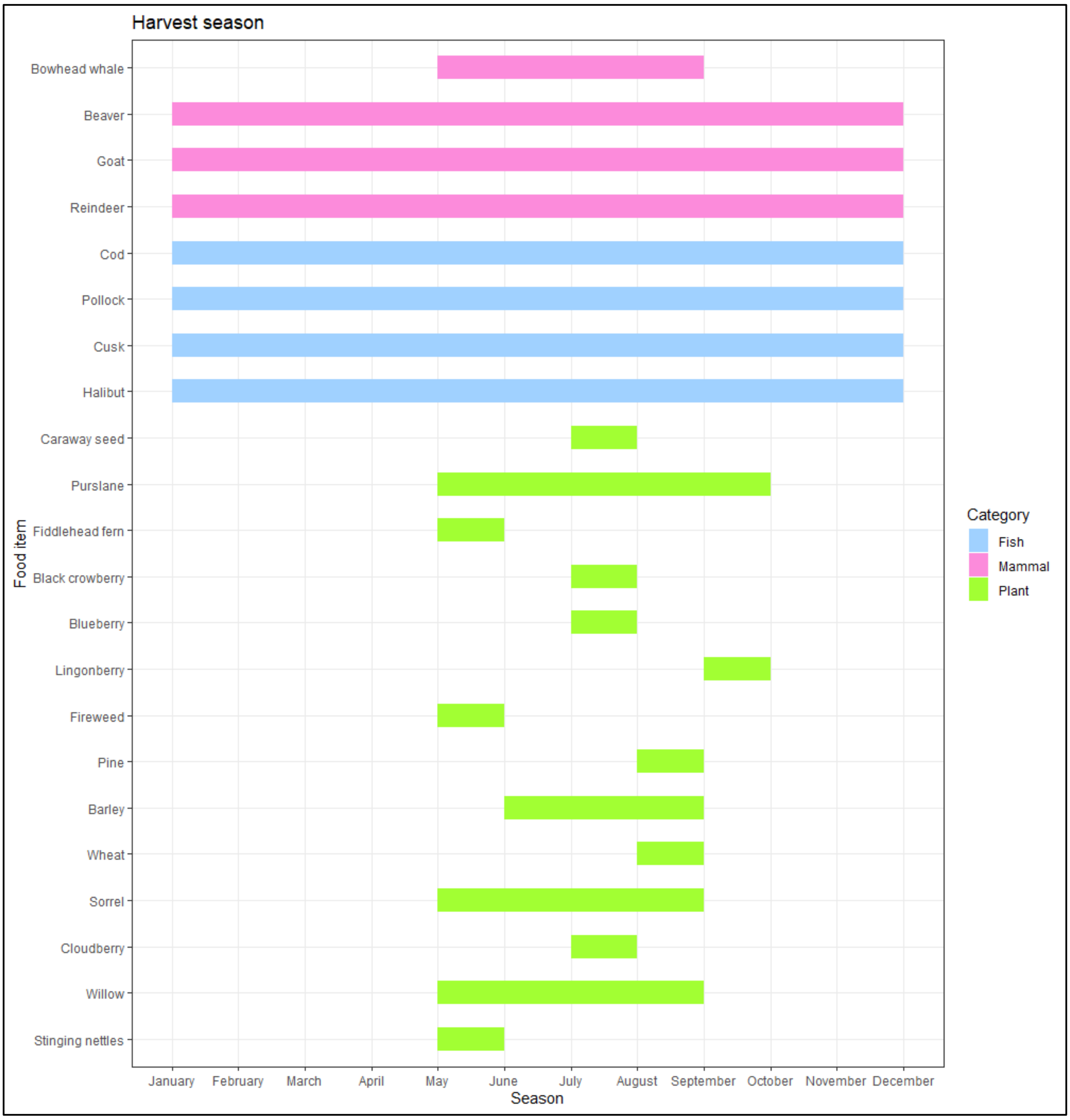

S37. List of taxa identified by sedaDNA removed from analyses.

* Sites: GB = Grassbaken (Baŋkgohppi), IG = Ingøy (Finnes).

** Negative controls: C001 = DNA extraction, C005 = DNA extract tube to plate, C006 = Ancient DNA lab, C007 = Post-PCR lab

| **Contaminants (those in negative controls as well as cultivars and domesticates that were not considered available at the time of the site occupancy).** | | | | |
|---|---|---|---|---|
| **Taxa removed** | **Category** | **Sites*** | **Present in negative controls**** | **Notes** |
| Anacardiaceae | Plant | GB, IG | No | Cashew; contamination. |
| Brassicaceae | Plant | GB | No | Suspected food contaminant as it does not match any species in PhyloNorway, PhyloAlps or ArcBorBryo. |
| *Ficus carica* | Plant | GB | No | Fig; contamination. |
| *Hordeum vulgare* | Plant | GB | | Cultivar. |
| Indigoferoid/millettioid clade | Plant | GB | No | Tribe of Fabaceae; assumed food contaminant. Above the species level. |
| *Larix* | Plant | GB | No | Found in single repeats at some northern Norwegian lakes in layers of 13,100-10,500 cal. BP and may represent contamination. |
| Musaceae | Plant | IG | No | Food contaminant. |
| *Persea* | Plant | GB, IG | No | Avocado; food contaminant. |
| *Picea abies* | Plant | GB, IG | C001 | GB: present in samples but not in negative controls. |
| Solanoideae | Plant | GB, IG | C005 | |

| | | | | |
|---|---|---|---|---|
| Solanoideae/*Solanum lycopersicum* | Plant | GB | C001 | Tomato; contamination. |
| *Tripleurospermum maritimum* | Plant | GB, IG | No | Assumed error in PhyloNorway as it matches Asteraceae in the three other reference libraries. Also, edible rating not available. |
| *Triticum* | Plant | GB, IG | No | Possible sequence error - only1 bp difference and also only in 1 sample at IG and 2 samples at GB. |
| *Triticum aestivum* | Plant | GB, IG | | Cultivar. |
| *Bos* | Mammal | GB, IG | C007 | |
| *Homo sapiens* | Mammal | GB, IG | C001, C005, C007 | |
| *Ovis* | Mammal | GB, IG | C007 | |
| *Dendrodrillus rubidus* | Annelid | IG | C001 | |
| **Higher taxonomic level/species not determined** (These taxa were not in negative controls) | | | | |
| **Taxa removed** | **Category** | **Sites*** | **Notes** | |
| Apioideae | Plant | GB | Higher taxonomic level. | |
| Aspleniineae (*Dryopteris crassirhizoma*) | Plant | GB | Higher taxonomic level. Dryopteris crassirhizoma not in Norway although BLAST 100% match to Dryopteris crassirhizoma. | |
| Asteraceae (*Hieracium, Pilosella, Taraxacum, Saussurea alpina*) | Plant | GB, IG | Species cannot be determined. | |
| Asteraceae (*Petasites, Taraxacum, Cicerbita*) | Plant | GB, IG | Species cannot be determined. | |
| Asteraceae/Asteroideae/Seneci oninae | Plant | IG | Species cannot be determined. | |
| Chlorellaceae | Plant | GB | Above the species level. | |
| Cichorioideae | Plant | IG | Species cannot be determined. | |
| Draba | Plant | IG | Species cannot be determined. Also, low edible rating. | |
| Lamioideae | Plant | GB | Above the species level and not in the region. | |
| Pooideae | Plant | GB, IG | Above the species level. | |

| | | | |
|---|---|---|---|
| *Salix/Populus tremula* | Plant | GB | Both *Salix* and *Populus tremula* also identified by itself. |
| **Taxa not in the region**<br>(These taxa were not in negative controls) | | | |
| **Taxa removed** | **Category** | **Sites*** | **Notes** |
| Humiriaceae (*Humiria balsamifera/Vantanea guianensis*) | Plant | GB | Neither Humiria balsamifera nor Vantanea guianensis in Norway. |
| Onagreae | Plant | GB | BLAST 100% match to Oenothera/Gaura/Camissonia/Clarkia species. These are not in the region. |
| *Solidago virgaurea* | Plants | GB, IG | |
| **Edible rating not available/Not edible (Edible rating below 3)**<br>(These taxa were not in negative controls) | | | |
| **Taxa removed** | **Category** | **Sites*** | **Notes** |
| *Agrostis/Calamagrostis* | Plant | GB, IG | Species with edible plant rating available that are in northern Norway are Agrostis capillaris only and low rating. |
| *Alnus incana* | Plant | GB, IG | Low edible rating. |
| *Alopecurus/Phleum* | Plant | GB | Only Alopecurus aequalis and Phleum pratense available in the edible plant database and low rating. |
| *Anthriscus sylvestris* | Plant | GB, IG | Used as a dye plant. |
| *Caltha palustris* | Plant | IG | Rating low. |
| *Campanula* cf. *rotundifolia* | Plant | GB | Rating low. |
| *Catabrosa aquatica/Poa annua* | Plant | GB, IG | Rating low. |
| *Ceratodon purpureus/ Rhabdoweisia crispata* | Plant | GB | Bryophyte excluded. |
| *Cornus suecica* | Plant | GB | Rating low. |
| *Deschampsia cespitosa/D. alpina* | Plant | GB | Edible rating available only for *Deschampsia cespitosa* and low. |
| *Dicranum spurium/D. scoparium* | Plant | GB, IG | Bryophyte excluded. |
| *Dryas octopetala* | Plant | GB | Rating low. |
| *Erysimum cheiranthoides/E. hieraciifolium* | Plant | GB | Rating low. |
| *Festuca/Lolium* | Plant | GB, IG | Lolium multiflorum native to the area but low edible rating. |
| *Geranium sylvaticum* | Plant | GB, IG | Rating low. |
| *Linnaea borealis* | Plant | GB | Rating low. |
| *Melampyrum* cf. *pratense* | Plant | GB | Edible rating not available. |
| *Myosotis arvensis* | Plant | GB | Edible rating not available. |
| *Phalaris arundinacea* | Plant | GB | Rating low. |

| | | | |
|---|---|---|---|
| *Poa pratensis/P. alpina/Anthoxanthum* | Plant | GB, IG | Rating low. |
| *Populus tremula* | Plant | GB | Rating low. |
| *Ptychostomum/ Rosulabryum/Bryum* | Plant | GB | Bryophyte excluded. |
| *Ranunculus acris/R. subborealis* | Plant | IG | Species cannot be determined as well as low edible rates. |
| *Ranunculus acris/R. subborealis/R. repens* | Plant | IG | Species cannot be determined as well as low edible rates. |
| *Ranunculus repens* | Plant | IG | Rating low. |
| *Rhinanthus alectorolophus* | Plant | GB, IG | Edible rating not available. |
| *Rhinanthus minor* | Plant | GB, IG | Used in children's games, but not much more. Low edible rates. |
| *Saussurea alpina/Lactuca sibirica* | Plant | IG | Edible rating available only for Lactuca sibirica but low. |
| *Scorzoneroides autumnalis* | Plant | IG | Edible rating not available. |
| *Silene dioica* | Plant | GB, IG | Rating low. |
| *Sphagnum fimbriatum* | Plant | IG | Bryophyte excluded. |
| *Trientalis europaea* | Plant | GB, IG | Rating low. |
| *Tripleurospermum maritimum* | Plant | GB | Edible rating not available. |
| Triticeae (*Elymus/Leymus*) | Plant | GB, IG | Edible rating available only for *Leymus arenaria* and low. |
| *Veronica longifolia* | Plant | GB | Rating low. |
| *Vicia cracca/V. sepium* | Plant | GB, IG | Rating low. |
| *Zostera marina* | Plant | GB | Rating low. |

S38. List of taxa identified by macrofossils that are removed from nutritional analysis.

| **Higher taxonomic level** | | | |
|---|---|---|---|
| **Taxa removed** | **Category** | **Sites *** | **Note** |
| Apiaceae | Plant | GB | Family level. |
| Asteroideae | Plant | GB | |
| Caryophyllaceae | Plant | GB | Family level. |
| Cetacea | Mammal | GB | |
| Chamerion | Plant | GB | |
| Cichorioideae | Plant | GB | |
| Cyperaceae | Plant | GB | |
| Ericales | Plant | GB | |
| Fabaceae | Plant | GB | |
| Picea | Plant | GB | |
| Poaceae | Plant | GB | |
| Rosaceae | Plant | GB | |
| **Macro/Microfossils considered not edible/low edible rating** | | | |
| **Taxa removed** | **Category** | **Sites *** | **Note** |
| *Carex* | Plant | GB, IG | Edible rating low (inner stem can be chewed but not considered important food). |
| *Drosera* | Plant | GB | Edible rating not available. |
| *Picea* | Plant | GB | Low edible rating. |
| *Ranunculus* | Plant | GB | Low edible rating. |
| *Saussurea alpina* | Plant | GB | Edible rating not available. |
| *Selaginella selaginoides* | Plant | GB | Edible rating not available. |
| *Sphagnum* | Plant | GB | Low edible rating. |

S39. List of edible taxa identified by sedaDNA and macrofossils removed from nutritional analysis due to lack of nutritional information.

| Taxa removed due to nutritional information not available | | | |
|---|---|---|---|
| **Taxa removed** | **Category** | **Sites *** | **Identified by** |
| *Achillea millefolium* | Plant | IG | sedaDNA |
| *Alchemilla alpina* | Plant | IG | sedaDNA |
| *Alnus* | Plant | GB | Macrofossil |
| Apiaceae | Plant | GB | Macrofossil |
| *Arnica angustifolia* | Plant | GB, IG | sedaDNA |
| *Artemisia* | Plant | GB | Macrofossil |
| Asteroideae | Plant | GB | Macrofossil |
| *Atriplex glabriuscula* | Plant | GB, IG | sedaDNA |
| *Avenella flexuosa* | Plant | GB, IG | sedaDNA |
| *Betula* | Plant | GB | Macrofossil |
| *Betula pendula* | Plant | GB, IG | sedaDNA |
| *Betula pubescens* | Plant | GB, IG | sedaDNA |
| *Bistorta vivipara* | Plant | IG | sedaDNA |
| *Carex* | Plant | GB | Macrofossil |
| Caryophyllaceae | Plant | GB | Macrofossil |
| *Chamerion* | Plant | GB | Macrofossil |
| Cichorioideae | Plant | GB | Macrofossil |
| *Cochlearia officinalis* | Plant | GB, IG | sedaDNA |
| Cyperaceae | Plant | GB | Macrofossil |
| *Drosera* | Plant | GB | Macrofossil |
| *Dryopteris crassirhizoma* | Plant | GB | sedaDNA |
| *Dryopteris filix−mas* | Plant | GB, IG | sedaDNA |
| *Equisetum arvense* | Plant | GB, IG | sedaDNA |
| *Equisetum hyemale* | Plant | GB, IG | sedaDNA |
| *Equisetum pratense* | Plant | GB | sedaDNA |
| Ericales | Plant | GB | Macrofossil |
| *Euphrasia officinalis* | Plant | GB, IG | sedaDNA |
| Fabaceae | Plant | GB | Macrofossil |
| *Filipendula* | Plant | GB | Macrofossil |
| *Filipendula ulmaria* | Plant | GB, IG | sedaDNA |

| | | | |
|---|---|---|---|
| *Geum rivale* | Plant | IG | sedaDNA |
| *Glechoma hederacea* | Plant | IG | sedaDNA |
| *Juniperus* | Plant | GB | Macrofossil |
| *Juniperus communis* | Plant | GB, IG | sedaDNA |
| *Larix* | Plant | GB | Macrofossil |
| *Leymus arenarius* | Plant | GB, IG | sedaDNA |
| *Ligusticum scoticum* | Plant | GB | sedaDNA |
| *Picea* | Plant | GB | Macrofossil |
| *Pinus* | Plant | GB | Macrofossil |
| Poaceae | Plant | GB | Macrofossil |
| *Polypodium* | Plant | GB | Macrofossil |
| *Populus tremula* | Plant | GB, IG | sedaDNA |
| *Potentilla anserina* | Plant | IG | sedaDNA |
| *Ranunculus* | Plant | GB | Macrofossil |
| Rosaceae | Plant | GB | Macrofossil |
| *Rumex* | Plant | GB | Macrofossil |
| *Saussurea alpina* | Plant | GB | Macrofossil |
| *Selaginella selaginoides* | Plant | GB | Macrofossil |
| *Silybum marianum* | Plant | GB | sedaDNA |
| *Sorbus aucuparia* | Plant | GB, IG | sedaDNA |
| *Sphagnum* | Plant | GB | Macrofossil |
| *Valeriana sambucifolia* | Plant | GB, IG | sedaDNA |
| *Canis* | Mammal | GB | sedaDNA |
| *Canis lupus* | Mammal | GB | sedaDNA |
| Cetacea | Mammal | GB | Macrofossil |
| *Halichoerus grypus* | Mammal | GB, IG | Macrofossil |
| *Lutra lutra* | Mammal | IG | Macrofossil |
| *Myodes glareolus* | Mammal | GB | sedaDNA |
| *Phoca groenlandica* | Mammal | GB | Macrofossil, sedaDNA |
| *Phoca vitulina* | Mammal | GB, IG | Macrofossil |
| Phocidae | Mammal | GB, IG | Macrofossil |
| Rodentia | Mammal | GB | Macrofossil |
| *Arctogadus glacialis* | Fish | IG | sedaDNA |
| Gadidae | Fish | IG | Macrofossil |
| *Hippoglossus** | Fish | IG | sedaDNA |
| *Pollachius** | Fish | IG | Macrofossil |
| *Arctica islandica* | Shellfish | GB | Macrofossil |
| *Echinus* | Echinoderm | GB | Macrofossil |

| | | | |
|---|---|---|---|
| *Littorina* | Shellfish | GB, IG | GB: Macrofossil, sedaDNA<br>IG: sedaDNA |
| *Littorina littorea* | Shellfish | GB | Macrofossil |
| *Strongylocentrotus droebachiensis* | Echinoderm | GB | sedaDNA |
| Alcidae | Bird | IG | Macrofossil |
| *Cepphus grylle* | Bird | IG | Macrofossil |
| *Corvus corax* | Bird | IG | Macrofossil |
| *Fratercula arctica* | Bird | GB | Macrofossil |
| *Fulmarus glacialis* | Bird | IG | Macrofossil |
| *Gallinago gallinago* | Bird | GB | Macrofossil |
| Laridae | Bird | IG | Macrofossil |
| *Larus* | Bird | IG | Macrofossil |
| *Larus argentatus* | Bird | GB | Macrofossil |
| *Larus marinus* | Bird | IG | Macrofossil |
| *Mergus merganser* | Bird | IG | Macrofossil |
| *Mergus serrator* | Bird | IG | Macrofossil |
| *Morus bassanus* | Bird | GB, IG | Macrofossil |
| Phalacrocorax | Bird | IG | Macrofossil |
| *Phalacrocorax aristotelis* | Bird | IG | Macrofossil |
| Phasianidae | Bird | GB | Macrofossil |
| *Pinguinus impennis* | Bird | GB | Macrofossil |
| *Rissa tridactyla* | Bird | GB, IG | Macrofossil |
| *Uria aalge* | Bird | GB | Macrofossil, sedaDNA |
| *Auxenochlorella protothecoides* | Algae | GB | sedaDNA |
| *Bracteacoccus bullatus* | Algae | GB | sedaDNA |
| *Ulvaria fusca* | Algae | GB | sedaDNA |
| *Dendrodrilus rubidus* | Annelid | IG | sedaDNA |
| *Enchytraeus albidus* | Annelid | GB, IG | sedaDNA |
| *Fridericia parathalassia* | Annelid | IG | sedaDNA |
| *Henlea perpusilla* | Annelid | GB | sedaDNA |
| *Lumbricus rubellus* | Annelid | IG | sedaDNA |
| Amphibia | Amphibian | IG | Macrofossil |

* Included at the species level.

S40. List of recommended daily allowance (RDA) in the R package NutrienTrackeR (Health Canada 2015; Rodriguez-Martinez et al. 2023)

| Gender/Pregnancy/Lactation, Age | Vitamins | | | | | | | | | | | | | |
|---|---|---|---|---|---|---|---|---|---|---|---|---|---|---|
| | A (ug) | C (mg) | D (ug) | E (mg) | K (ug) | B1 (mg) | B2 (mg) | B3 (mg) | B6 (mg) | B9 (ug) | B12 (ug) | B5 (mg) | B7 (ug) | B4 (mg) |
| Infant 0-0.5 | 400 | 40 | 10 | 4 | 2.0 | 0.2 | 0.3 | 2 | 0.1 | 65 | 0.4 | 1.7 | 5 | 125 |
| Infant 0.6-0.9 | 500 | 50 | 10 | 5 | 2.5 | 0.3 | 0.4 | 4 | 0.3 | 80 | 0.5 | 1.8 | 6 | 150 |
| Children 1-3 | 300 | 15 | 15 | 6 | 30.0 | 0.5 | 0.5 | 6 | 0.5 | 150 | 0.9 | 2.0 | 8 | 200 |
| Children 4-8 | 400 | 25 | 15 | 7 | 55.0 | 0.6 | 0.6 | 8 | 0.6 | 200 | 1.2 | 3.0 | 12 | 250 |
| Male 9-13 | 600 | 45 | 15 | 11 | 60.0 | 0.9 | 0.9 | 12 | 1.0 | 300 | 1.8 | 4.0 | 20 | 375 |
| Male 14-18 | 900 | 75 | 15 | 15 | 75.0 | 1.2 | 1.3 | 16 | 1.3 | 400 | 2.4 | 5.0 | 25 | 550 |
| Male 19-30 | 900 | 90 | 15 | 15 | 120.0 | 1.2 | 1.3 | 16 | 1.3 | 400 | 2.4 | 5.0 | 30 | 550 |
| Male 31-50 | 900 | 90 | 15 | 15 | 120.0 | 1.2 | 1.3 | 16 | 1.3 | 400 | 2.4 | 5.0 | 30 | 550 |
| Male 51-70 | 900 | 90 | 15 | 15 | 120.0 | 1.2 | 1.3 | 16 | 1.7 | 400 | 2.4 | 5.0 | 30 | 550 |
| Male >70 | 900 | 90 | 20 | 15 | 120.0 | 1.2 | 1.3 | 16 | 1.7 | 400 | 2.4 | 5.0 | 30 | 550 |
| Female 9-13 | 600 | 45 | 15 | 11 | 60.0 | 0.9 | 0.9 | 12 | 1.0 | 300 | 1.8 | 4.0 | 20 | 375 |
| Female 14-18 | 700 | 65 | 15 | 15 | 75.0 | 1.0 | 1.0 | 14 | 1.2 | 400 | 2.4 | 5.0 | 25 | 400 |
| Female 19-30 | 700 | 75 | 15 | 15 | 90.0 | 1.1 | 1.1 | 14 | 1.3 | 400 | 2.4 | 5.0 | 30 | 425 |
| Female 31-50 | 700 | 75 | 15 | 15 | 90.0 | 1.1 | 1.1 | 14 | 1.3 | 400 | 2.4 | 5.0 | 30 | 425 |
| Female 51-70 | 700 | 75 | 15 | 15 | 90.0 | 1.1 | 1.1 | 14 | 1.5 | 400 | 2.4 | 5.0 | 30 | 425 |
| Female >70 | 700 | 75 | 20 | 15 | 90.0 | 1.1 | 1.1 | 14 | 1.5 | 400 | 2.4 | 5.0 | 30 | 425 |
| Pregnant 14-18 | 750 | 80 | 15 | 15 | 75.0 | 1.4 | 1.4 | 18 | 1.9 | 600 | 2.6 | 6.0 | 30 | 450 |
| Pregnant 19-30 | 770 | 85 | 15 | 15 | 90.0 | 1.4 | 1.4 | 18 | 1.9 | 600 | 2.6 | 6.0 | 30 | 450 |
| Pregnant 31-50 | 770 | 85 | 15 | 15 | 90.0 | 1.4 | 1.4 | 18 | 1.9 | 600 | 2.6 | 6.0 | 30 | 450 |
| Lactation 14-18 | 1200 | 115 | 15 | 19 | 75.0 | 1.4 | 1.6 | 17 | 2.0 | 500 | 2.8 | 7.0 | 35 | 550 |
| Lactation 19-30 | 1300 | 120 | 15 | 19 | 90.0 | 1.4 | 1.6 | 17 | 2.0 | 500 | 2.8 | 7.0 | 35 | 550 |
| Lactation 31-50 | 1300 | 120 | 15 | 19 | 90.0 | 1.4 | 1.6 | 17 | 2.0 | 500 | 2.8 | 7.0 | 35 | 550 |

S40. Cont.

| Gender/Pregnancy/Lactation, Age | Minerals | | | | | | | | | | | | | | |
|---|---|---|---|---|---|---|---|---|---|---|---|---|---|---|---|
| | Ca (mg) | Cr (ug) | Cu (mg) | F (ug) | I (ug) | Fe (mg) | Mg (mg) | Mn (mg) | Mo (ug) | P (mg) | Se (ug) | Zn (mg) | K (mg) | Na (mg) | $Cl^-$ (g) |
| Infant 0-0.5 | 200 | 0.2 | 0.20 | 10 | 110 | 0.27 | 30 | 0.0 | 2 | 100 | 15 | 2 | 400 | 120 | 0.18 |
| Infant 0.6-0.9 | 260 | 5.5 | 0.22 | 500 | 130 | 11.00 | 75 | 0.6 | 3 | 275 | 20 | 3 | 700 | 370 | 0.57 |
| Children 1-3 | 700 | 11.0 | 0.34 | 700 | 90 | 7.00 | 80 | 1.2 | 17 | 460 | 20 | 3 | 3000 | 1000 | 1.50 |
| Children 4-8 | 1000 | 15.0 | 0.44 | 1000 | 90 | 10.00 | 130 | 1.5 | 22 | 500 | 30 | 5 | 3800 | 1200 | 1.90 |
| Male 9-13 | 1300 | 25.0 | 0.70 | 2000 | 120 | 8.00 | 240 | 1.9 | 34 | 1250 | 40 | 8 | 4500 | 1500 | 2.30 |
| Male 14-18 | 1300 | 35.0 | 0.89 | 3000 | 150 | 11.0 | 410 | 2.2 | 43 | 1250 | 55 | 11 | 4700 | 1500 | 2.30 |
| Male 19-30 | 1000 | 35.0 | 0.90 | 4000 | 150 | 8.00 | 400 | 2.3 | 45 | 700 | 55 | 11 | 4700 | 1500 | 2.30 |
| Male 31-50 | 1000 | 35.0 | 0.90 | 4000 | 150 | 8.00 | 420 | 2.3 | 45 | 700 | 55 | 11 | 4700 | 1500 | 2.30 |
| Male 51-70 | 1000 | 30.0 | 0.90 | 4000 | 150 | 8.00 | 420 | 2.3 | 45 | 700 | 55 | 11 | 4700 | 1300 | 2.00 |
| Male >70 | 1200 | 30.0 | 0.90 | 4000 | 150 | 8.00 | 420 | 2.3 | 45 | 700 | 55 | 11 | 4700 | 1200 | 1.80 |
| Female 9-13 | 1300 | 21.0 | 0.70 | 2000 | 120 | 8.00 | 240 | 1.6 | 34 | 1250 | 40 | 8 | 4500 | 1500 | 2.30 |
| Female 14-18 | 1300 | 24.0 | 0.89 | 3000 | 150 | 15.00 | 360 | 1.6 | 43 | 1250 | 55 | 9 | 4700 | 1500 | 2.30 |
| Female 19-30 | 1000 | 25.0 | 0.90 | 3000 | 150 | 18.00 | 310 | 1.8 | 45 | 700 | 55 | 8 | 4700 | 1500 | 2.30 |
| Female 31-50 | 1000 | 25.0 | 0.90 | 3000 | 150 | 18.00 | 320 | 1.8 | 45 | 700 | 55 | 8 | 4700 | 1500 | 2.30 |
| Female 51-70 | 1200 | 20.0 | 0.90 | 3000 | 150 | 8.00 | 320 | 1.8 | 45 | 700 | 55 | 8 | 4700 | 1300 | 2.00 |
| Female >70 | 1200 | 20.0 | 0.90 | 3000 | 150 | 8.00 | 320 | 1.8 | 45 | 700 | 55 | 8 | 4700 | 1200 | 1.80 |
| Pregnant 14-18 | 1300 | 29.0 | 1.00 | 3000 | 220 | 27.00 | 400 | 2.0 | 50 | 1250 | 60 | 12 | 4700 | 1500 | 2.30 |
| Pregnant 19-30 | 1000 | 30.0 | 1.00 | 3000 | 220 | 27.00 | 350 | 2.0 | 50 | 700 | 60 | 11 | 4700 | 1500 | 2.30 |
| Pregnant 31-50 | 1000 | 30.0 | 1.00 | 3000 | 220 | 27.00 | 360 | 2.0 | 50 | 700 | 60 | 11 | 4700 | 1500 | 2.30 |
| Lactation 14-18 | 1300 | 44.0 | 1.30 | 3000 | 290 | 10.00 | 360 | 2.6 | 50 | 1250 | 70 | 13 | 5100 | 1500 | 2.30 |
| Lactation 19-30 | 1000 | 45.0 | 1.30 | 3000 | 290 | 9.00 | 310 | 2.6 | 50 | 700 | 70 | 12 | 5100 | 1500 | 2.30 |
| Lactation 31-50 | 1000 | 45.0 | 1.30 | 3000 | 290 | 9.00 | 320 | 2.6 | 50 | 700 | 70 | 12 | 5100 | 1500 | 2.30 |

S40. Cont.

| Gender/Pregnancy/Lactation, Age | Carbohydrate (g) | Fiber (g) | Protein (g) |
|---|---|---|---|
| Infant 0-0.5 | 60 | NA | 9.1 |
| Infant 0.6-0.9 | 95 | NA | 11.0 |
| Children 1-3 | 130 | 19 | 13.0 |
| Children 4-8 | 130 | 25 | 19.0 |
| Male 9-13 | 130 | 31 | 34.0 |
| Male 14-18 | 130 | 38 | 52.0 |
| Male 19-30 | 130 | 38 | 56.0 |
| Male 31-50 | 130 | 38 | 56.0 |
| Male 51-70 | 130 | 30 | 56.0 |
| Male >70 | 130 | 30 | 56.0 |
| Female 9-13 | 130 | 26 | 34.0 |
| Female 14-18 | 130 | 26 | 46.0 |
| Female 19-30 | 130 | 25 | 46.0 |
| Female 31-50 | 130 | 25 | 46.0 |
| Female 51-70 | 130 | 21 | 46.0 |
| Female >70 | 130 | 21 | 46.0 |
| Pregnant 14-18 | 175 | 28 | 71.0 |
| Pregnant 19-30 | 175 | 28 | 71.0 |
| Pregnant 31-50 | 175 | 28 | 71.0 |
| Lactation 14-18 | 210 | 20 | 71.0 |
| Lactation 19-30 | 210 | 29 | 71.0 |
| Lactation 31-50 | 210 | 29 | 71.0 |

Martens, Vibeke Vandrup, Elin Rose Myrvoll, Ove Bergersen, Michel Vorenhout, Paula Utigard Sandvik, and Sean Denham. 2017. "InSituFarms. Archaeological Deposits in a Changing Climate: Case Baŋkgohppi, Unjárgga gielda/Nesseby k., Finnmark." 77. NIKU Rapport. Oslo: Norsk Institutt for Kulturminneforskning.

Reimer, Paula J., William E. N. Austin, Edouard Bard, Alex Bayliss, Paul G. Blackwell, Christopher Bronk Ramsey, Martin Butzin, et al. 2020. "The IntCal20 Northern Hemisphere Radiocarbon Age Calibration Curve (0–55 Cal kBP)." *Radiocarbon* 62 (4): 725–57. https://doi.org/10.1017/RDC.2020.41.

Rodriguez-Martinez, Andrea, Rafael Ayala, Mark Balchunas, Daniel Gutierrez-Del-Rio, Ting-Hua Chen, Yacine Debbabi, and Lara Selles Vidal. 2023. "NutrienTrackeR: Food Composition Information and Dietary Assessment." https://CRAN.R-project.org/package=NutrienTrackeR.

Wickler, Stephen. 2016. "The Centrality of Small Islands in Arctic Norway From the Viking Age to Recent Historic Period." *The Journal of Island and Coastal Archaeology* 11 (2): 171–94. https://doi.org/10.1080/15564894.2015.1134728.

Wickler, Stephen. 2026. "Maritime Transformations in Northern Norway during the Viking Age and Medieval Period: Innovation and Independence on the Edge of Europe." In *Harvesting the Margins of the Viking World: Decentralizing Economies in Late Iron Age Fennoscandia*, edited by Charlotta Hillerdal and Kristin Ilves, 6:79–98. The Archaeology of Northern Europe. Turnhout, Belgium: Brepols Publishers.
